## Supplementary Information for "Active Learning-Assisted Directed Evolution"

#### Contents


| | |
|---|---|
| General Information | S1 |
| Cloning and Sequence Design | S3 |
| Cloning Protocols and Results | S8 |
| Preparation of Authentic Standards | S18 |
| Preparation of Calibration Curves for Analytical Yield Determination | S26 |
| Control and Validation Experiments | S36 |
| Biocatalytic Cyclopropanation of Olefins | S40 |
| Chiral Traces for Determination of Enantiopurity | S73 |
| $^{1}H$ NMR Spectra of Authentic Standards | S75 |
| References | S86 |

### General Information

**Safety Statement:**

All chemical transformations were performed in a well-ventilated fume hood to avoid inhalation and exposure. Other than that, no unexpected or unusually high safety concerns were raised with these methods. Safety notes for individual synthetic procedures will be documented alongside the procedure.

**General Information:**

All chemical transformations were performed in a well-ventilated fume hood to avoid inhalation and exposure to chemicals. Reagents and solvents were obtained commercially (Sigma-Aldrich, Alfa Aesar, VWR, Fisher Scientific, Matrix Scientific, Oakwood Chemical, TCI America, and other suppliers) and used without prior purification unless otherwise stated. Organic solutions were concentrated under reduced pressure on an IKA RV 10 rotary evaporator. Thin-layer chromatography (TLC) was performed on commercial Millipore Silica Gel 60 plates containing the F254 fluorescent indicator. Visualization of the developed chromatographs was performed by irradiation with UV light or by treatment with an appropriate TLC staining solution (e.g., Ceric Ammonium Molybdate, $KMnO_4$, or Bromocresol Green) followed by heating if necessary. Chromatographic purification was accomplished by flash chromatography using a Biotage Isolera One instrument.

**Spectral Data:**

All NMR spectra were obtained at the Caltech Liquid NMR Facility. For all cyclopropane compounds, $^1$H NMR were recorded on a Bruker Prodigy 400 MHz instrument (400 MHz and 101

MHz). For intermediates, $^1H$ spectra were also recorded using a Varian 300 MHz spectrometer (300 MHz), a Varian 500 MHz spectrometer (500 MHz), and a Varian 600 MHz spectrometer (600 MHz). $^1H$ spectra are referred to residual $CDCl_3$ solvent signals referenced at δ 7.26 ppm. Data for $^1H$ NMR are reported as follows: chemical shift (δ ppm), integration, multiplicity (s = singlet, d = doublet, t = triplet, q = quartet, p = pentad, sext = sextet, hept = heptet, m = multiplet, br s = broad singlet), and coupling constant (Hz).

**Gas Chromatography Data:**

GC chromatography (GC) was performed on an Agilent Technologies 7820A GC system equipped with a split-mode capillary injection system and flame-ionization detector. For achiral analyses, an Agilent J&W HP-5 Column was used as the stationary phase. For chiral analyses, the specific stationary phase is provided along with the chiral traces.

### Cloning and Sequence Design

#### Relevant Protoglobin Sequences:

**Table S1.** The amino acid sequences of wild-type *Par*Pgb and the previously engineered variant, ParLQ. Residues in each sequence highlighted in yellow are the sites which were investigated in this work for wet-lab experimentation: W56, Y57, L59, Q60, and F89 in ParLQ

| Protein Variant | Amino Acid Sequence |
|---|---|
| wild-type *Pyrobaculum arsenenticum* Protoglobin (*Par*Pgb) | MAVPGYDFGKVPDAPISDADFESLKKTVM WGEEDEKYRKMACEALKGQVEDILDLWY GWVGSNQHLIYYFGDKSGRPIPQYLEAVRK RFGLWIIDTLCKPLDRQWLNYMYEIGLRHH RTKKGKTDGVDTVEHIPLRYMIAFIAPIGLTI KPILEKSGHPPEAVERMWAAWVKLVVLQV AIWSYPYAKTGEW |
| *Pyrobaculum arsenenticum* Protoglobin W59L V60Q (ParLQ) | MAVPGYDFGKVPDAPISDADFESLKKTVM WGEEDEKYRKMACEALKGQVEDILDLWY GLQGSNQHLIYYFGDKSGRPIPQYLEAVRK RFGLWIIDTLCKPLDRQWLNYMYEIGLRHH RTKKGKTDGVDTVEHIPLRYMIAFIAPIGLTI KPILEKSGHPPEAVERMWAAWVKLVVLQV AIWSYPYAKTGEW |

.

#### DNA Sequence of ParLQ

ATGGCGGTTCCCGGCTACGATTTTGGCAAAGTCCCGGATGCCCCAATCTCAGACGCG
GATTTTGAGAGTTTAAAAAAAACCGTGATGTGGGGTGAGGAAGATGAGAAATATCG
CAAAATGGCTTGCGAAGCCTTAAAGGGTCAAGTAGAAGATATTTTAGATTTGTGGTA
CGGCCTGCAGGGAAGCAATCAACACCTTATCTACTACTTCGGTGATAAGAGTGGTCG
TCCAATTCCGCAATACCTGGAAGCGGTCCGCAAGCGTTTCGGGTTGTGGATCATTGA
TACATTGTGTAAGCCACTGGACCGCCAGTGGTTGAATTACATGTACGAAATTGGCCT
TCGCCATCACCGTACCAAGAAAGGGAAGACAGATGGCGTAGATACTGTTGAACATA
TCCCATTACGCTACATGATTGCTTTCATCGCTCCCATCGGTCTGACTATTAAGCCGAT
CTTGGAAAAATCGGGACATCCGCCAGAGGCCGTGGAGCGTATGTGGGCAGCATGGG
TTAAGTTGGTGGTGTTACAGGTAGCTATCTGGTCGTACCCCTATGCAAAGACGGGCG
AATGGCTCGAGCACCACCACCACCACCAC

**Table S2.** Codons utilized when ALDE-predicted mutations were incorporated into the ParLQ DNA sequence. Codons which are generally recognized as the most prevalently found in the *E. coli* genome were selected.

| Amino Acid | Codon |
|---|---|
| A | GCC |
| C | TGC |
| D | GAT |
| E | GAA |
| F | TTT |
| G | GGC |
| H | CAT |
| I | ATT |
| K | AAA |
| L | TTA |
| M | ATG |
| N | AAC |
| P | CCG |
| Q | CAG |
| R | CGT |
| S | AGC |
| T | ACC |
| V | GTG |
| W | TGG |
| Y | TAT |

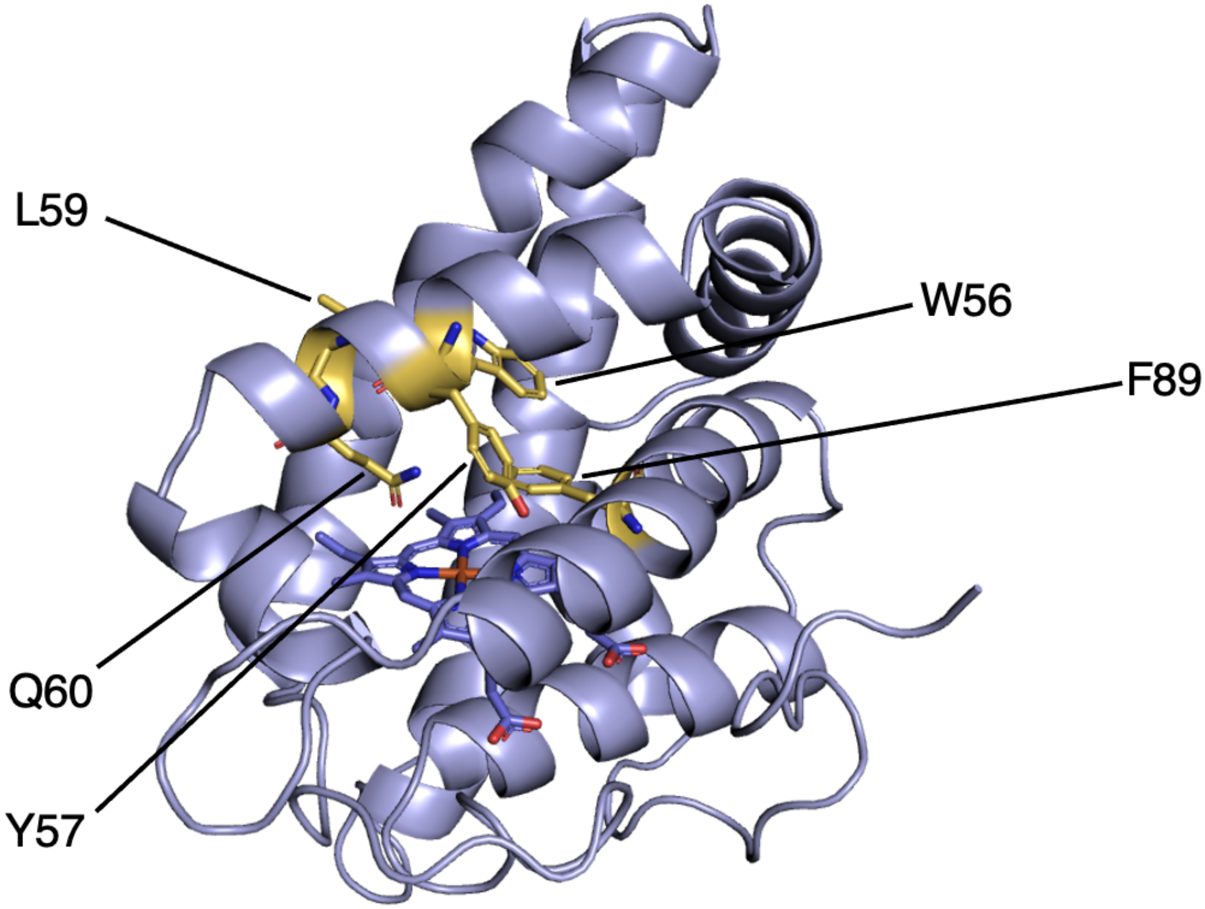


**Figure S1.** Homology model of ParLQ with docked heme. Structural modeling was performed with AlphaFold3.[1] Residues which were mutated in this study are illustrated in yellow.

**Nomenclature for Variant Naming**

Single-site mutants are named using the standard nomenclature: (original AA)(site)(new AA). The mutant W56F refers to a variant of ParLQ mutated from tryptophan to phenylalanine. Five-site multi-mutants are named as a string of the five amino acids to which the positions of interest have been mutated. The variant MPFDY refers to a variant of ParLQ bearing the mutations W56M, Y57P, L59F, Q60D, and F89Y. In this nomenclature, ParLQ is named WLYQF.

#### Primer Design for Site-Saturation Mutagenesis (SSM):

*General Cloning Primers*

**Table S3.** Primers 007 and primers 008 are used to generate amplicons of linearized pET-22b(+)[2] backbone. Primers 005 and 006 were used with primers internal to the protoglobin gene (**Table S4**) to generate mutant protoglobin genes. All primers were ordered from IDT (Coralville, IA).

| Primer Name | Direction | Sequence | Description |
|---|---|---|---|
| 005 | Forward | 5'-GAAATAATTTTGTTTAACTTTAAGAAGGAGATATACATATG-3' | Upstream of N-term, anneals with 007 |
| 006 | Reverse | 5'-GCCGGATCTCAGTGGTGGTGGTGGTGGTGCTCGAG-3' | Downstream of C-term, anneals with 008 |
| 007 | Reverse | 5'-CATATGTATATCTCCTTCTTAAAGTTAAACAAAATTATTTC-3' | Upstream of N-term, anneals with 005 |
| 008 | Forward | 5'-CTCGAGCACCACCACCACCACCACTGAGATCCGGC-3' | Downstream of C-term, anneals with 006 |

*Cloning Primers for Site-Saturation Mutagenesis*

**Table S4.** Primers containing degenerate codons at sites of interest for SSM. Primers were used in conjunction with either primer 005 or 006 (**Table S3**) to generate mutant protoglobin genes. All primers were ordered from IDT (Coralville, IA).

| **Primer Name** | **Direction** | **Sequence** | **Description** |
|---|---|---|---|
| ParLQ_56X_F | Forward | 5'-GGTCAAGTAGAAGATATTTTAGATTTGNNKTACGGCCTGCAGGGAAGCAATC-3' | SSM for site 56 |
| ParLQ_56X_R | Reverse | 5'-CAAATCTAAAATATCTTCTACTTGACCCTTTAAGGCTTC-3' | SSM for site 56 |
| ParLQ_57X_F | Forward | 5'-CAAGTAGAAGATATTTTAGATTTGTGGNNKGGCCTGCAGGGAAGCAATCAAC-3' | SSM for site 57 |
| ParLQ_57X_R | Reverse | 5'-CCACAAATCTAAAATATCTTCTACTTGACCCTTTAAGGC-3' | SSM for site 57 |
| ParLQ_59X_F | Forward | 5'-TATTTTAGATTTGTGGTACGGCNNKCAGGGAAGCAATCAACACCTTATCTACTAC-3' | SSM for site 59 |
| ParLQ_59X_R | Reverse | 5'-GCCGTACCACAAATCTAAAATATCTTCTACTTGACCC-3' | SSM for site 59 |
| ParLQ_60X_F | Forward | 5'-TTTGTGGTACGGCCTGNNKGGAAGCAATCAACACCTTATCTACTACTTCGG-3' | SSM for site 60 |
| ParLQ_60X_R | Reverse | 5'-CAGGCCGTACCACAAATCTAAAATATCTTCTACTTG-3' | SM for site 60 |
| ParLQ_89X_F | Forward | 5'-GCGGTCCGCAAGCGTNNKGGGTTGTGGATC-3' | SSM for site 89 |
| ParLQ_89X_R | Reverse | 5'-CCMNNACGCTTGCGGACCGCTTCCAGG-3' | SSM for site 89 |
| ParLQ_quadNNK_F | Forward | 5'-CAAGTAGAAGATATTTTAGATTTG NNKNNKGGCNNKNNKGGAAGCAATCAACACCTTATC-3' | Multisite SSM for sites 56, 57, 59, and 60 |
| ParLQ_quadNNK_R | Reverse | 5'-GATAAGGTGTTGATTGCTTCCMNNMNNGCCMNNMNNCAAATCTAAAATATCTTCTACTTG-3' | Multisite SSM for sites 56, 57, 59, and 60 |

### Cloning Protocols and Results

*Protocols for the Cloning of Random ParLQ Variants*

**Cloning for Single Site-Saturation Mutagenesis (SSM):**

Chemically competent *Escherichia coli* (*E. coli*) cells (T7 Express Competent *E. coli*) were purchased from New England Biolabs (NEB, Ipswich, MA). Additionally, Phusion polymerase and *Dpn*I were purchased from NEB. SSM experiments were performed using primers bearing degenerate codons (NNK) using a modified QuikChange™ protocol (**Table S4**).[3] The following PCR amplicons were generated using a parent plasmid (pET-22b(+) harboring ParLQ):

**Table S5.** Primer combinations for the generation of PCR amplicons for the construction of expression plasmids containing mutagenized ParLQ variants. '*site*' refers to the specific site which is being saturated.

| Fragment Name | Forward Primer | Reverse Primer |
|---|---|---|
| SSM_Frag1 | 005 | ParLQ_*site*_R |
| SSM_Frag2 | ParLQ_*site*_F | 006 |
| pET_Backbone | 008 | 007 |

The PCR conditions were as follows (final concentrations): Phusion HF Buffer 1x, 0.2 mM dNTPs each, 0.5 µM of forward primers, 0.5 µM reverse primer, and 0.02 U/µL of Phusion polymerase. The standard Phusion PCR protocol was used.[4] Upon completion of PCRs, the remaining template was digested with *Dpn*I. Gel purification was performed with a Zymoclean Gel DNA Recovery Kit (Zymo Research Corp, Irvine, CA). The purified PCR product was then assembled using the Gibson assembly protocol.[5]

**Transformation of *E. coli* with the Genes Coding for Single-Site Mutants:**

96-Well deep-well plates were shaken in an INFORS HT Multitron Shaker in all instances. The assembly products obtained were used to transform T7 Express Competent *E. coli* (High Efficency) cells (NEB, Ipswich, MA) following the protocol recommended by the manufacturer. Upon heat-shock, freshly transformed *E. coli* cells were recovered in 0.4 mL Luria-Bertani medium (LB) (Research Products Int.) at 37 ºC with shaking at 220 rpm for 30 minutes before being plated on LB-agar plates with 100 µg/mL ampicillin (LB-Amp agar plates). Single colonies from LB-agar plates were picked using sterilized toothpicks to individually inoculate 400 µL of LB containing 100 µg/mL of ampicillin (LB-Amp) in 2-mL 96-well deep-well plates. The plates were incubated at 37 ºC and shaken at 220 rpm for 16–18 hours. The following morning, 50 µL of preculture from each well were added to the wells of a 96-well flat-bottom tissue culture plate (ThermoFisher) preloaded with 50 µL of 50% glycerol solution. These glycerol stocks were stored at -80 °C. Additionally, the sequences of protoglobin genes contained in every well were sequenced using evSeq.[6]

**Rearray of Single-Site Mutants:**

Single-site mutagenesis mutants were rearrayed from the previously described randomly picked plate to reduce screening burden and to collect data in triplicate. The sequences of protoglobin genes contained in every well of the randomly picked plates were collected using evSeq,[6] yielding the following plate maps:

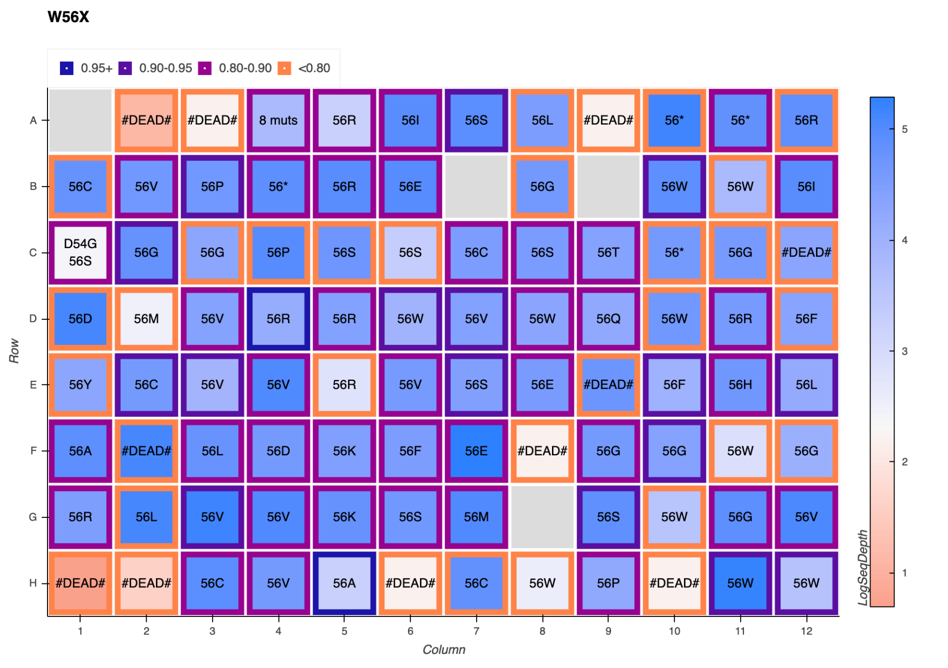


**Figure S2.** evSeq data for single site-saturation mutagenesis of ParLQ at site W56.

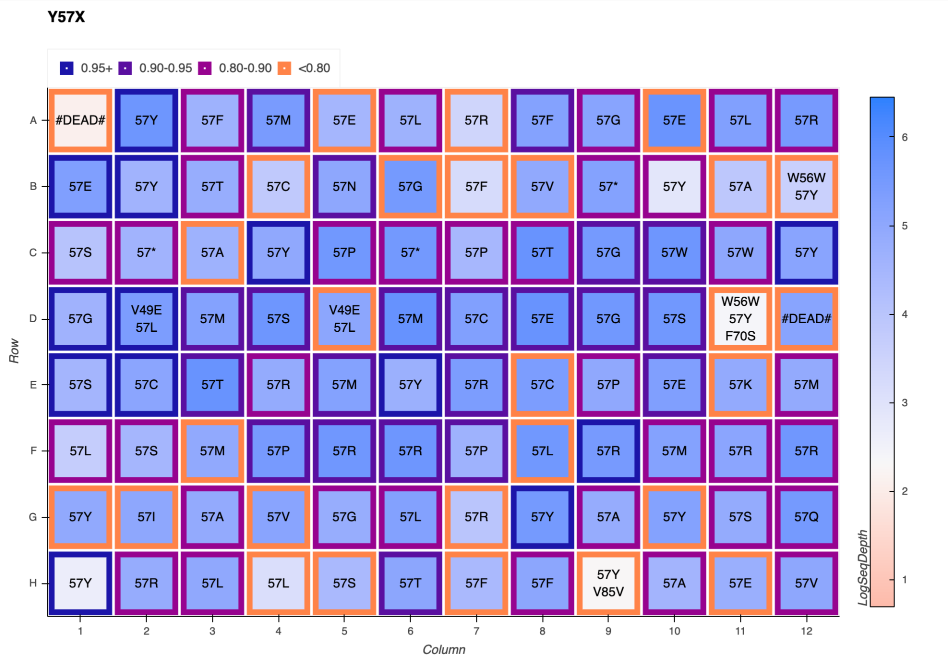


**Figure S3.** evSeq data for single site-saturation mutagenesis of ParLQ at site Y57.

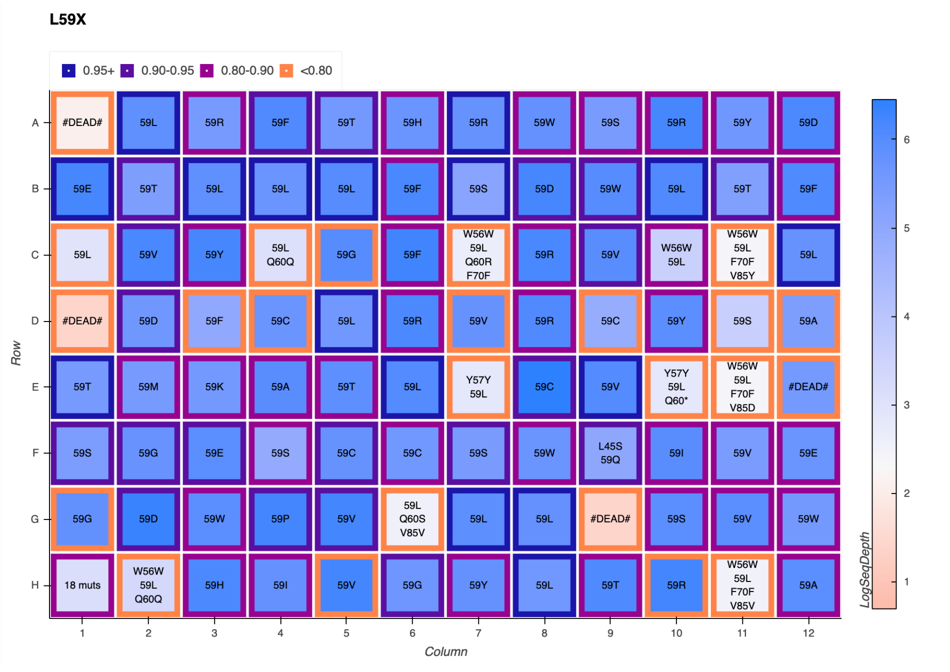


**Figure S4.** evSeq data for single site-saturation mutagenesis of ParLQ at site L59.

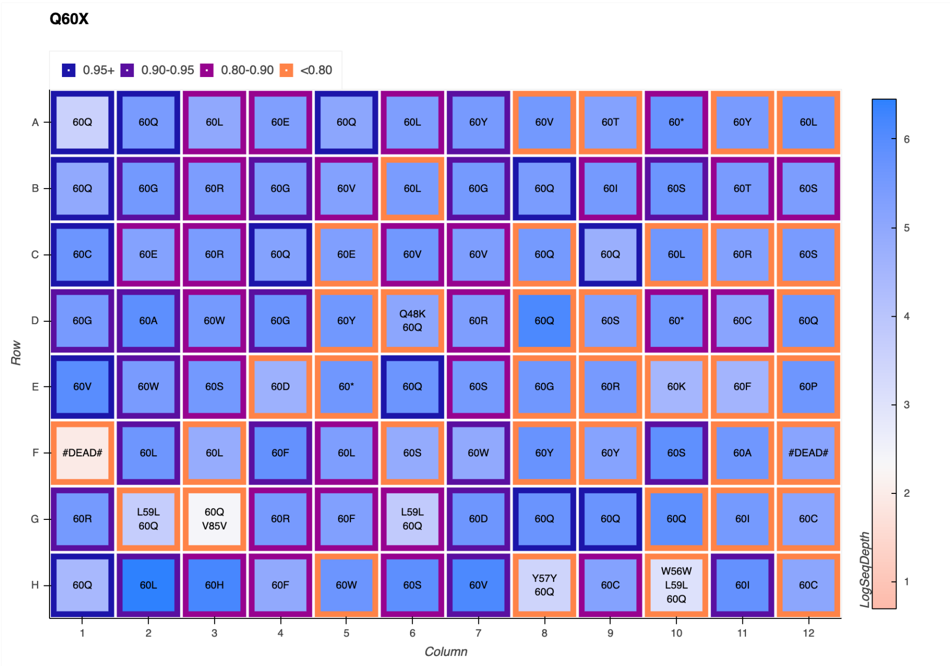


**Figure S5.** evSeq data for single site-saturation mutagenesis of ParLQ at site Q60.

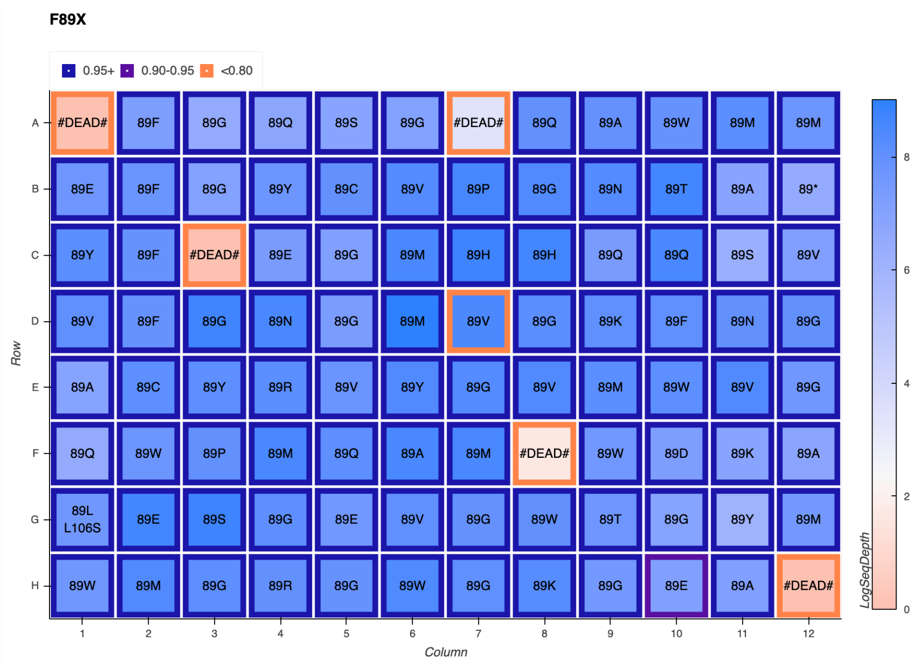


**Figure S6.** evSeq data for single site-saturation mutagenesis of ParLQ at site F89.

Protoglobin mutants for each site were rearrayed from wells containing each attained single-site mutant. However, to simulate screening of a random library, if a mutant was not found in this round of sequencing, then it was excluded from screening. In cases where a mutation was found multiple times on a plate, the well for that mutation with the highest confidence and sequencing depth was selected for rearray plate inoculation. Libraries were arrayed in the following pattern:

**Table S6.** Single site mutants were rearrayed in triplicate. If a mutation was not observed in the picked library, then the wells corresponding to that mutation are left sterile.

| | 1 | 2 | 3 | 4 | 5 | 6 | 7 | 8 | 9 | 10 | 11 | 12 |
|---|---|---|---|---|---|---|---|---|---|---|---|---|
| A | Empty | Empty | Empty | Empty | Empty | Empty | Empty | Empty | Empty | Empty | Empty | Empty |
| B | Empty | A | C | D | E | F | G | H | I | K | L | Empty |
| C | Empty | A | C | D | E | F | G | H | I | K | L | Empty |
| D | Empty | A | C | D | E | F | G | H | I | K | L | Empty |
| E | Empty | M | N | P | Q | R | S | T | V | W | Y | Empty |
| F | Empty | M | N | P | Q | R | S | T | V | W | Y | Empty |
| G | Empty | M | N | P | Q | R | S | T | V | W | Y | Empty |
| H | Empty | Empty | Empty | Empty | Empty | Empty | Empty | Empty | Empty | Empty | Empty | Empty |

The center 60 wells of a 2-mL 96-well deep-well plate were filled with 400 µL LB-Amp. Previously generated 96-well plates were removed from -80 °C storage and placed on dry ice. Pipet tips were used to scratch the frozen glycerol stock surface and used to inoculate the aforementioned deep-well plate according to **Table S6**. These overnight cultures were incubated at 37 ºC and shaken at 220 rpm for 16–18 hours. The following morning, 50 µL of overnight culture from each well were added to the wells of a 96-well flat-bottom tissue culture plate (ThermoFisher) preloaded with 50 µL of 50% glycerol solution. These glycerol stocks were stored at -80°C for future inoculation.

**Cloning for Multisite-Saturation Mutagenesis:**

Mutations were simultaneously incorporated as with single site-saturation mutagenesis using the ParLQ_quadNNK primers. Upon transformation, the 4-site library transformation was recovered in 0.4 mL of LB medium at 37 °C and 220 rpm for 30 minutes. From this recovered transformation mixture, 50 µL was transferred into 6 mL of LB-Amp in a 15-mL culture tube. This culture was allowed to shake overnight at 37 °C and 220 rpm. The following morning, this library overnight culture was miniprepped using a QIAprep Spin Miniprep Kit (Qiagen, Hilden, Germany). The miniprepped plasmid DNA pool was used as the new template for mutagenesis with the primers for site-saturation mutagenesis of site 89X. T7 Express Competent *E. coli* were transformed with the Gibson products for the new five-site library using the recommended protocol. Upon heat-shock transformation, the freshly transformed *E. coli* cells were recovered in 0.4 mL LB medium at 37 ºC with shaking at 220 rpm for 30 minutes before being plated on LB-Amp agar plates. Single colonies from LB-agar plates were picked with sterilized toothpicks to individually inoculate 400 µL of LB-Amp in 2-mL 96-well deep-well plates across four separate plates. The plates were incubated at 37 ºC and shaken at 220 rpm for 16–18 hours. The following morning, 50 µL of preculture from each well were added to the wells of a 96-well flat-bottom tissue culture plate (ThermoFisher) preloaded with 50 µL of 50% glycerol solution. These glycerol stocks were stored at -80°C for future inoculation. Additionally, the sequences of protoglobin genes contained in every well were sequenced using LevSeq sequencing.[7]

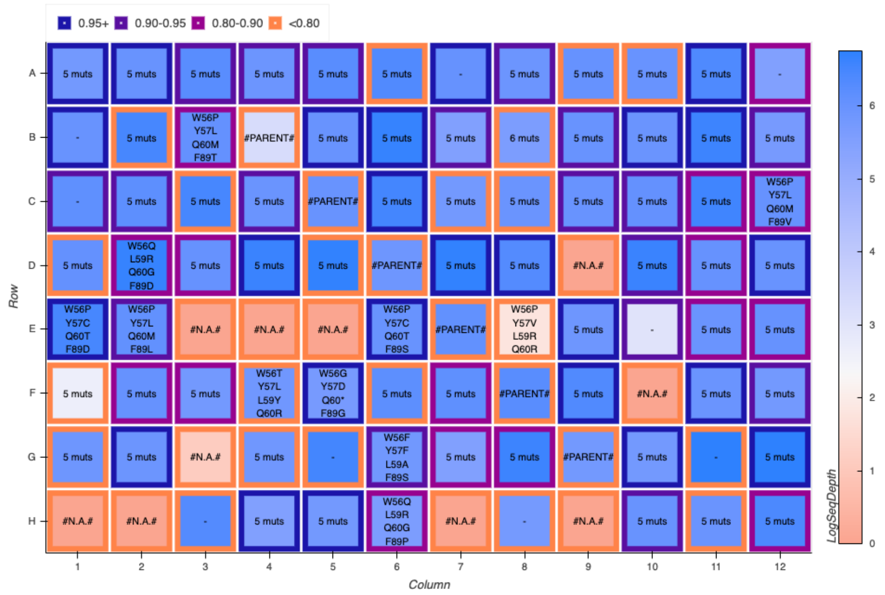


**Figure S7.** LevSeq plate sequencing data for plate 1 of the randomly generated multi-mutant library.

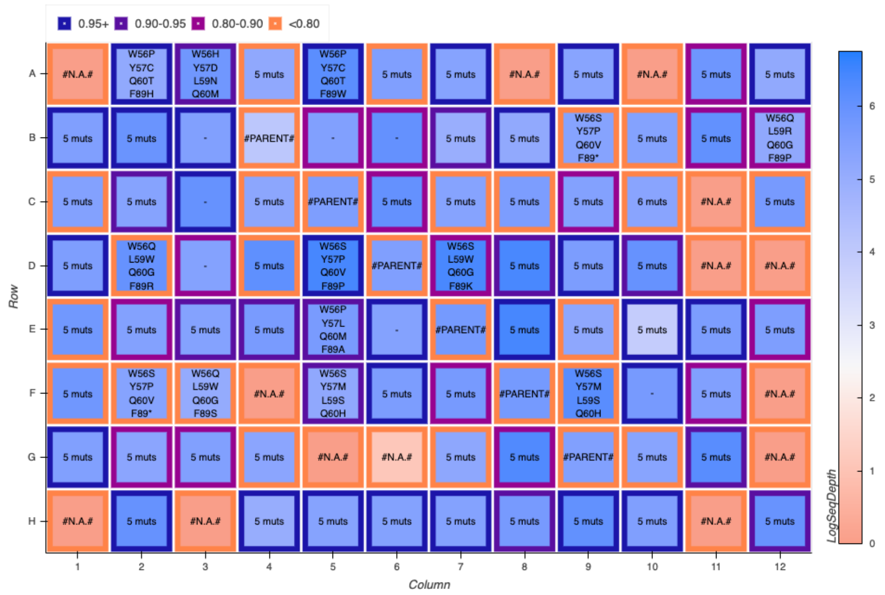


**Figure S8.** LevSeq plate sequencing data for plate 2 of the randomly generated multi-mutant library.

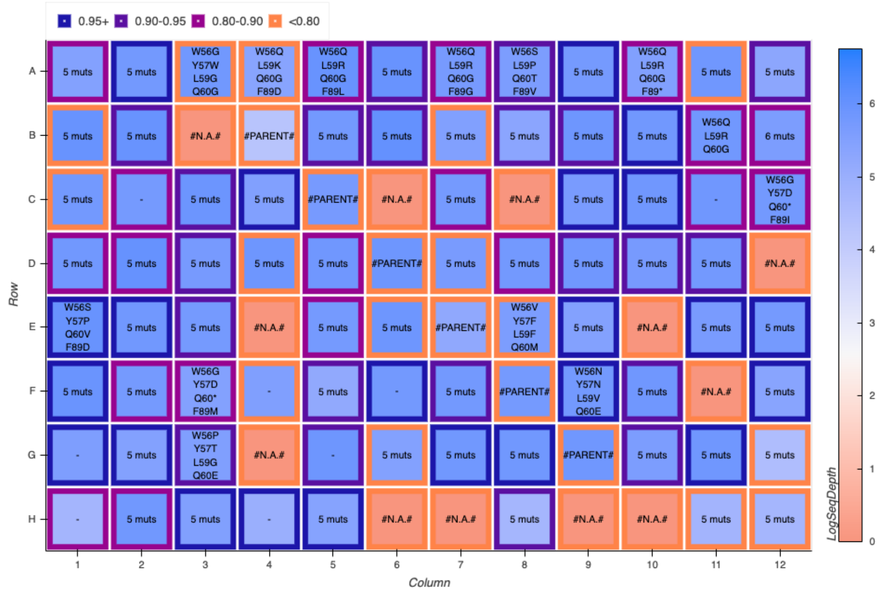


**Figure S9.** LevSeq plate sequencing data for plate 3 of the randomly generated multi-mutant library.

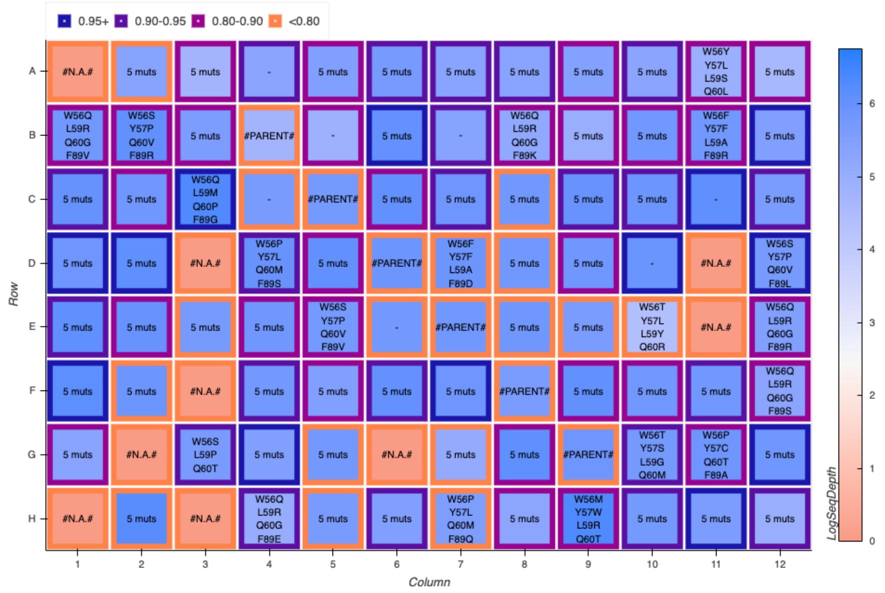


**Figure S10.** LevSeq plate sequencing data for plate 4 of the randomly generated multi-mutant library.

*Protocols for the Cloning of ALDE Predicted Sequences*

**96-Well Plate Gibson Protocol:**

Exact genes encoding ParLQ mutants predicted by Active Learning-Assisted Directed Evolution (ALDE) were synthesized and delivered by Elegen Corp. (San Carlos, CA). DNA fragments were received as dry residues in 96-well PCR plates in 2–4-μg quantities. These DNA samples were dissolved in 100 μL of double-distilled $H_2O$ ($ddH_2O$), yielding concentrations between 20–40 ng/μL. A 0.7-μL aliquot of these resuspended gene solutions were added to the wells of a 96-well PCR plate (Globe Scientific Inc., Mahwah, NJ). To each well of this plate was then added 1.0 μL of an aqueous solution containing 60 ng/μL of linearized pET-22b(+) backbone with overhangs designed for Gibson ligation with the DNA samples. Finally, to each well were added 5 μL of Gibson assembly mix.[5] The 96-well plate was then incubated at 50 °C for 60 minutes, after which the resulting Gibson products are placed on ice. These Gibson products could then either be directly used for transformation or stored at -20 °C for future use.

**Transformation Protocol for 96-Well Plate Scale:**

To each well of the previously described Gibson assembly plate were added 5 μL of T7 Express Competent *E. coli*. The cell solutions were allowed to incubate on ice for 20 minutes, after which they were heat-shocked at 42 °C for 10 seconds in a water bath. The cells were then recovered with the addition of 100 μL of LB but without outgrowth at 37 °C. Immediately, 10 μL of each transformation mixture were used to inoculate the wells of a 2-mL 96-well deep-well plate in which the wells had been preloaded with 400 μL LB-Amp. This plate was incubated at 37 ºC and shaken at 220 rpm for 16–18 hours. The following morning the plate was removed from the incubator and allowed to sit at room temperature for 8–10 hours. After this rest phase, 1 μL from each well was used to reinoculated yet another 96-well deep-well plate preloaded with 400 μL LB-

Amp. This cell passage plate was incubated at 37 ºC and shaken at 220 rpm for 16–18 hours. The following morning, 50 µL of preculture from each well were added to the wells of a 96-well flat-bottom tissue culture plate (ThermoFisher) preloaded with 50 µL of 50% glycerol solution. These glycerol stocks were stored at -80°C for future use.

### Preparation of Authentic Standards

#### General Procedure for Cyclopropane Synthesis

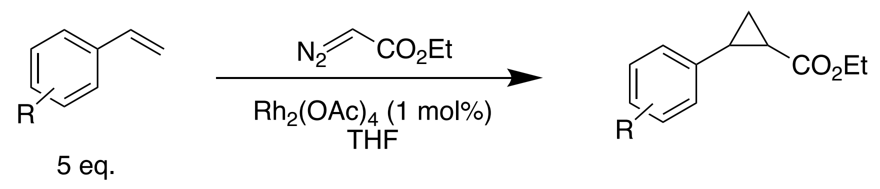


The protocol for cyclopropane authentic standard synthesis was adapted from the procedure of Doyle and coworkers.[8] Rhodium acetate dimer (40 µmol, 18 mg) was added to a 40-mL dram vial equipped with a stir bar. The vial was sealed with a septum cap. A solution of olefin (20 mmol) was dissolved in 2 mL anhydrous THF and added to the sealed vial. Separately, a solution of ethyl diazoacetate (4 mmol, 0.4 mL) was dissolved in 1.2 mL of THF. The ethyl diazoacetate solution was added to the reaction over the course of two hours while the reaction was stirred at room temperature. The reaction was allowed to proceed overnight. The crude reaction mixture was filtered through a plug of activated alumina, and the filtrate was concentrated under reduced pressure. The resultant concentrate was loaded on a SNAP Ultra silica flash cartridge and separated using an Isolera flash purification system (Biotage, Charlotte, NC) with a hexane/ethyl acetate gradient from 0–10% ethyl acetate. Fractions containing the desired product were pooled and concentrated under reduced pressure. Only in certain cases could the *trans*- and *cis*- cyclopropane products be chromatographically separated.

**Ethyl 2-(4-methoxyphenyl)cyclopropane-1-carboxylate (2a)**

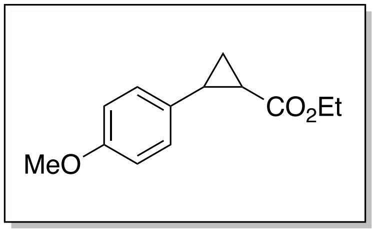


Compound **2a** was synthesized with the general procedure for cyclopropane synthesis from 4-vinylanisole (**1a**) and EDA. The *trans*- and *cis*- products were separated by column chromatography, yielding 346 mg (39% yield) of the *trans*- diastereomer and 229 mg (26% yield) of the *cis*- diastereomer were isolated. Both *trans*- and *cis*-**2a** have been previously characterized in the literature.[9]

*trans*-***2a***: $^1$H NMR (400 MHz, $CDCl_3$) δ 7.04 – 6.89 (m, 2H), 6.82 – 6.68 (m, 2H), 4.09 (q, *J* = 7.1 Hz, 2H), 3.71 (s, 3H), 2.41 (ddd, *J* = 9.2, 6.6, 4.2 Hz, 1H), 1.75 (ddd, *J* = 8.4, 5.2, 4.2 Hz, 1H), 1.53 – 1.44 (m, 2H), 1.25 – 1.14 (m, 4H).

*cis*-**2a**: $^1$H NMR (400 MHz, $CDCl_3$) δ 7.22 – 7.14 (m, 2H), 6.84 – 6.76 (m, 2H), 3.89 (q, *J* = 7.1 Hz, 2H), 3.77 (s, 3H), 2.52 (td, *J* = 8.8, 7.3 Hz, 1H), 2.08 – 1.98 (m, 1H), 1.65 (ddd, *J* = 7.5, 5.6, 5.0 Hz, 1H), 1.35 – 1.23 (m, 1H), 1.01 (t, *J* = 7.1 Hz, 3H).

**Ethyl 2-phenylcyclopropane-1-carboxylate (2b)**

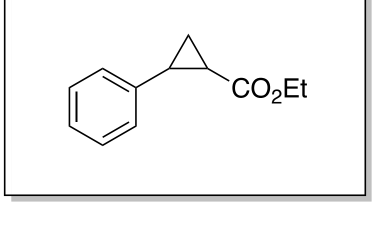


Compound **2b** was synthesized with the general procedure for cyclopropane synthesis from styrene (**1b**) and EDA. After separation by column chromatography, 124 mg (16% yield) of an 8:1 mixture of *trans*-/*cis*- isomers and 97 mg (13% yield) of pure *cis*-**2b** were isolated. Both *trans*- and *cis*-**2b** have been previously characterized in the literature.[10,11]

*trans*-**2b**: $^1$H NMR (400 MHz, $CDCl_3$) δ 7.34 – 7.21 (m, 2H), 7.24 – 7.16 (m, 1H), 7.14 – 7.06 (m, 2H), 4.17 (q, *J* = 7.1 Hz, 2H), 2.62 – 2.47 (m, 1H), 1.90 (ddd, *J* = 8.4, 5.3, 4.2 Hz, 1H), 1.60 (ddd, *J* = 9.2, 5.3, 4.5 Hz, 1H), 1.38 – 1.23 (m, 4H).

*cis*-**2b**: $^1$H NMR (400 MHz, $CDCl_3$) δ 7.29 (s, 2H), 7.33 – 7.17 (m, 2H), 3.90 (q, *J* = 7.1 Hz, 2H), 2.61 (td, *J* = 9.0, 7.5 Hz, 1H), 2.10 (ddd, *J* = 9.4, 7.8, 5.6 Hz, 1H), 1.74 (dt, *J* = 7.5, 5.4 Hz, 1H), 1.40 – 1.24 (m, 2H), 0.99 (t, *J* = 7.1 Hz, 3H).

**Ethyl 2-(*p*-tolyl)cyclopropane-1-carboxylate** (**2c**)

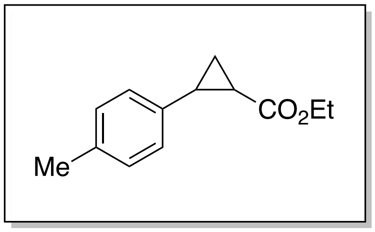


Compound **2c** was synthesized with the general procedure for cyclopropane synthesis from 1-methyl-4-vinylbenzene (**1c**) and EDA. After separation by column chromatography, 254 mg (31% yield) of a 1.92:1 mixture of *trans*-/*cis*- isomers (determined by NMR integration ratios) of **2c** was isolated. Both *trans*- and *cis*-**2c** have been previously characterized in the literature.[12] Reported *trans*- and *cis*- NMR peak assignments are assigned from the same NMR spectrum.

*trans*-**2c**: $^1$H NMR (400 MHz, $CDCl_3$) δ 7.12 – 6.95 (m, 5H), 4.23 – 4.12 (m, 2H), 2.49 (ddd, *J* = 10.0, 6.3, 4.0 Hz, 1H), 2.31 (s, 3H), 1.86 (dddd, *J* = 8.3, 5.1, 4.2, 0.8 Hz, 1H), 1.63 – 1.51 (m, 2H), 1.38 – 1.23 (m, 5H).

*cis*-**2c**: $^1$H NMR (400 MHz, $CDCl_3$) δ 7.19 – 7.09 (m, 4H), 3.89 (q, *J* = 7.1 Hz, 2H), 2.54 (q, *J* = 7.9, 7.3 Hz, 1H), 2.30 (s, 3H), 2.05 (ddd, *J* = 9.4, 7.8, 5.6 Hz, 1H), 1.68 (dt, *J* = 7.2, 5.0 Hz, 1H), 1.41 – 1.31 (m, 1H), 1.06 – 0.96 (m, 3H).

**Ethyl 2-(*m*-tolyl)cyclopropane-1-carboxylate (2d)**

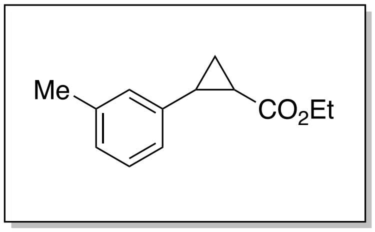


Compound **2d** was synthesized with the general procedure for cyclopropane synthesis from 1-methyl-3-vinylbenzene (**1d**) and EDA. After separation by column chromatography, 242 mg (30% yield) of a 1.87:1 mixture of *trans-*/*cis-* isomers (determined by NMR integration ratios) of **2d** was isolated. Both *trans-* and *cis*-**2d** have been previously characterized in the literature.[12] Reported *trans-* and *cis-* NMR peak assignments are assigned from the same NMR spectrum.

*trans*-**2d:** $^1$H NMR (400 MHz, $CDCl_3$) δ 7.04 – 6.98 (m, 2H), 6.94 – 6.87 (m, 2H), 4.17 (q, *J* = 7.1 Hz, 2H), 2.48 (ddd, *J* = 9.2, 6.5, 4.2 Hz, 1H), 2.33 – 2.32 (m, 3H), 1.89 (ddd, *J* = 8.4, 5.3, 4.2 Hz, 1H), 1.58 (ddd, *J* = 9.2, 5.3, 4.5 Hz, 2H), 1.36 – 1.19 (m, 5H).

*cis*-**2d**: $^1$H NMR (400 MHz, $CDCl_3$) δ 7.19 – 7.04 (m, 5H), 3.89 (q, *J* = 7.1 Hz, 2H), 2.55 (q, *J* = 8.6 Hz, 1H), 2.31 (d, *J* = 0.7 Hz, 3H), 2.06 (ddd, *J* = 9.3, 7.8, 5.6 Hz, 1H), 1.69 (ddd, *J* = 7.5, 5.6, 5.0 Hz, 1H), 0.99 (t, *J* = 7.1 Hz, 3H).

**Ethyl 2-(*o*-tolyl)cyclopropane-1-carboxylate (2e)**

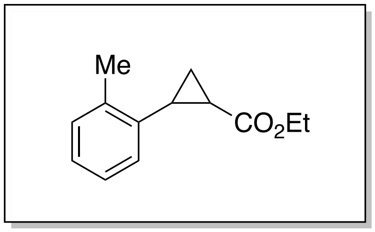


Compound **2e** was synthesized with the general procedure for cyclopropane synthesis from 1-methyl-2-vinylbenzene (**1e**) and EDA. The *trans*- and *cis*- products were separated by column chromatography, yielding 118 mg (14% yield) of the *trans*- diastereomer and 81 mg (10% yield) of the *cis*- diastereomer were isolated. Both *trans*- and *cis*-**2e** have been previously characterized in the literature.[12]

*trans*-**2e**: $^1$H NMR (400 MHz, $CDCl_3$) δ 7.24 – 7.06 (m, 3H), 7.03 – 6.95 (m, 1H), 4.30 – 4.12 (m, 2H), 2.56 – 2.42 (m, 1H), 2.39 (s, 3H), 1.79 (dtd, *J* = 8.3, 4.8, 0.8 Hz, 1H), 1.57 (ddd, *J* = 9.3, 5.1, 4.3 Hz, 1H), 1.30 (td, *J* = 7.1, 0.8 Hz, 4H).

*cis*-**2e**: $^1$H NMR (400 MHz, $CDCl_3$) δ 7.25 – 7.16 (m, 1H), 7.21 – 7.07 (m, 3H), 3.85 (q, *J* = 7.1 Hz, 2H), 2.45 (q, *J* = 8.4 Hz, 1H), 2.16 (ddd, *J* = 9.1, 7.9, 5.4 Hz, 1H), 1.75 (dt, *J* = 7.6, 5.2 Hz, 1H), 1.61 – 1.55 (m, 1H), 1.40 – 1.26 (m, 1H), 0.93 (t, *J* = 7.1 Hz, 3H).

**Ethyl 2-(4-chlorophenyl)cyclopropane-1-carboxylate (2f)**

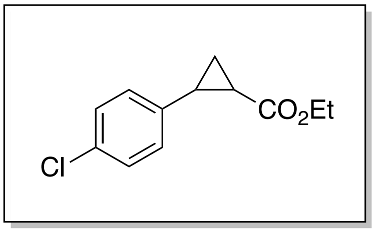


Compound **2f** was synthesized with the general procedure for cyclopropane synthesis from 1-chloro-4-vinylbenzene (**1f**) and EDA. After separation by column chromatography, 226 mg (25% yield) of a 2:1 mixture of *trans-*/*cis-* isomers (determined by NMR integration ratios) of **2f** was isolated. Both *trans-* and *cis-***2f** have been previously characterized in the literature.[13,14]

*trans*-**2f**: $^1$H NMR (400 MHz, $CDCl_3$) δ 7.23 – 7.18 (m, 2H), 7.09 – 7.00 (m, 2H), 4.18 (q, $J$ = 7.0 Hz, 2H), 2.52 – 2.44 (m, 1H), 1.87 (ddd, $J$ = 8.5, 5.3, 4.1 Hz, 1H), 1.61 (ddd, $J$ = 9.2, 5.4, 4.6 Hz, 2H), 1.29 (t, $J$ = 7.1 Hz, 3H).

*cis*-**2f**: $^1$H NMR (400 MHz, $CDCl_3$) δ 7.27 – 7.22 (m, 4H), 3.90 (q, $J$ = 7.1 Hz, 2H), 2.57 – 2.50 (m, 1H), 2.08 (ddd, $J$ = 9.2, 7.8, 5.6 Hz, 1H), 1.67 (dt, $J$ = 7.5, 5.4 Hz, 1H), 1.39 – 1.31 (m, 1H), 1.02 (t, $J$ = 7.1 Hz, 3H).

**Ethyl 2-(4-bromophenyl)cyclopropane-1-carboxylate (2g)**

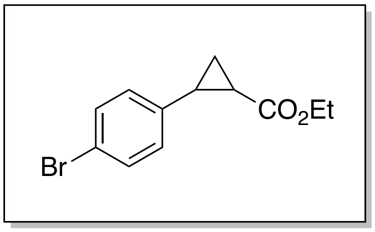


Compound **2g** was synthesized with the general procedure for cyclopropane synthesis from 1-bromo-4-vinylbenzene (**1g**) and EDA. The *trans-* and *cis-* products were separated by column chromatography, yielding 334 mg (31% yield) of the *trans*- diastereomer and 297 mg (28% yield) of the *cis*- diastereomer were isolated. Both *trans*- and *cis*-**2g** have been previously characterized in the literature.[9,15]

*trans*-**2g**: $^{1}$H NMR (400 MHz, $CDCl_3$) δ 7.43 – 7.35 (m, 2H), 7.00 – 6.93 (m, 2H), 4.17 (q, $J$ = 7.1 Hz, 2H), 2.47 (ddd, $J$ = 9.4, 6.5, 4.1 Hz, 1H), 1.91 – 1.82 (m, 1H), 1.60 (dt, $J$ = 9.5, 5.0 Hz, 1H), 1.32 – 1.22 (m, 4H).

*cis*-**2g**: $^{1}$H NMR (400 MHz, $CDCl_3$) δ 7.47 – 7.34 (m, 2H), 7.14 (d, $J$ = 8.2 Hz, 2H), 3.90 (q, $J$ = 7.1 Hz, 2H), 2.50 (q, $J$ = 8.5 Hz, 1H), 2.08 (ddd, $J$ = 8.9, 7.9, 5.6 Hz, 1H), 1.67 (dt, $J$ = 7.5, 5.4 Hz, 1H), 1.57 (d, $J$ = 1.8 Hz, 1H), 1.35 (dt, $J$ = 8.4, 4.2 Hz, 1H), 1.34 – 1.22 (m, 0H), 1.02 (t, $J$ = 7.1 Hz, 3H).

**Ethyl 2-(4-(trifluoromethyl)phenyl)cyclopropane-1-carboxylate** (**2h**)

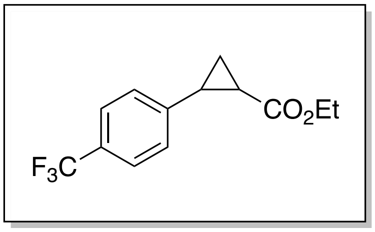


Compound **2h** was synthesized with the general procedure for cyclopropane synthesis from 1-(trifluoromethyl)-4-vinylbenzene (**1h**) and EDA, with the exception that the reaction was on a 1-mmol basis for EDA with all corresponding quantities being adjusted accordingly. After separation by column chromatography, 89 mg (30% yield) of a 3:1 mixture of *trans*-/*cis*- isomers (determined by NMR integration ratios) of **2h** was isolated. Both *trans*- and *cis*-**2h** have been previously characterized in the literature.[12]

*trans*-**2h**: $^1$H NMR (400 MHz, $CDCl_3$) δ 7.57 – 7.49 (m, 2H), 7.22 – 7.18 (m, 1H), 4.18 (q, *J* = 7.2 Hz, 2H), 2.58 – 2.51 (m, 1H), 1.94 (ddd, *J* = 8.5, 5.4, 4.2 Hz, 1H), 1.66 (ddd, *J* = 9.2, 5.4, 4.7 Hz, 1H), 1.36 – 1.31 (m, 1H), 1.29 (t, *J* = 7.1 Hz, 3H).

*cis*-**2h**: $^1$H NMR (400 MHz, $CDCl_3$) δ 7.44 (d, *J* = 4.1 Hz, 4H), 7.31 (dq, *J* = 7.5, 0.8 Hz, 2H), 3.82 (qd, *J* = 7.1, 2.1 Hz, 2H), 2.52 (d, *J* = 7.9 Hz, 1H), 2.07 (ddd, *J* = 9.3, 7.9, 5.7 Hz, 1H), 1.67 (dt, *J* = 7.5, 5.4 Hz, 1H), 1.33 (ddd, *J* = 8.7, 7.9, 5.2 Hz, 1H), 0.92 (t, *J* = 7.1 Hz, 3H).

**Ethyl 2-(naphthalen-2-yl)cyclopropane-1-carboxylate (2i)**

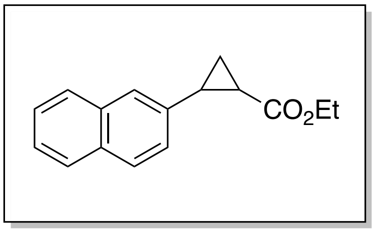


Compound **2i** was synthesized with the general procedure for cyclopropane synthesis from 2-vinylnapthylene (**1i**) and EDA. After separation by column chromatography, 467 mg (48% yield) of a 1.82:1 mixture of *trans*-/*cis*- isomers (determined by NMR integration ratios) of **2h** was isolated. Both *trans*- and *cis*-**2h** have been previously characterized in the literature.[12]

*trans*-**2i**: $^1$H NMR (400 MHz, $CDCl_3$) δ 7.85 – 7.56 (m, 7H), 4.23 – 4.16 (m, 2H), 2.71 – 2.65 (m, 1H), 2.01 (ddd, *J* = 8.4, 5.3, 4.2 Hz, 1H), 1.68 (ddd, *J* = 9.2, 5.3, 4.6 Hz, 1H), 1.43 (ddd, *J* = 8.4, 6.4, 4.6 Hz, 2H), 1.30 (t, *J* = 7.1 Hz, 3H).

*cis*-**2i**: $^1$H NMR (400 MHz, $CDCl_3$) δ 8.05 – 7.89 (m, 3H), 7.69 – 7.59 (m, 2H), 7.21 (dd, *J* = 8.5, 1.8 Hz, 2H), 3.83 (q, *J* = 7.1 Hz, 2H), 2.73 (dd, *J* = 8.0, 1.7 Hz, 1H), 2.16 (ddd, *J* = 9.3, 7.8, 5.6 Hz, 1H), 1.86 (dt, *J* = 7.5, 5.3 Hz, 1H), 1.47 – 1.44 (m, 1H).

### Preparation of Calibration Curves for Analytical Yield Determination

All reactions were assayed by gas chromatography equipped with flame ionization detection (GC-FID). To quantify cyclopropane yields, calibration curves were made with the synthesized authentic standards. Calibration curves were constructed by one of two methods.

**Calibration Curve Construction – Method A:**

Method A was used in cases where the two diastereomers of the cyclopropane could be separated by column chromatography (compounds **2a**, **2b**, **2e**, and **2g**). For these compounds, a separate 10 mM stock of cyclopropane was made for each diastereomer in a solution of 1:1 solution of ethyl acetate:cyclohexane with 1,3,5-trimethoxybenzene as an internal standard (1.0 mM concentration). This stock solution was diluted in the same of ethyl acetate:cyclohexane solution containing 1.0 mM standard to final concentrations of 0.5–10 mM cyclopropane. Samples were then analyzed by GC-FID.

**Calibration Curve Construction – Method B:**

Method A was used in cases where the two diastereomers of the cyclopropane were isolated as a mixture after column chromatography (compounds **2c**, **2d**, **2f**, **2h**, and **2i**). For these compounds, a single 20 mM stock of cyclopropane, containing both diastereomers, was prepared in a 1:1 solution of ethyl acetate:cyclohexane with 1,3,5-trimethoxybenzene as an internal standard (1.0 mM concentration). This stock solution was diluted in the same of ethyl acetate:cyclohexane solution containing 1.0 mM standard to final concentrations of 0.5–20 mM cyclopropane. Concentrations for the separate diastereomers were computed according to the molar ratios determined by NMR, and it was assumed that the presence of both diastereomers in each sample would not affect analytical signal. Samples were then analyzed by GC-FID.

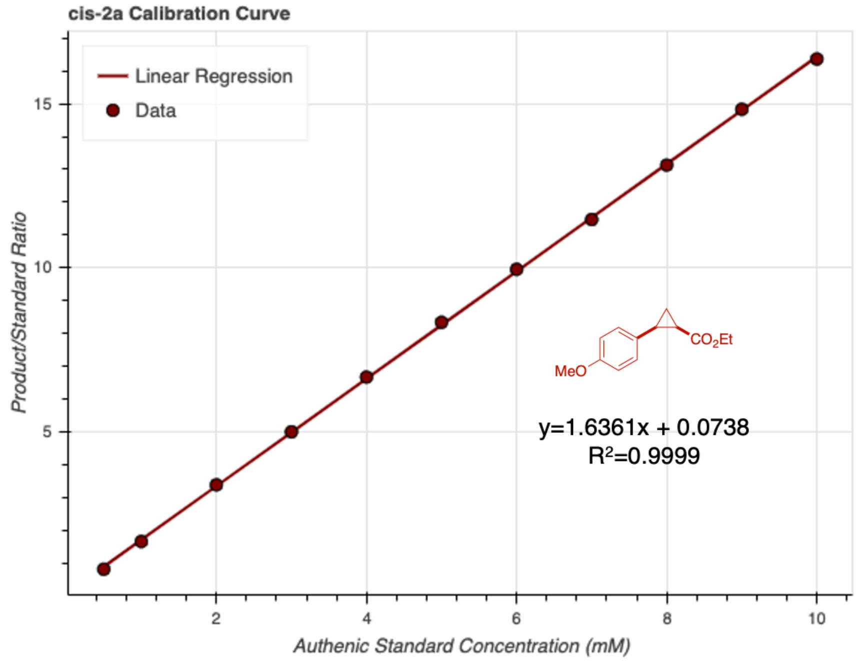


**Figure S11.** Achiral GC-FID calibration curve for *cis*-**2a**. Samples were generated using method A.

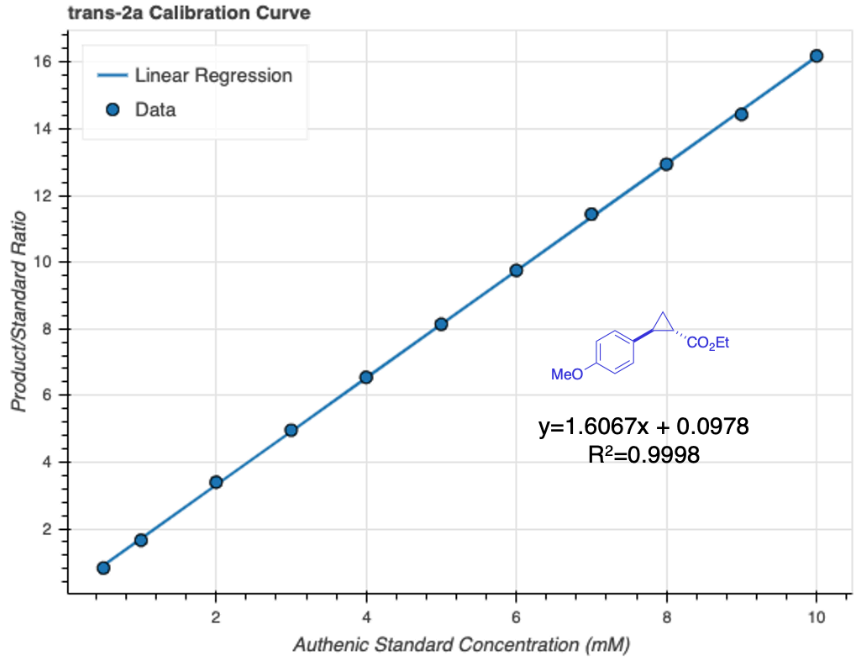


**Figure S12.** Achiral GC-FID calibration curve for *trans*-**2a**. Samples were generated using method A.

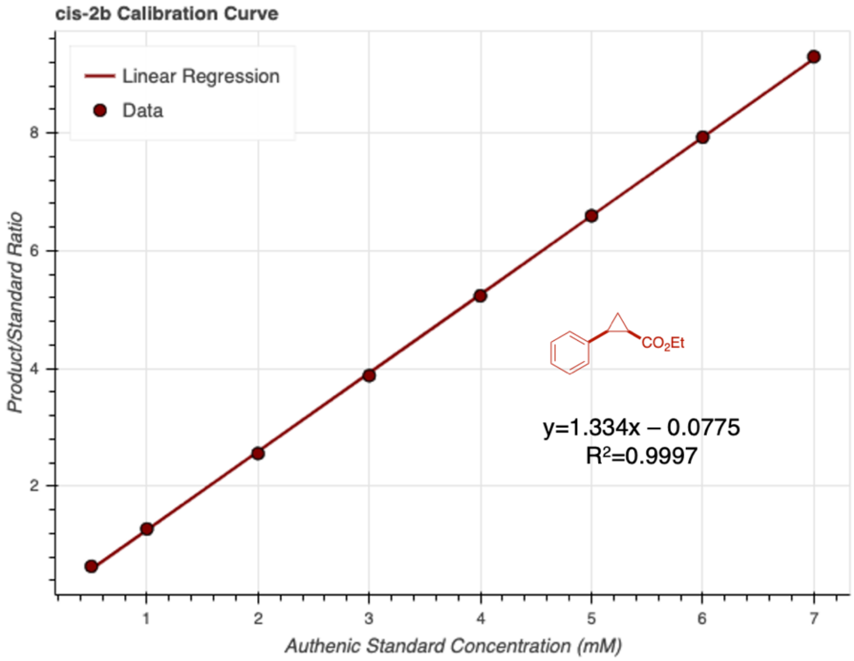


**Figure S13.** Achiral GC-FID calibration curve for *cis*-**2b**. Samples were generated using method A.

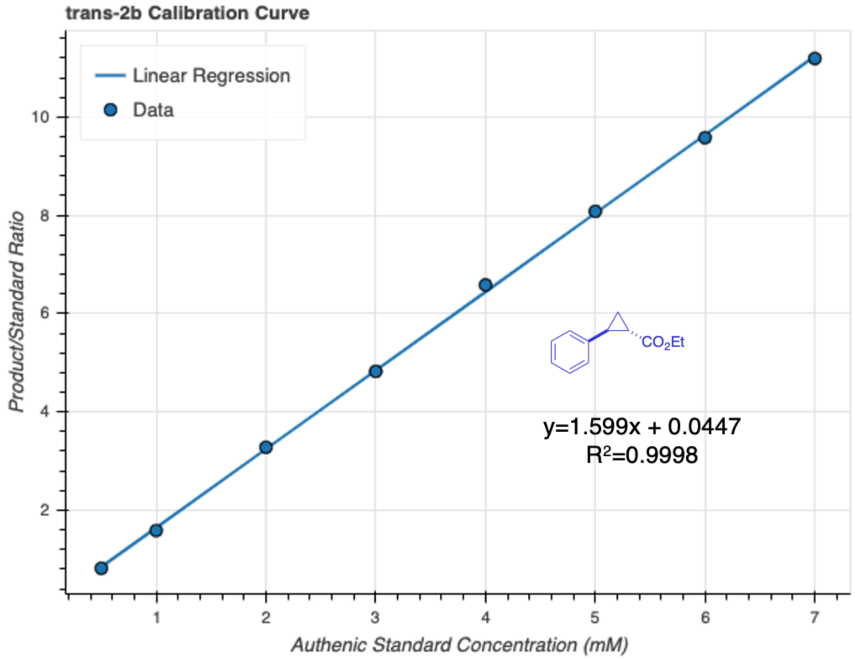


**Figure S14.** Achiral GC-FID calibration curve for *trans*-**2b**. Samples were generated using method A.

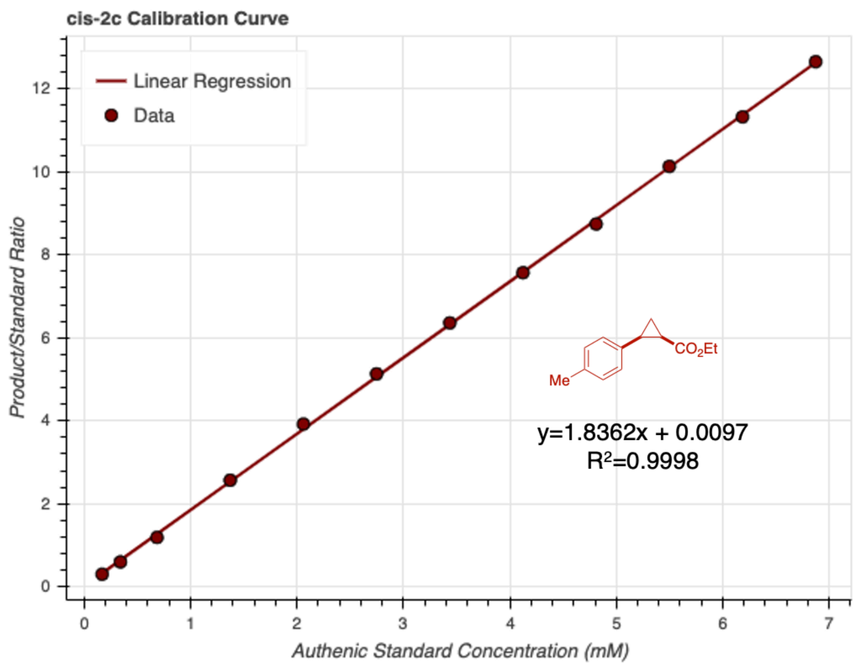


**Figure S15.** Achiral GC-FID calibration curve for *cis*-**2c**. Samples were generated using method B.

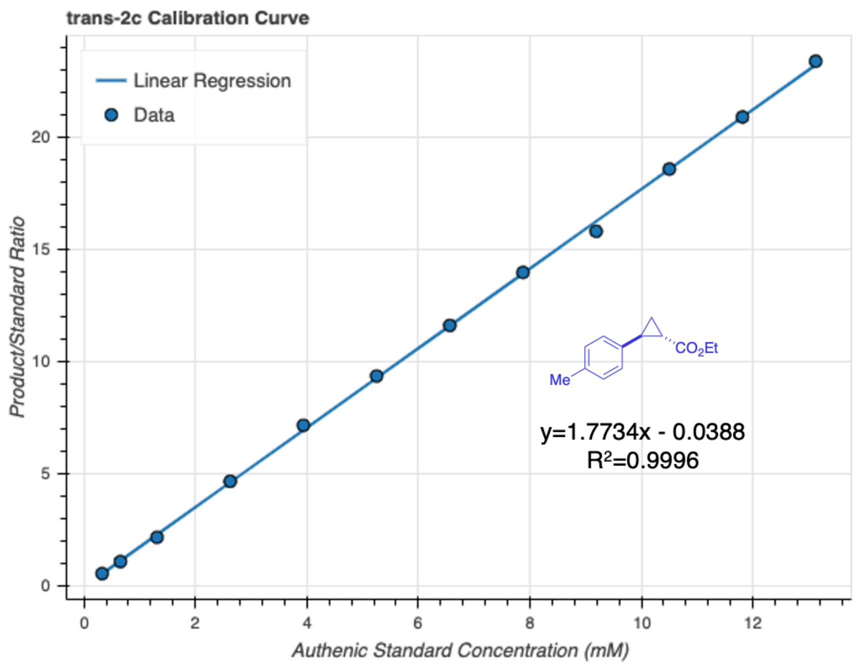


**Figure S16.** Achiral GC-FID calibration curve for *trans*-**2c**. Samples were generated using method B.

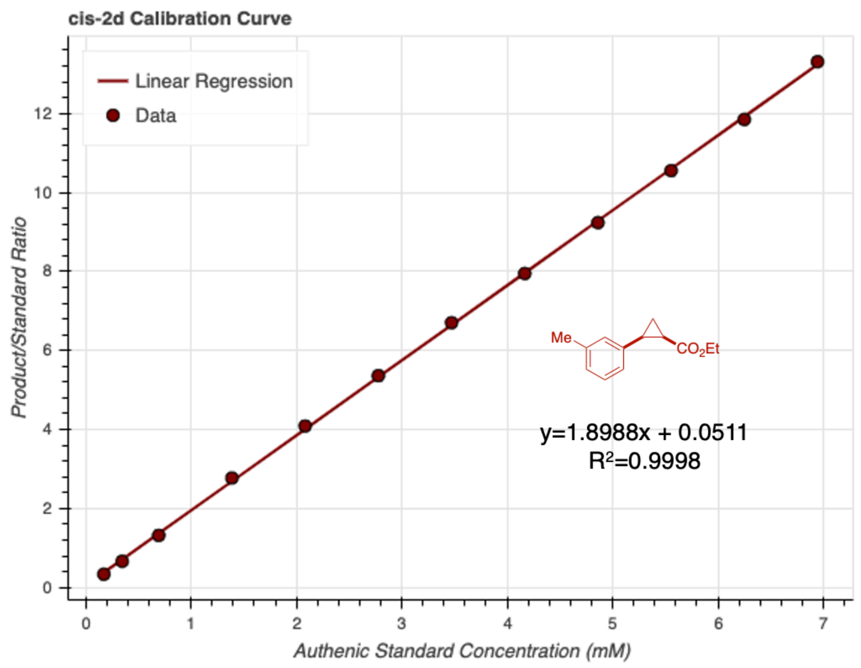


**Figure S17.** Achiral GC-FID calibration curve for *cis*-**2d**. Samples were generated using method B.

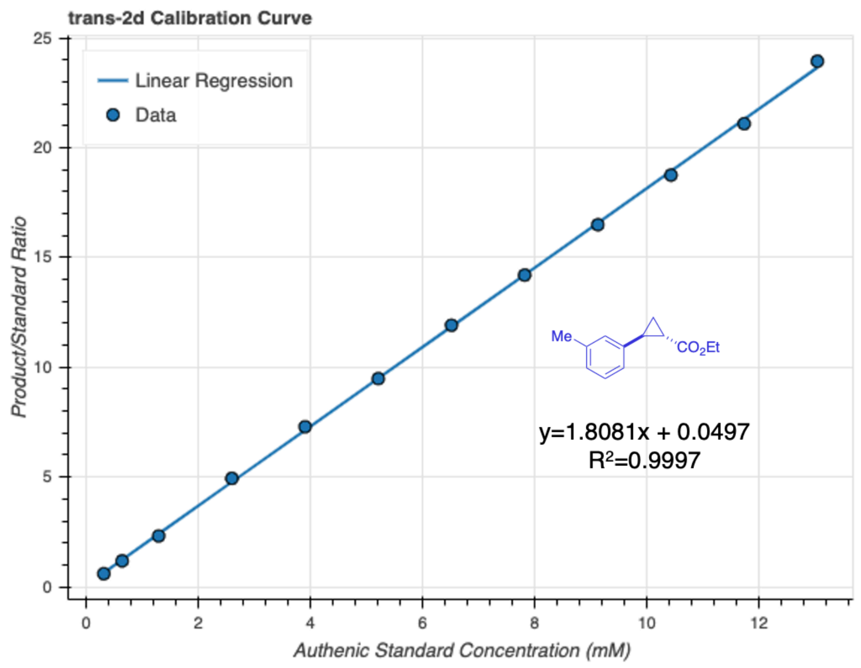


**Figure S18.** Achiral GC-FID calibration curve for *trans*-**2d**. Samples were generated using method B.

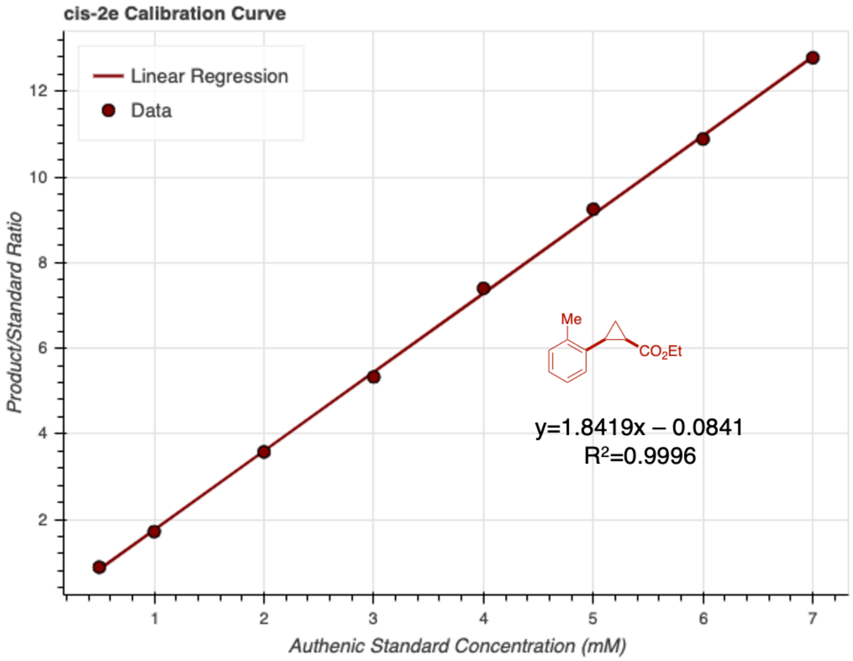


**Figure S19.** Achiral GC-FID calibration curve for *cis*-**2e**. Samples were generated using method A.

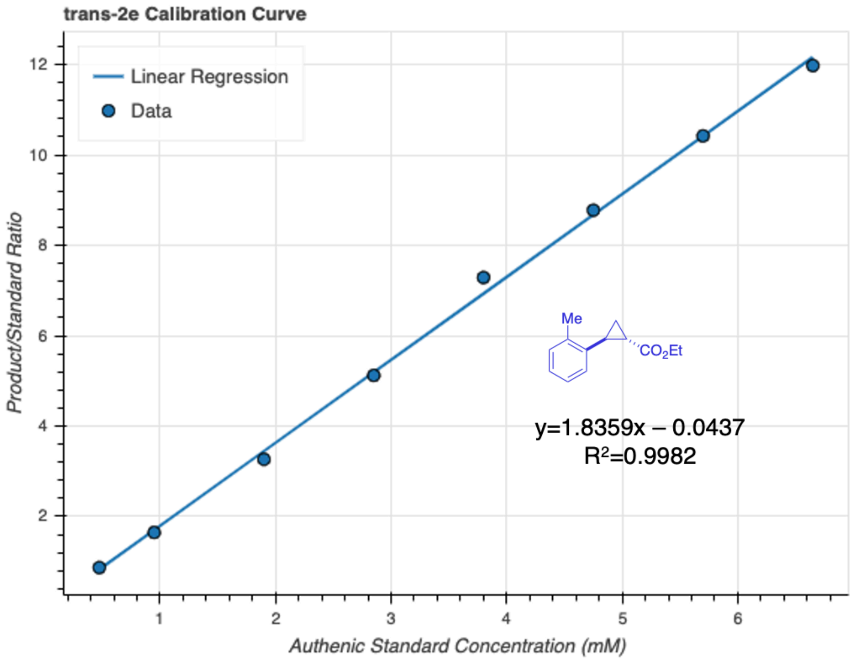


**Figure S20.** Achiral GC-FID calibration curve for *trans*-**2e**. Samples were generated using method A.

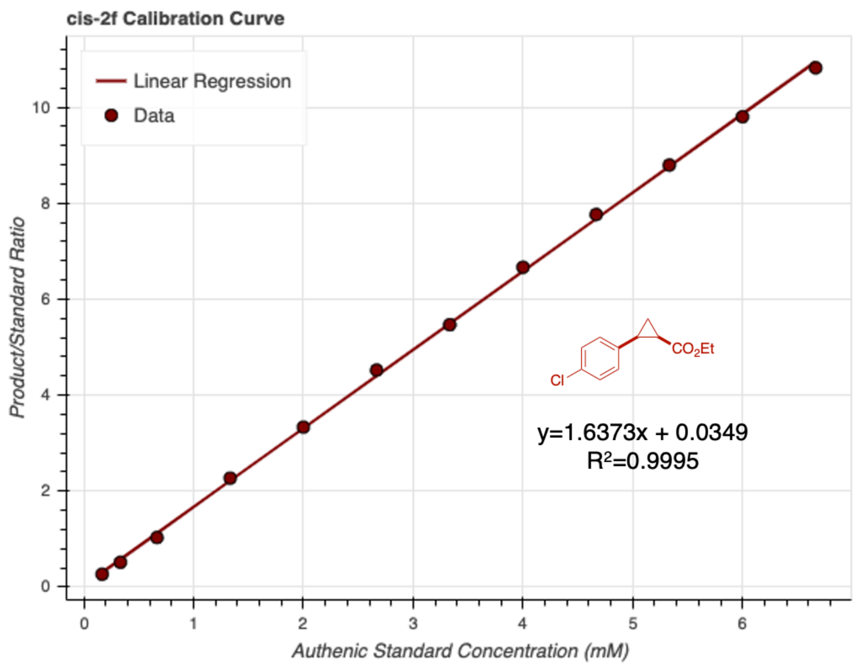


**Figure S21.** Achiral GC-FID calibration curve for *cis*-**2f**. Samples were generated using method B.

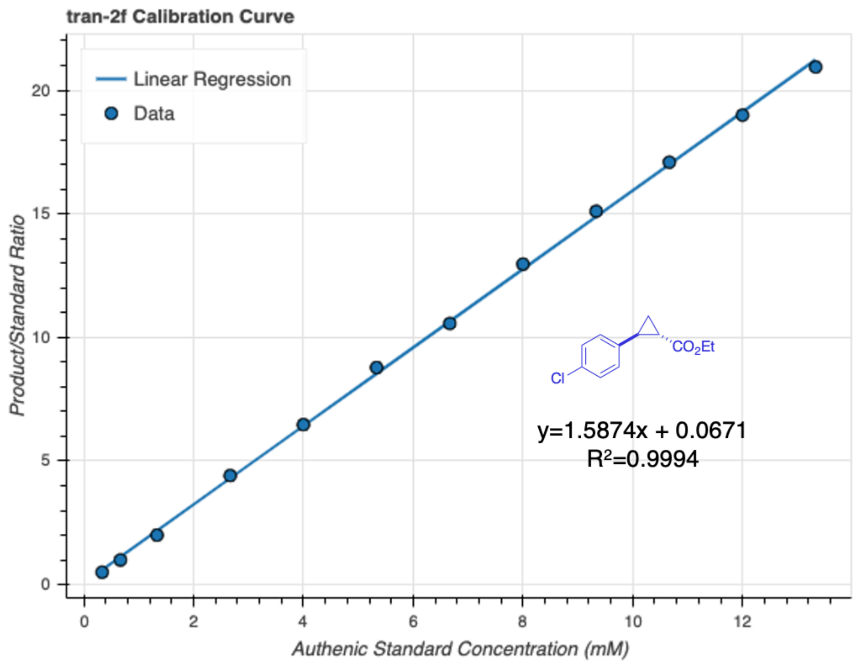


**Figure S22.** Achiral GC-FID calibration curve for *trans*-**2f**. Samples were generated using method B.

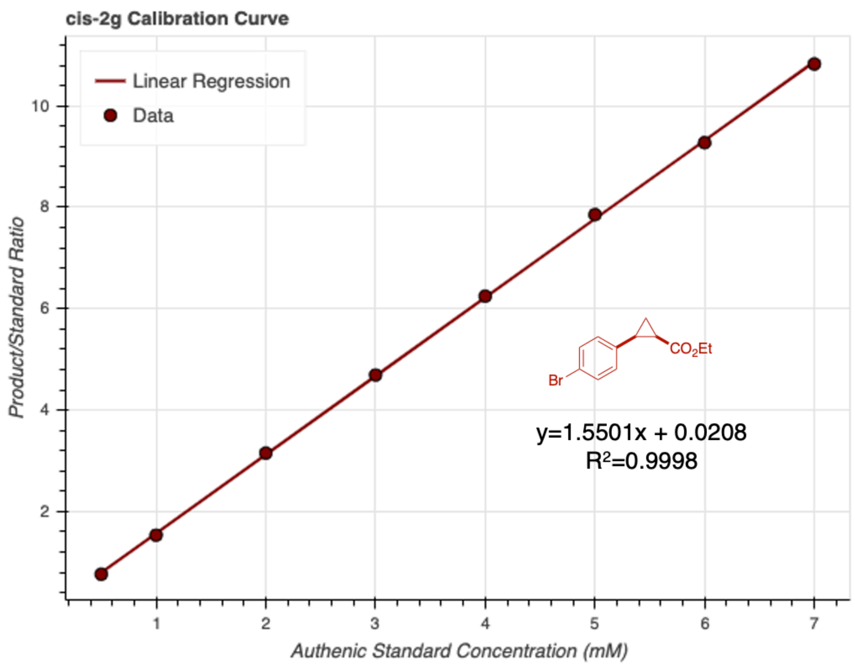


**Figure S23.** Achiral GC-FID calibration curve for *cis*-**2g**. G Samples were generated using method A.

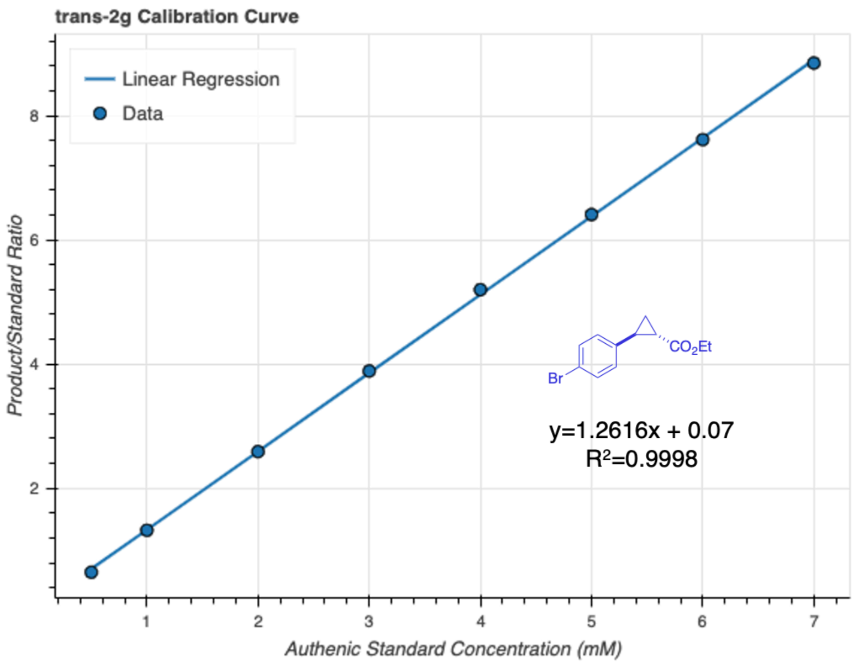


**Figure S24.** Achiral GC-FID calibration curve for *trans*-**2g**. Samples were generated using method A.

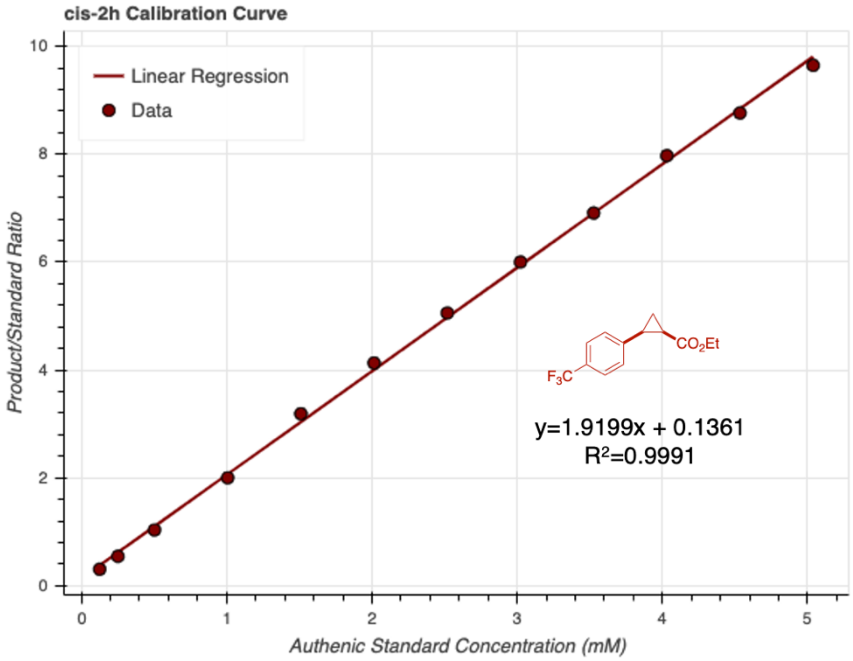


**Figure S25.** Achiral GC-FID calibration curve for *cis*-**2h**. Samples were generated using method B.

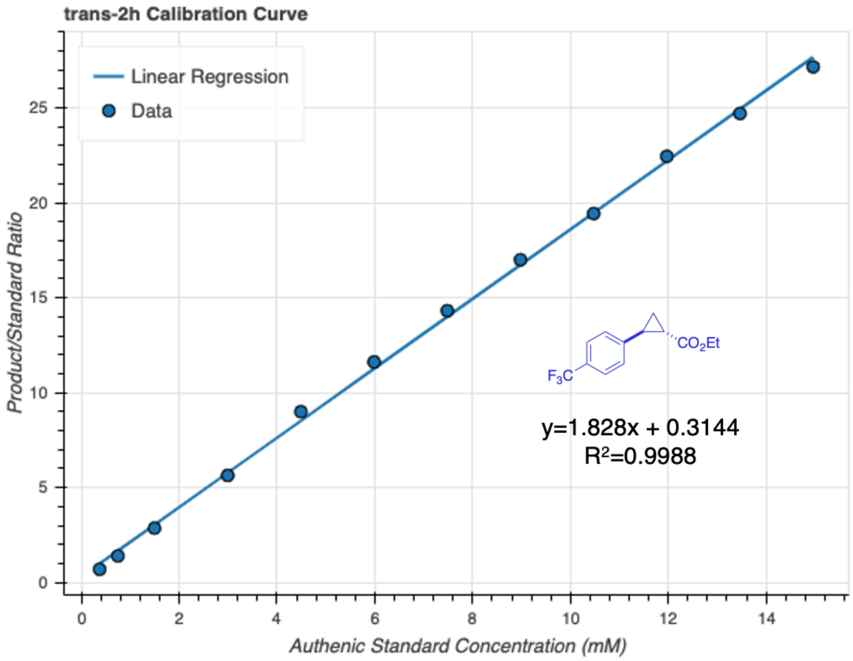


**Figure S26.** Achiral GC-FID calibration curve for *trans*-**2h**. Samples were generated using method B.

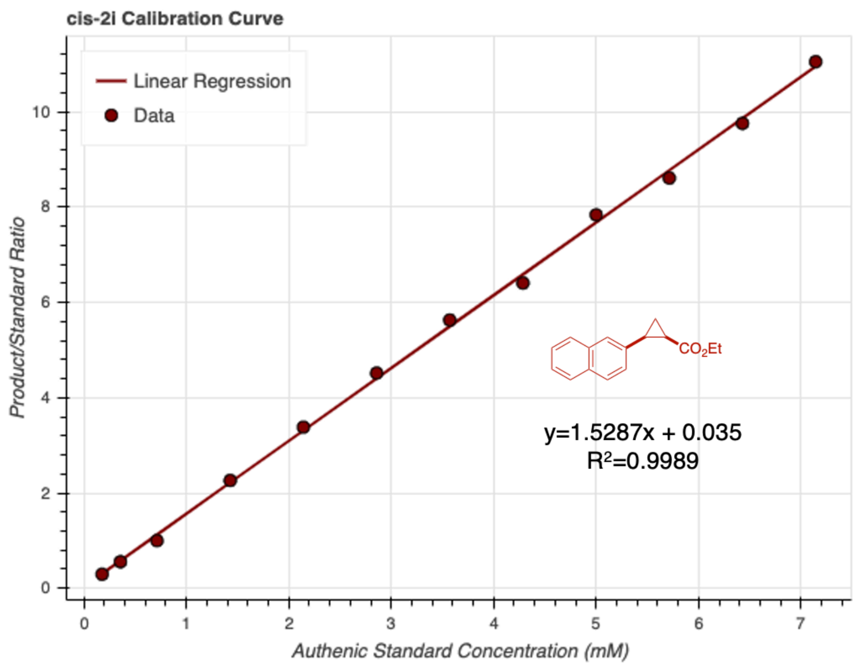


**Figure S27.** Achiral GC-FID calibration curve for *cis*-**2i**. Samples were generated using method B.

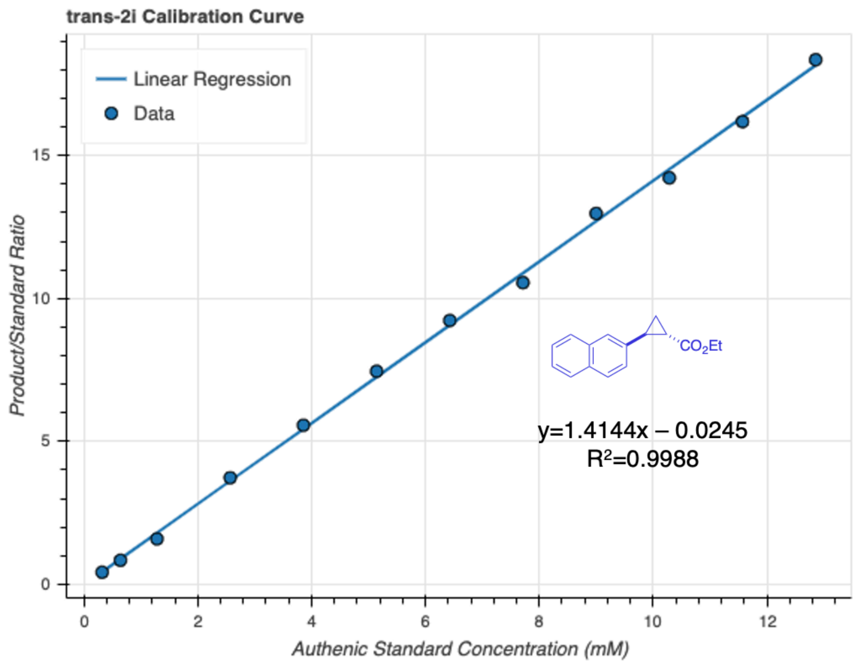


**Figure S28.** Achiral GC-FID calibration curve for *trans*-**2i**. Samples were generated using method B.

#### Control and Validation Experiments

*Protocols for Small-Scale Reaction Setup in GC Vials*

**Small-Scale Protein Expression:**

Single colonies from LB-Agar plates were picked using a sterile pipette tip and were used to inoculate 6 mL of LB-Amp in a 15-mL culture tube. Cultures were incubated at 37 ºC with shaking at 220 rpm overnight in an Innova 4000 shaking incubator. A 1-mL aliquot of each of these overnight cultures was used to inoculate 50 mL of Terrific Broth with 100 µg/mL of ampicillin (TB-Amp) (0.5% v/v starter culture in expression culture) in 125-mL unbaffled Erlenmeyer flasks. The expression cultures were incubated at 37 ºC and 220 rpm for 2.5 hours in an Innova 42 shaking incubator, at which point they are moved to room temperature for 25 minutes. Protein expression was then induced by direct addition of 50 µL of stock solutions containing 500 mM isopropyl-β-D-thiogalactoside (IPTG) and 1.0 M 5-aminolevulinic acid (ALA) such that the final concentrations were 0.5 mM and 1.0 mM, respectively. The cultures were shaken at 22 ºC and 140 rpm for 16–18 hours in an Innova 42 shaker.

**Small-Scale Biocatalytic or Control Reactions:**

The corresponding 50-mL expression cultures were pelleted (4,000 × g for 15 minutes at 4 ºC) and resuspended in 5 mL of M9-N buffer. The optical density at 600 nm ($OD_{600}$) of this suspension was measured and adjusted to $OD_{600}$ = 31.5 with the addition of more M9-N buffer. A 380 µL aliquot of the cell suspension was added to 2-mL GC vials (Agilent). For control reactions, compounds and additives were added to GC vials according to the conditions described in **Figure S29**. These whole-cell suspensions were transferred into a vinyl Coy anaerobic chamber, at which point 10 µL of a 400 mM solution of styrene in MeCN followed by 10 µL of a 600 mM solution of EDA in MeCN were added such that the final reaction concentrations were 10 mM of styrene and 15 mM of EDA. The GC vials were tightly capped with screwcaps with a septum and were allowed to shake at room temperature for 16 hours. Once complete, the reactions were transferred to a 1.7-mL Eppendorf tube and 600 μL of a 1:1 solution of ethyl acetate:cyclohexane with 1,3,5-trimethoxybenzene as an internal standard (1.0 mM concentration). The layers are vigorously mixed, and the samples were centrifuged (14,000 × g for 10 minutes at RT). Afterwards, an aliquot of the organic layer was subjected to GC analysis.

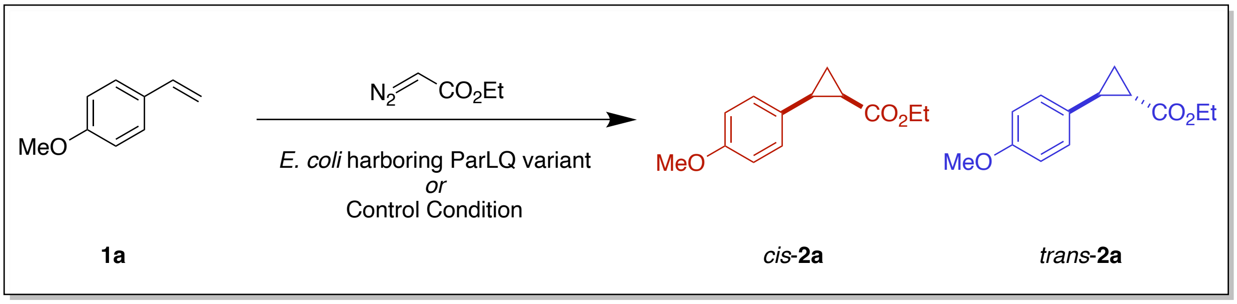


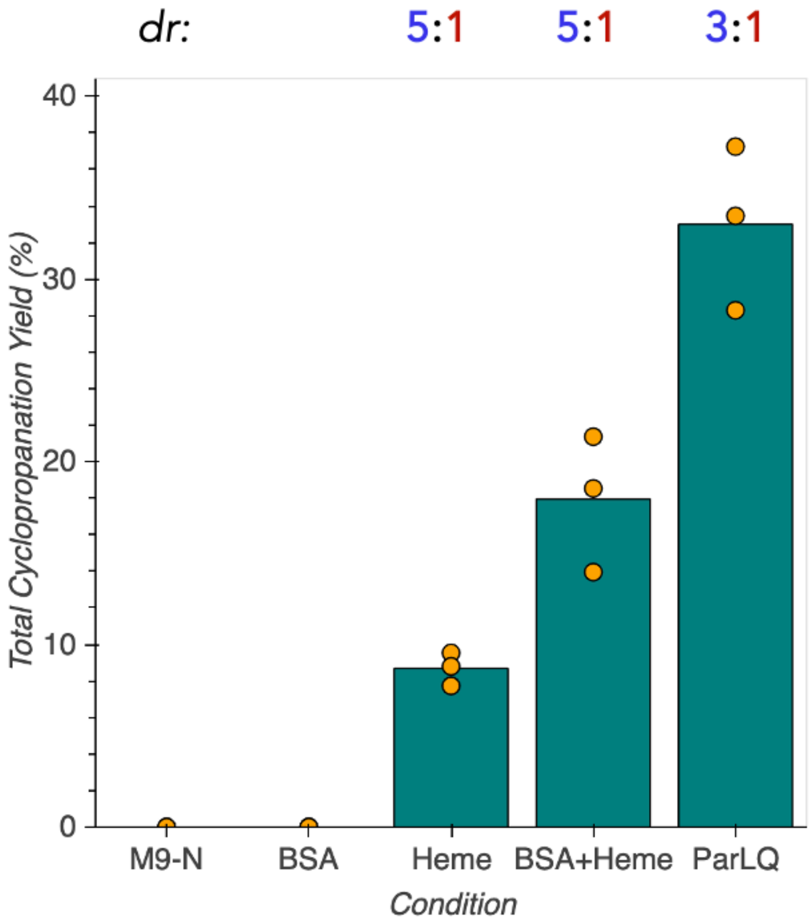


**Figure S29.** Cyclopropanation yields and selectivities for control conditions relative to ParLQ. All reactions are run in 380 µL M9-N (pH=7.6) and 20 µL MeCN. BSA was loaded at 1 mg/mL and heme was loaded at 1 mM. Diastereoselectivities are shown above the plot as the trans:cis cyclopropane ratio.

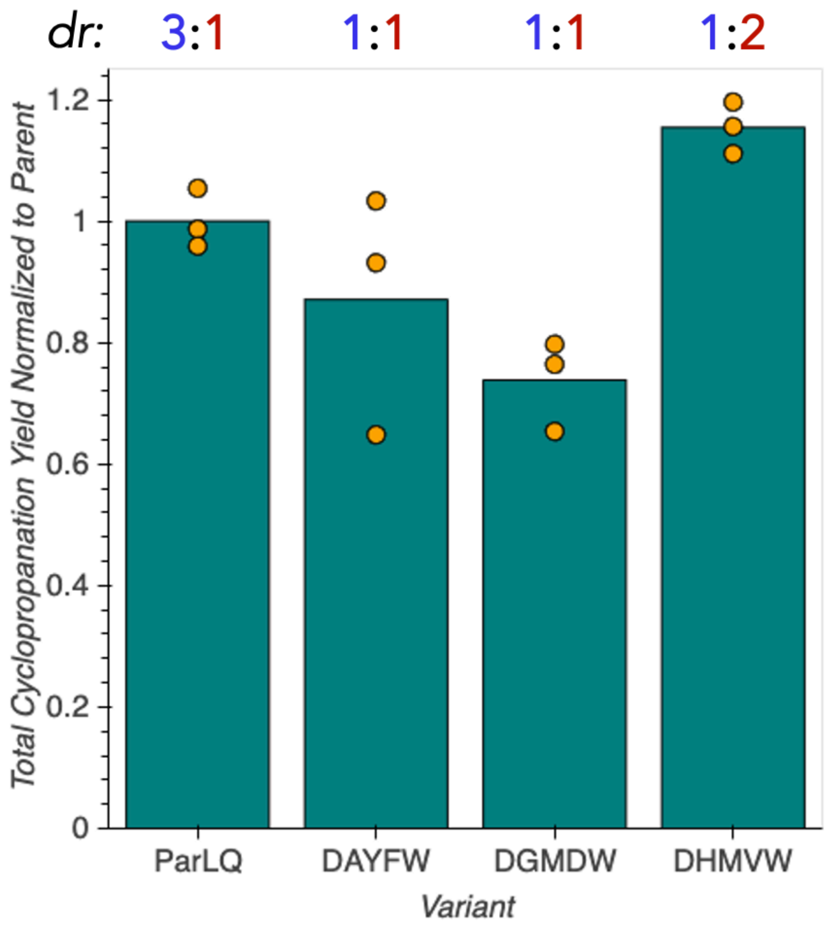


**Figure S30.** Cyclopropanation yields and selectivities for select 5-site multi-mutants of ParLQ. These variants are the result of the recombination of the single-site mutations which showed the most improvement for various possible acquisition functions. DAYFW recombines the top cyclopropane yielding mutations, DGMDW recombines the mutants which were the most selective for *cis*- formation, and DHMVW recombines the variants which had the highest objective function. Diastereoselectivities are shown above the plot as the trans:cis cyclopropane ratio.

### Biocatalytic Cyclopropanation of Olefins

**Substrates Utilized in Plate Screening**

**Table S7**. Table of styrenyl substrates investigated in this study.

| Substrate | Name | Structure |
|---|---|---|
| **1a** | 4-vinylanisole | 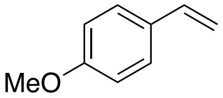 |
| **1b** | Styrene | 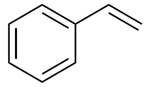 |
| **1c** | 1-methyl-4-vinylbenzene | 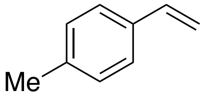 |
| **1d** | 1-methyl-3-vinylbenzene | 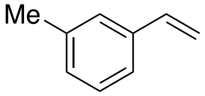 |
| **1e** | 1-methyl-2-vinylbenzene | 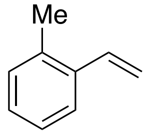 |
| **1f** | 1-chloro-4-vinylbenzene | 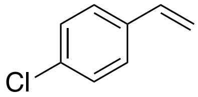 |
| **1g** | 1-bromo-4-vinylbenzene | 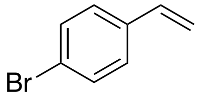 |
| **1h** | 1-(trifluoromethyl)-4-vinylbenzene | 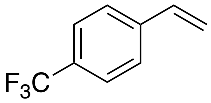 |
| **1i** | 2-vinylnaphthylene | 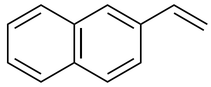 |

*Protocols for the Screening of Protoglobin Variants in 96-well Plate Format*

**96-Well Plate Library Expression:**

The wells of a 2-mL 96-well deep-well plate were filled with 400 µL LB-Amp. Previously generated 96-well plate glycerol stocks were removed from -80°C storage and placed on dry ice. Multichannel pipet tips were used to scratch the frozen glycerol stock surface and used to inoculate the aforementioned deep-well plate. These overnight cultures were incubated at 37 ºC and shaken at 220 rpm for 16–18 hours. For expression cultures, the following morning 50 µL of the precultures were used to inoculate 900 µL TB-amp per well in 96-well deep-well plates. The expression cultures were initially incubated at 37 ºC and 220 rpm for 2.5 hours, at which point they were allowed to sit at room temperature for 30 minutes. Expression of proteins was induced with IPTG and cellular heme production was increased with ALA. An induction mixture containing IPTG and ALA in TB-amp (50 µL) was added to each well such that the final concentrations of IPTG and ALA were 0.5 mM and 1.0 mM respectively. The total culture volumes were 1 mL per well. The plates were then incubated at 22 ºC and 220 rpm overnight.

**96-Well Plate Library Reactions and Screening:**

Expression cultures containing *E. coli* expressing hemoproteins of interest were centrifuged at 4,000 × g for 10 minutes at 4 ºC. The supernatant was discarded, and nitrogen-free M9 minimal media (M9-N, 380 µL) was added to each well. The pellets were resuspended in this media via shaking at room temperature for 30 minutes. The plates were then pumped into a vinyl Coy anaerobic chamber (0–30 ppm $O_2$). To each well were added 20 µL of a MeCN solution with 200 mM of the desired styrene substrate and 300 mM of ethyl diazoacetate (EDA). The final reaction volume was 400 µL, and the final concentrations of the desired styrene and EDA were 10 mM and 15 mM, respectively. The plates were then sealed carefully with a foil cover and shaken at room temperature for 16 hours in the Coy chamber. Once complete, plates are worked up for processing by adding 600 µL of a 1:1 solution of ethyl acetate:cyclohexane containing 1,3,5-trimethoxybenzene as an internal standard (1.0 mM concentration). A silicone sealing mat (AWSM1003S, ArcticWhite) was used to cover each of the plates, and the two layers were thoroughly mixed by rapid inversion of the plates. The plates were then centrifuged (5,000 × g for

5 minutes at room temperature) to separate the phases. Afterwards, a 200-μL aliquot of the organic layer was transferred to a GC vial insert in a GC vial, and the samples were assayed by GC-FID.

**Protoglobin Compilation Screening Data**

Initial reaction screening was performed on a compilation of protoglobin sequences. This compilation plate was primarily composed of mutants of the protoglobin from *Aeropernix pyrum*. Reactions were prepared according to the protocols for the screening of protoglobin variants in 96-well plate format (page S41). Cyclopropanation yields for formation of *cis*-**2a** and *trans*-**2a** are presented.

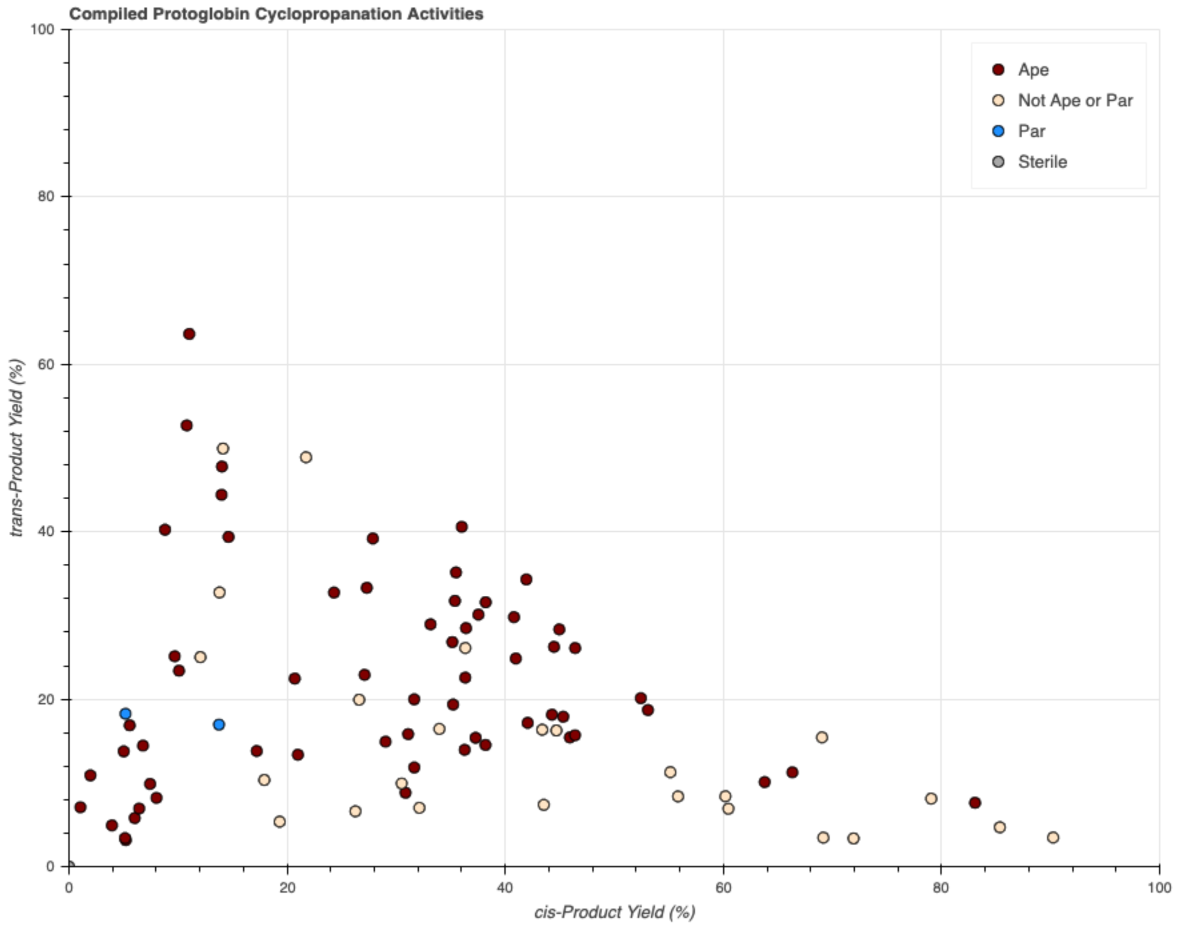


**Figure S31.** Yields of various protoglobin homologs and mutants for the formation of *cis*- and *trans*-**2a**.

#### ParLQ Single-Site Mutant Screening Data

Reactions of single-site mutants were prepared according to the protocols for the screening of protoglobin variants in 96-well plate format (page S41). Total cyclopropanation activity for formation of **2a**, *cis*-**2a**:*trans*-**2a** diastereoselectivity, and resulting objective function data are presented for reactions in triplicate.

#### Activity of W56X Mutants

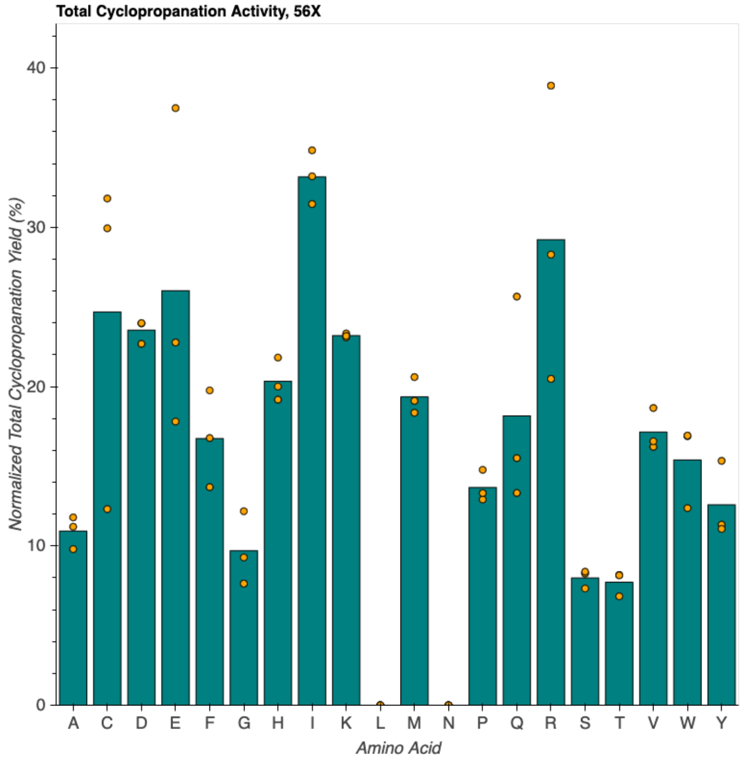


**Figure S32.** Total cyclopropanation activity of single-site mutants at position 56X. The original amino acid at this position was W. Amino acids L and Q were not observed in cloning and so were not screened.

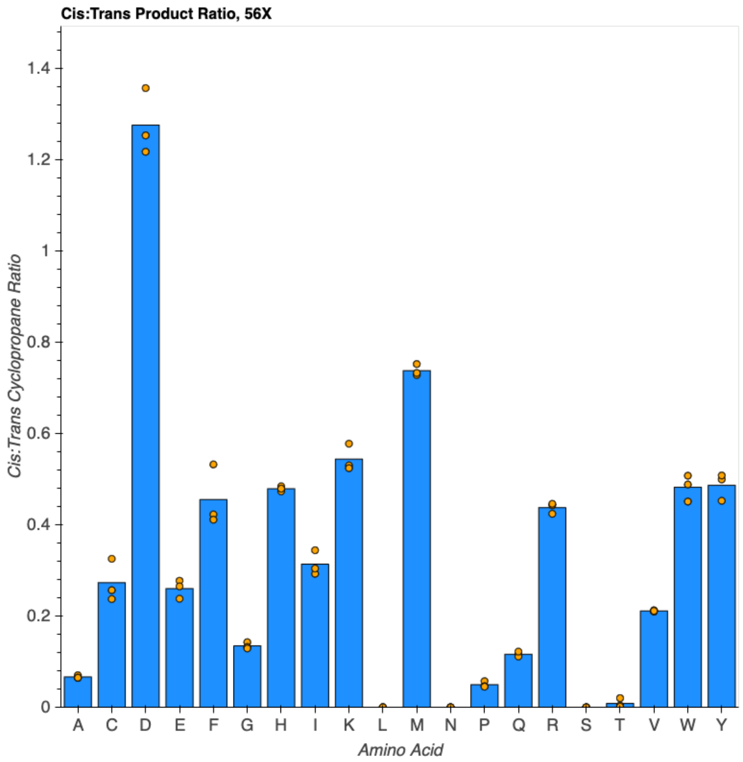


**Figure S33.** Diastereoselectivity of single-site mutants at position 56X. The original amino acid at this position was W. Amino acids L and Q were not observed in cloning and so were not screened.

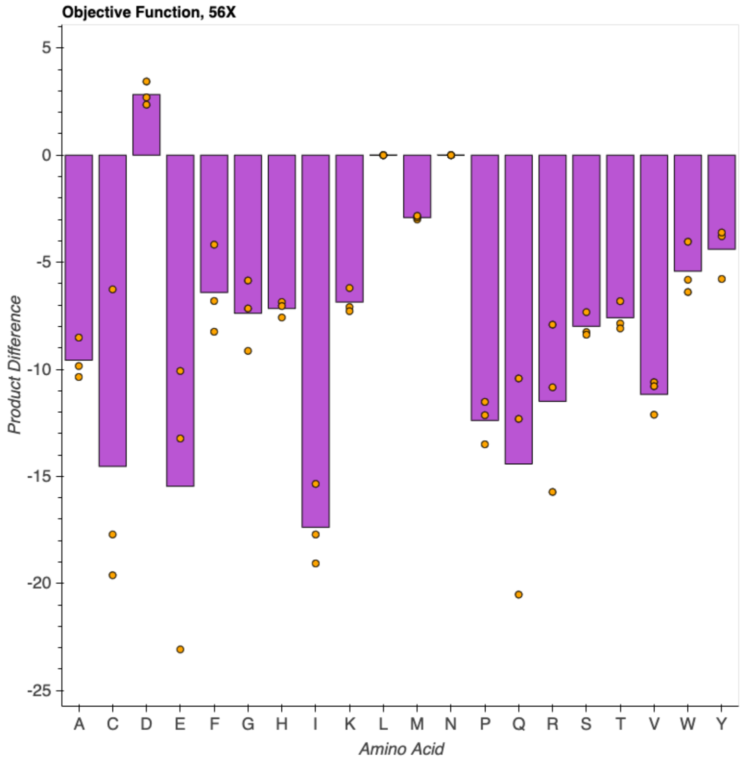


**Figure S34.** Objective function observed for single-site mutants at position 56X. The original amino acid at this position was W. Amino acids L and Q were not observed in cloning and so were not screened.

#### Activity of Y57X Mutants

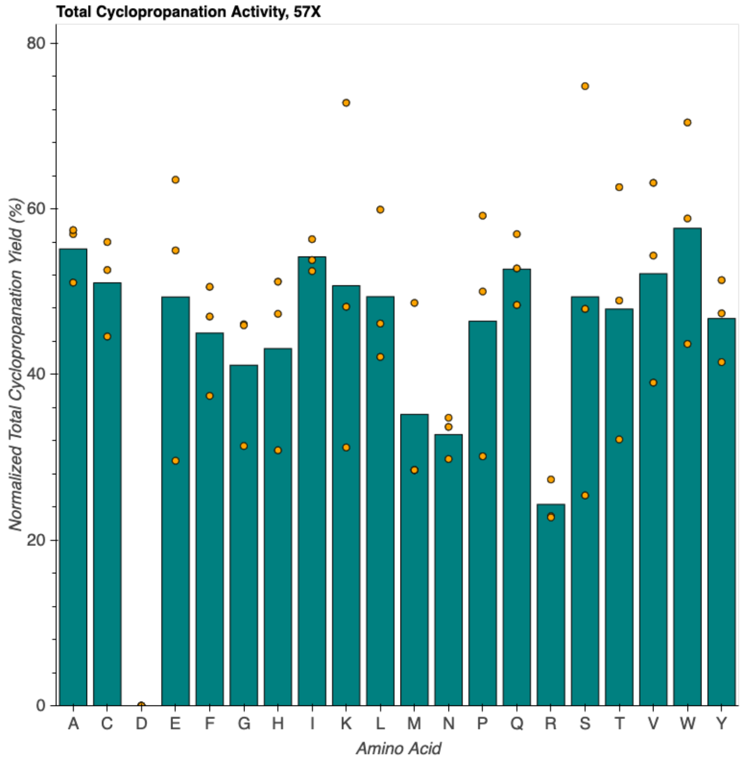


**Figure S35.** Total cyclopropanation activity of single-site mutants at position 57X. The original amino acid at this position was Y. Amino acid D was not observed in cloning and so was not screened.

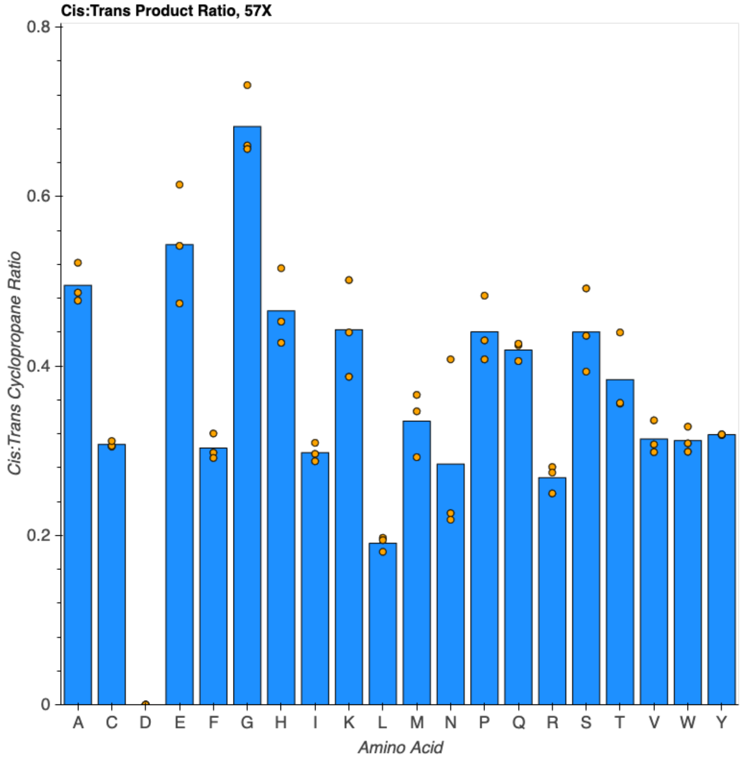


**Figure S36.** Diastereoselectivity of single-site mutants at position 57X. The original amino acid at this position was Y. Amino acid D was not observed in cloning and so was not screened.

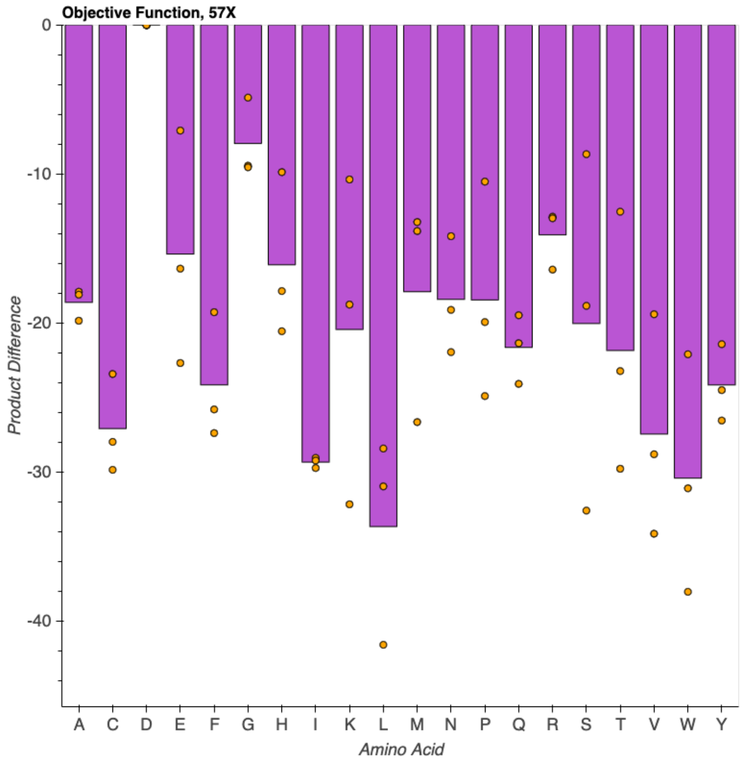


**Figure S37.** Objective function observed for single-site mutants at position 57X. The original amino acid at this position was Y. Amino acid D was not observed in cloning and so was not screened.

#### Activity of L59X Mutants

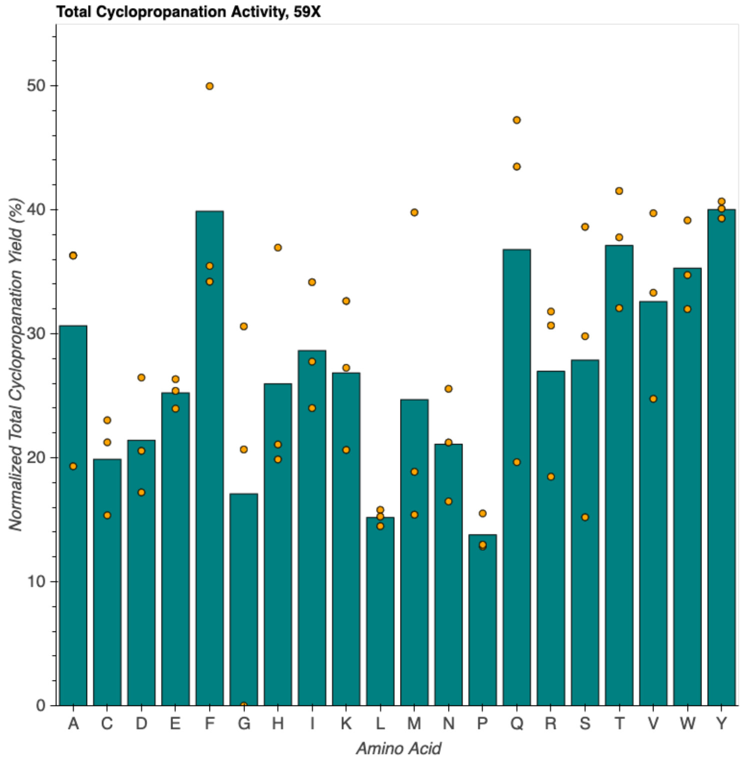


**Figure S38.** Total cyclopropanation activity of single-site mutants at position 59X. The original amino acid at this position was L.

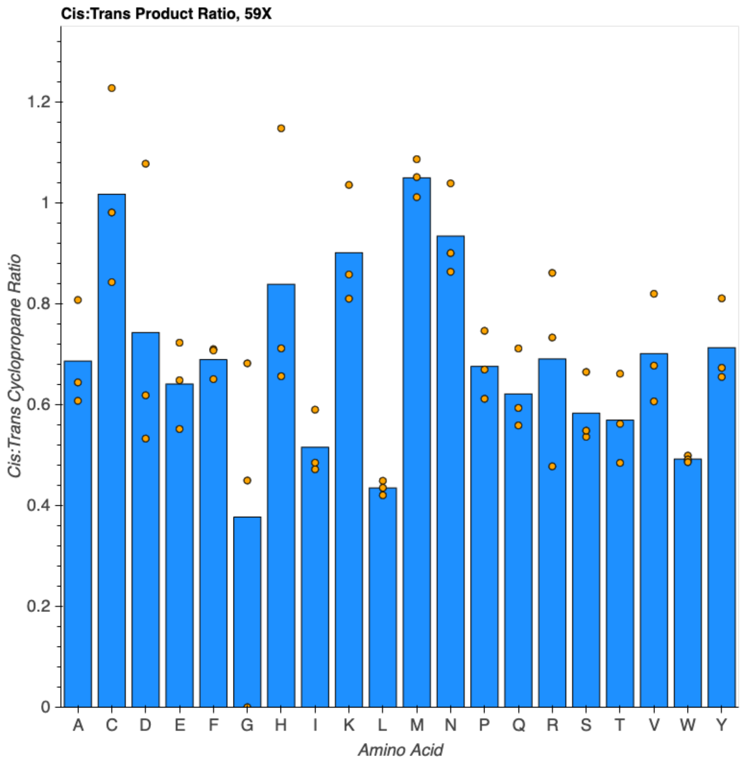


**Figure S39.** Diastereoselectivity of single-site mutants at position 59X. The original amino acid at this position was L.

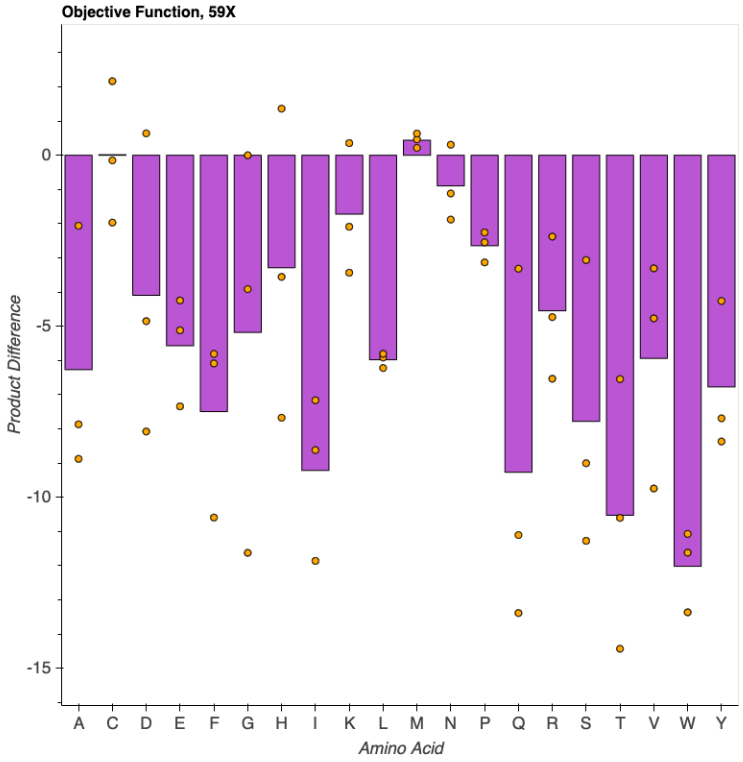


**Figure S40.** Objective function observed for single-site mutants at position 59X. The original amino acid at this position was L.

#### Activity of Q60X Mutants

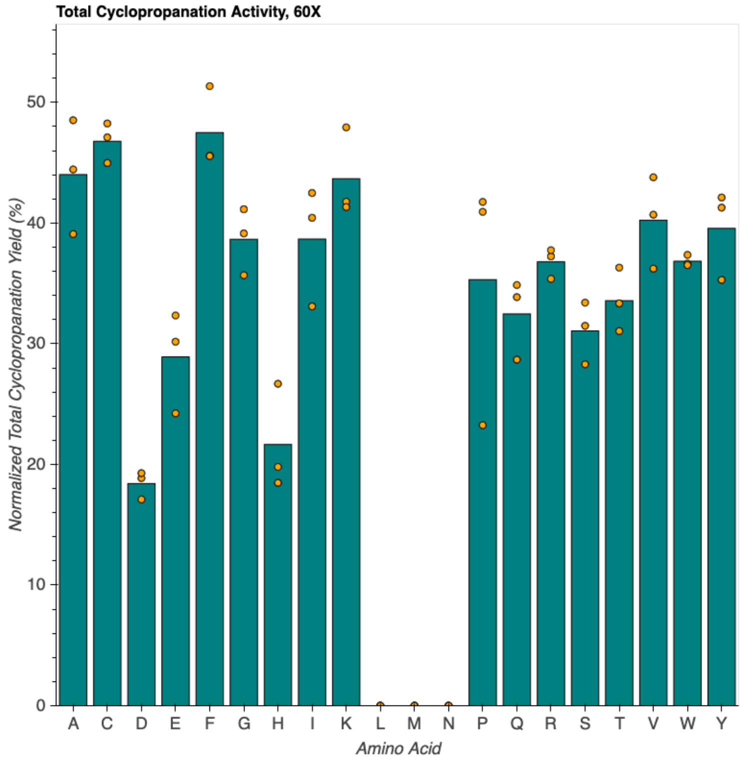


**Figure S41.** Total cyclopropanation activity of single-site mutants at position 60X. The original amino acid at this position was Q. Amino acids L, M, and N were not observed in cloning and so were not screened.

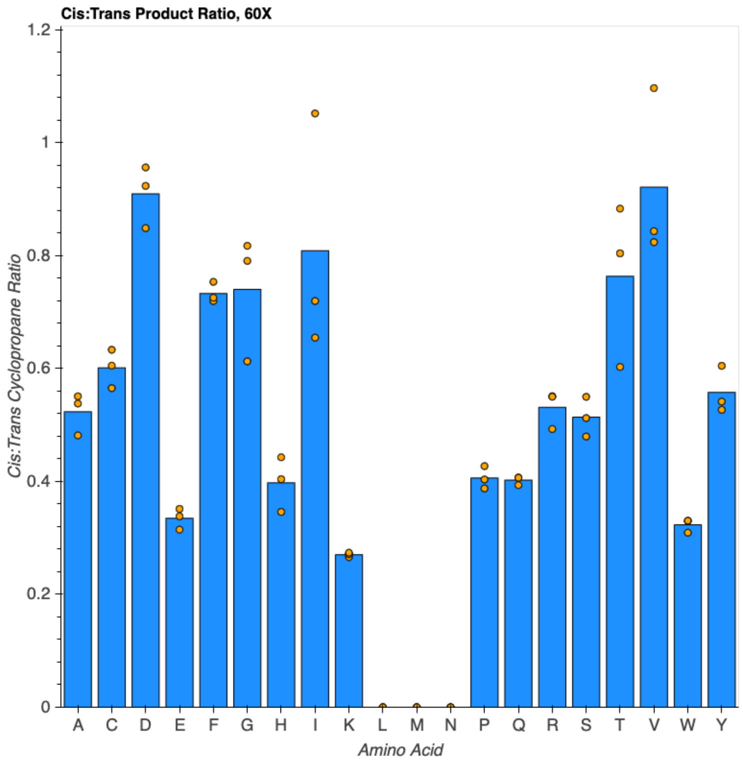


**Figure S42.** Diastereoselectivity of single-site mutants at position 60X. The original amino acid at this position was Q. Amino acids L, M, and N were not observed in cloning and so were not screened.

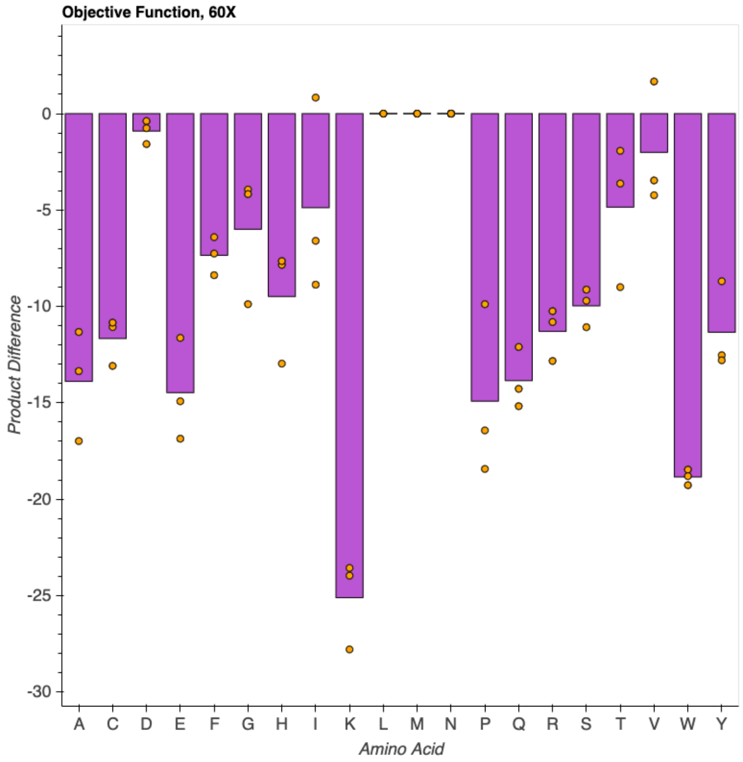


**Figure S43.** Objective function observed for single-site mutants at position 60X. The original amino acid at this position was Q. Amino acids L, M, and N were not observed in cloning and so were not screened.

#### Activity of F89X Mutants

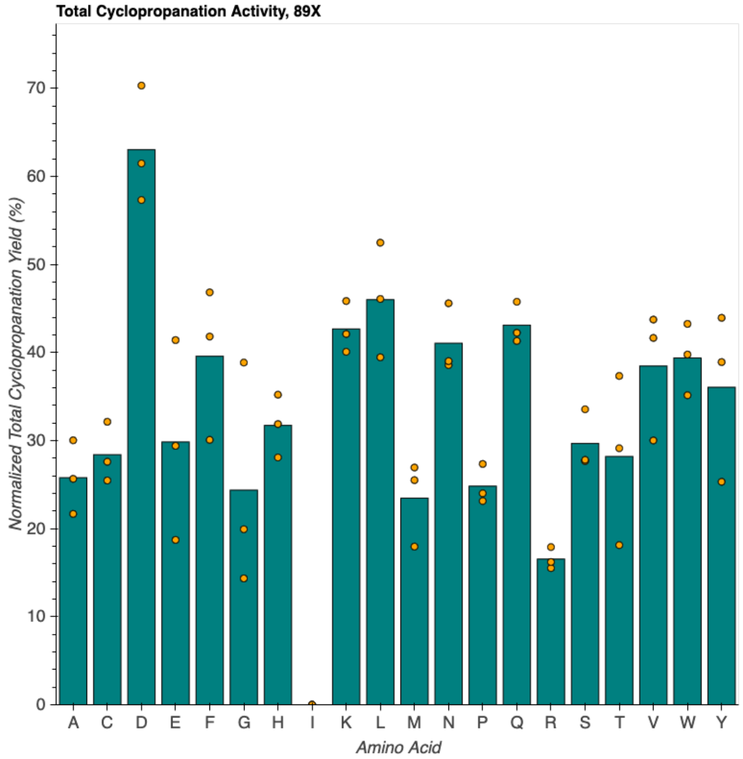


**Figure S44.** Total cyclopropanation activity of single-site mutants at position 89X. The original amino acid at this position was Q. Amino acid I was not observed in cloning and so was not screened.

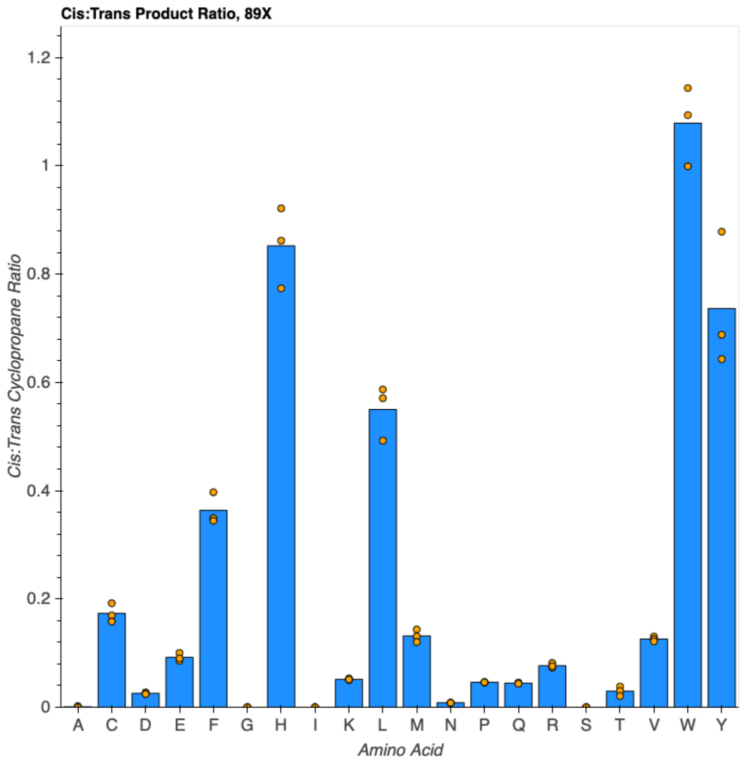


**Figure S45.** Diastereoselectivity of single-site mutants at position 89X. The original amino acid at this position was Q. Amino acid I was not observed in cloning and so was not screened.

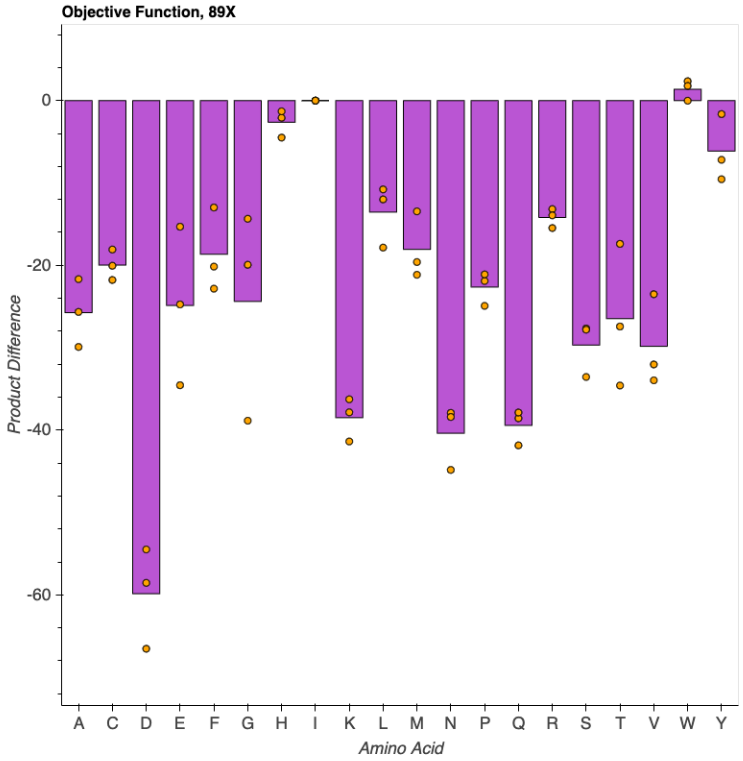


**Figure S46.** Total cyclopropanation activity of single-site mutants at position 89X. The original amino acid at this position was Q. Amino acid I was not observed in cloning and so was not screened.

**ParLQ 5-site Multi-Mutant Screening Data with the Model Substrate *1a***

Reactions were prepared according to the protocols for the screening of protoglobin variants in 96-well plate format (page S41). Total cyclopropanation for formation of *cis*-**2a** and *trans*-**2a** are presented.

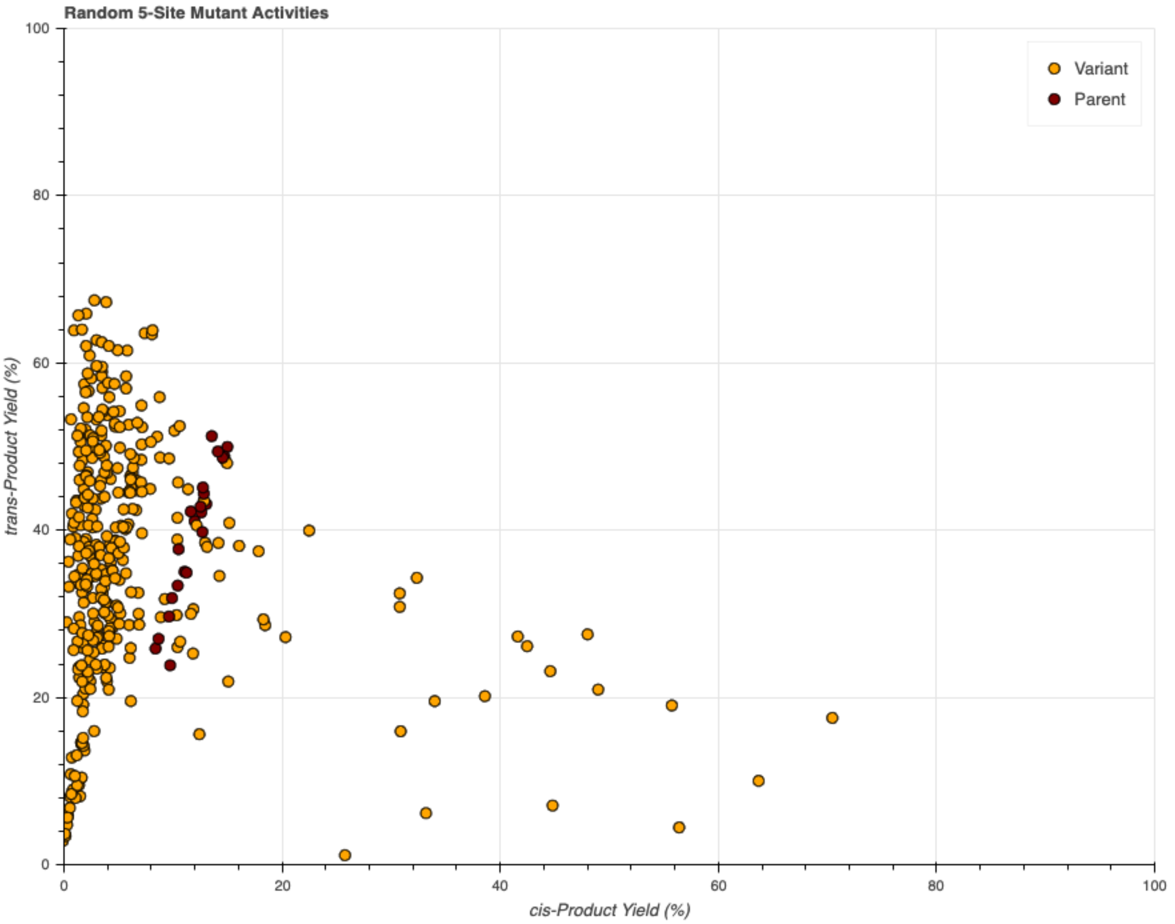


**Figure S47.** Yields of various random ParLQ 5-site mutants for the formation of *cis*- and *trans*-**2a**. Only ~10% of variants in this random library had improved selectivity for *cis*-**2a.**

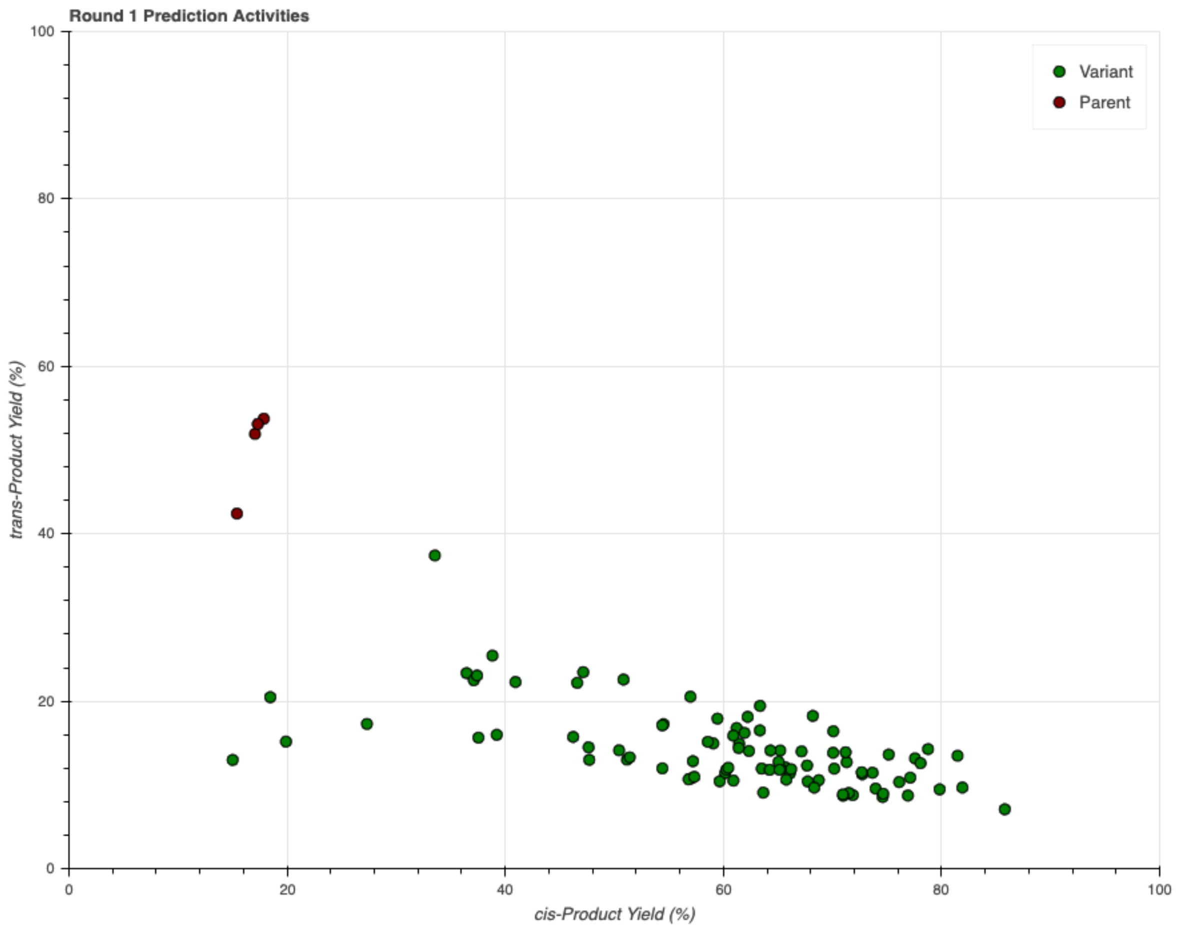


**Figure S48.** Activities of the first round of mutants predicted by ALDE. Reactions with this library of variants were run in triplicate, and the average yields for the formation of *cis*- and *trans*-**2a** observed for each variant are shown.

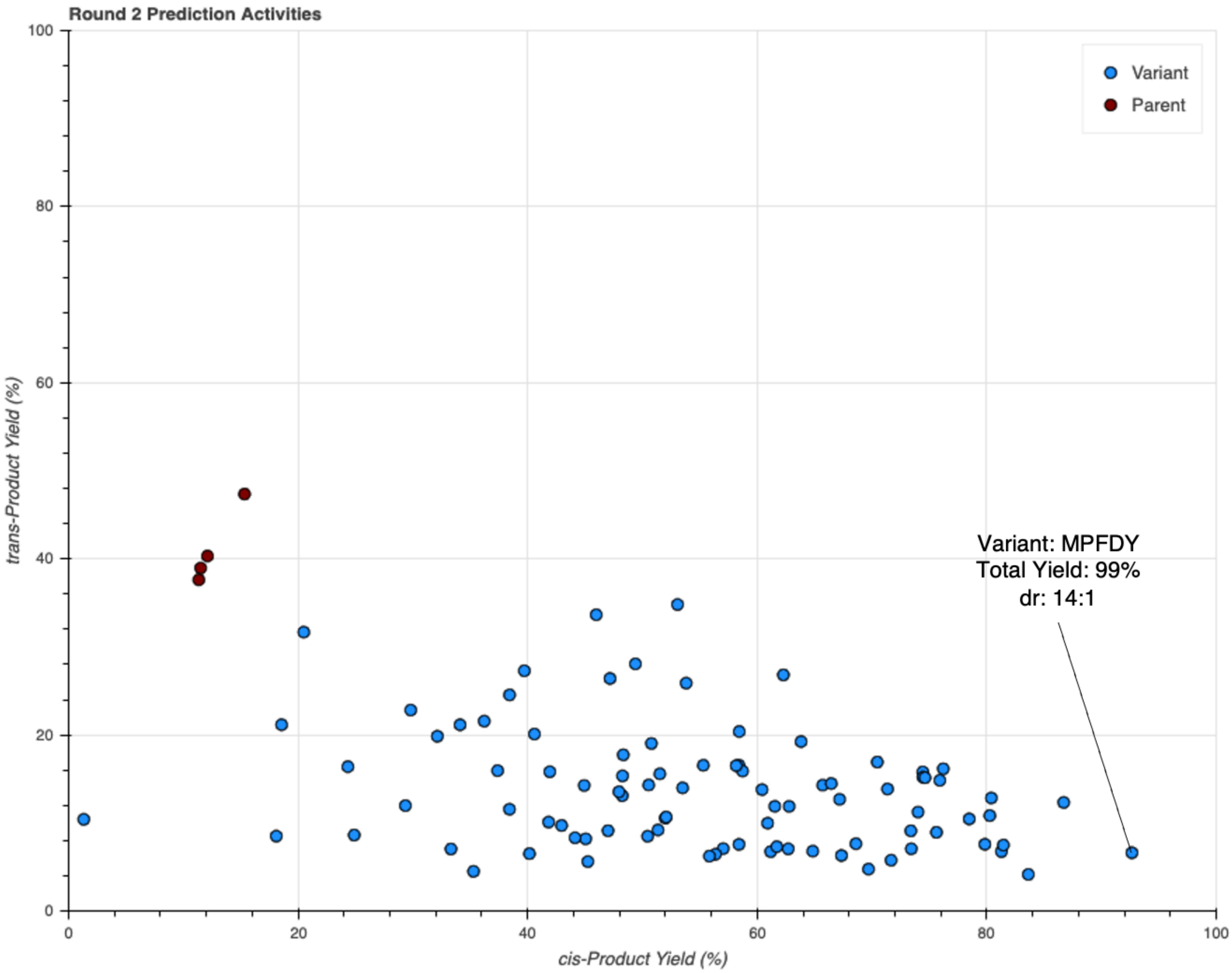


**Figure S49.** Activities of the second round of mutants predicted by ALDE. Reactions with this library of variants were run in triplicate, and the average yields for the formation of *cis*- and *trans*-**2a** observed for each variant are shown. The highest-performing variant, MPFDY, demonstrated a yield of 99% with a 14:1 selectivity for *cis*-**2a**.

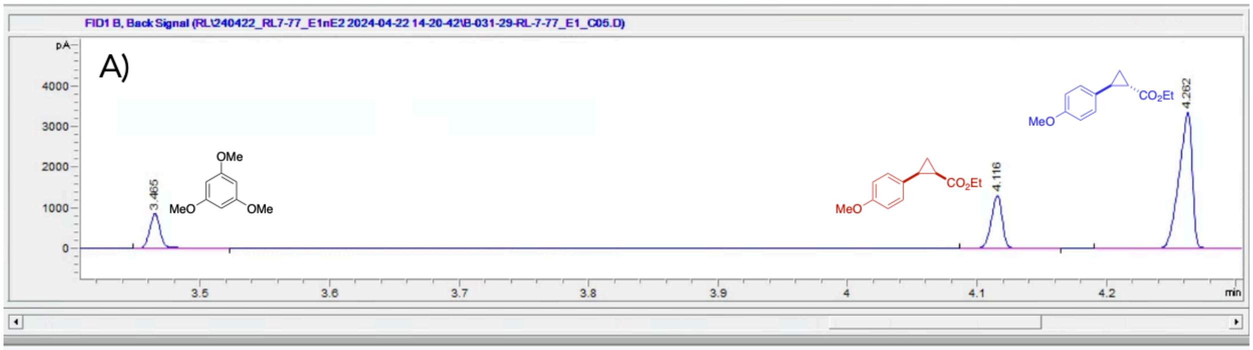


| # | Time | Type | Area | Height | Width | Area% | Symmetry |
|---|---|---|---|---|---|---|---|
| 1 | 3.465 | BB | 482.3 | 827.1 | 9.2036E-3 | 13.332 | 0.864 |
| 2 | 4.116 | BB | 758.2 | 1270.9 | 9.3604E-3 | 20.958 | 1.074 |
| 3 | 4.262 | VV R | 2377.3 | 3317.8 | 0.0115 | 65.710 | 1.488 |

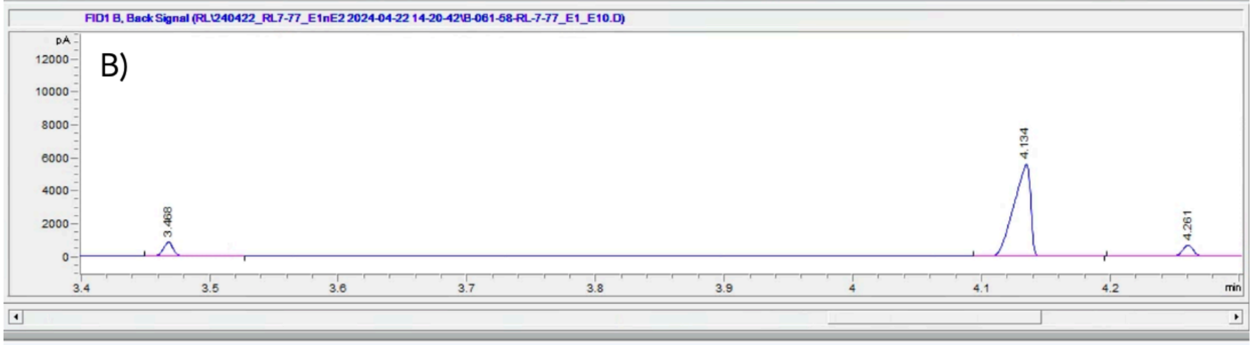


| # | Time | Type | Area | Height | Width | Area% | Symmetry |
|---|---|---|---|---|---|---|---|
| 1 | 3.468 | BB | 488.8 | 837.1 | 9.2129E-3 | 8.206 | 0.876 |
| 2 | 4.134 | BB | 5044.5 | 5634.2 | 0.0144 | 84.688 | 1.996 |
| 3 | 4.261 | VV R | 423.3 | 661.6 | 9.689E-3 | 7.106 | 0.968 |

**Figure S50.** A) Representative GC-FID trace for reactions of ParLQ with **1a** and EDA. B) GC-FID trace for the top-performing predicted variant, MPFDY, in reactions with **1a** and EDA.

**Reactions of ALDE Round 2 Predictions with Non-Model Substrates**

Reactions were prepared according to the protocols for the screening of protoglobin variants in 96-well plate format (page S41). Total cyclopropanation for the formation of the *cis*- and *trans*-stereoisomers of each substrate are presented.

*Substrate 1b – Styrene*

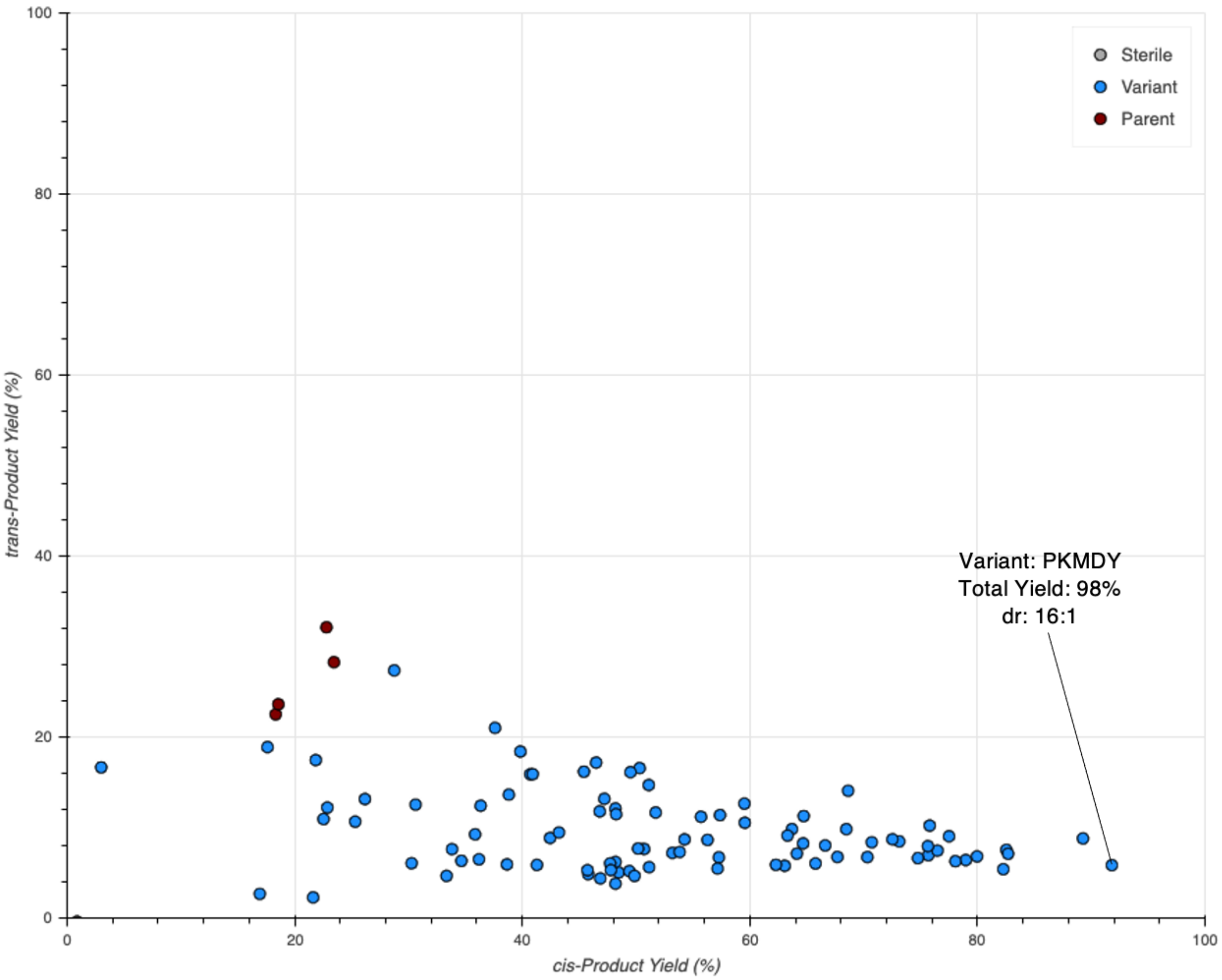


**Figure S51.** Activities of the second round of mutants predicted by ALDE with substrate **1b**. Observed yields for the formation of *cis*- and *trans*-**2b** are shown. The highest-performing variant, PKMDY, demonstrated a yield of 98% with a 16:1 selectivity for *cis*-**2b**.

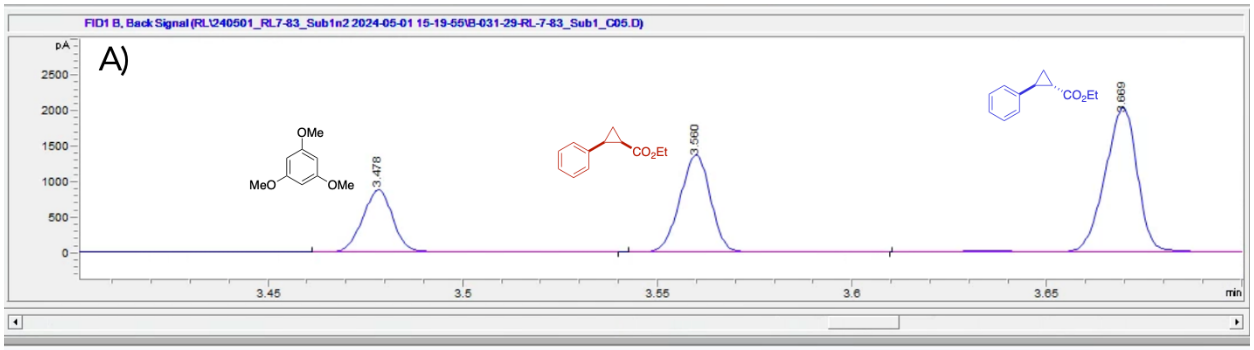


| # | Time | Type | Area | Height | Width | Area% | Symmetry |
|---|---|---|---|---|---|---|---|
| 1 | 3.478 | BB | 484.7 | 854.5 | 9.0178E-3 | 19.803 | 0.934 |
| 2 | 3.56 | BB | 752.7 | 1299.2 | 9.1591E-3 | 30.750 | 1.002 |
| 3 | 3.669 | VV R | 1210.3 | 1903.9 | 0.0105 | 49.448 | 1.109 |

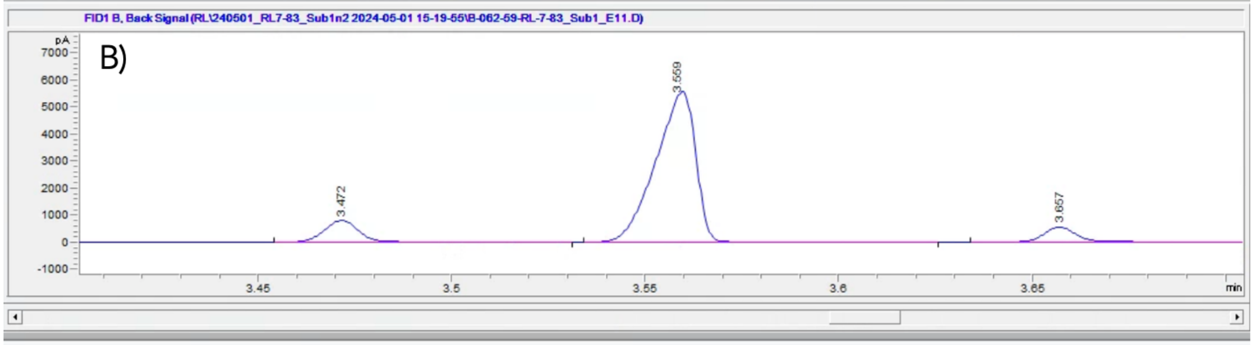


| # | Time | Type | Area | Height | Width | Area% | Symmetry |
|---|---|---|---|---|---|---|---|
| 1 | 3.472 | BB | 490.4 | 768.6 | 0.0106 | 10.259 | 0.899 |
| 2 | 3.559 | BB | 3963.3 | 5332.3 | 0.0111 | 82.908 | 1.429 |
| 3 | 3.657 | BB | 326.6 | 536.3 | 9.5048E-3 | 6.833 | 0.793 |

**Figure S52.** A) Representative GC-FID trace for reactions of ParLQ with **1b** and EDA. B) GC-FID trace for the top-performing variant, PKMDY, in reactions with **1b** and EDA.

*Substrate 1c - 1-methyl-4-vinylbenzene*

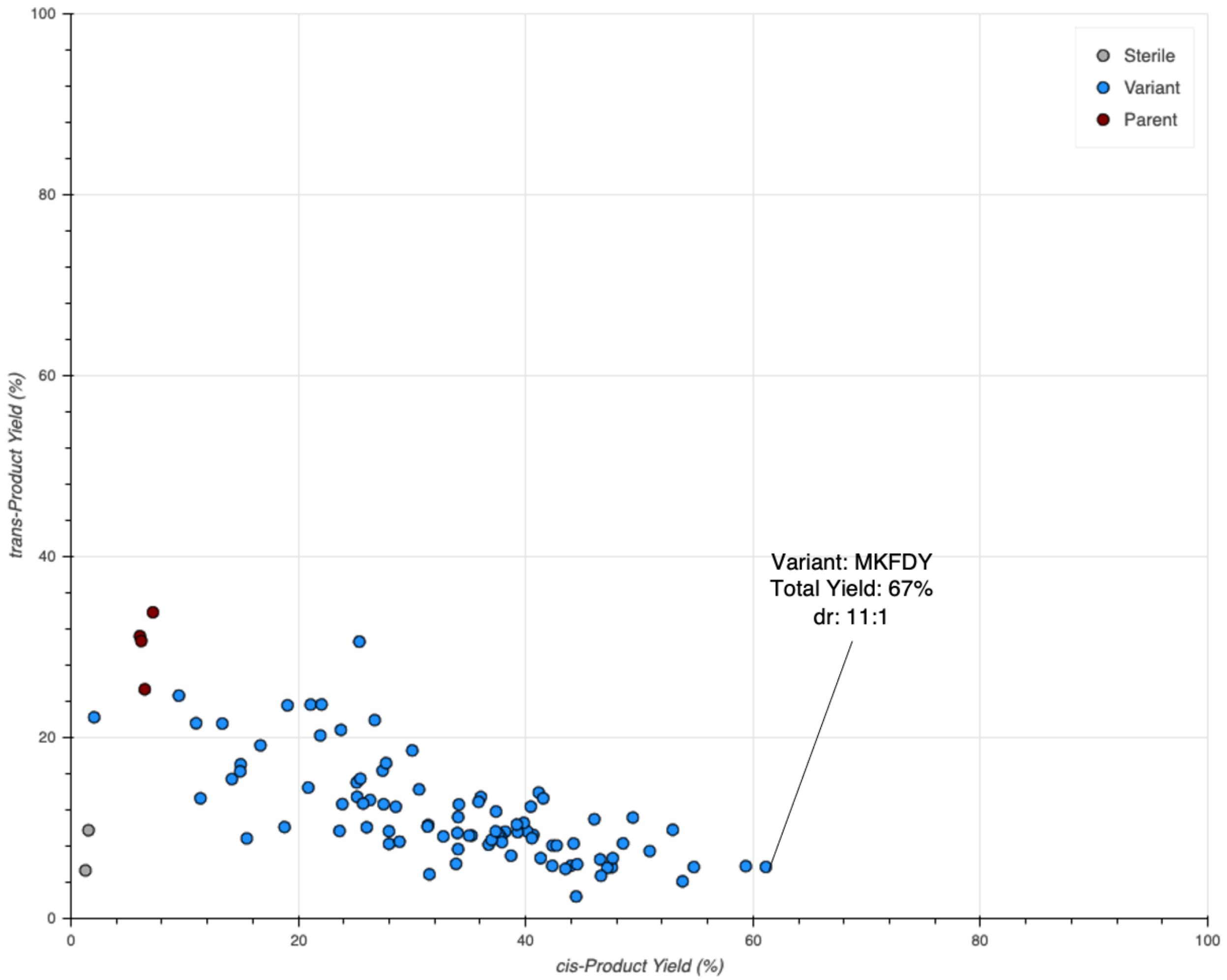


**Figure S53.** Activities of the second round of mutants predicted by ALDE with substrate **1c**. Observed yields for the formation of *cis*- and *trans*-**2c** are shown. The highest-performing variant, MKFDY, demonstrated a yield of 67% with a 11:1 selectivity for *cis*-**2c**.

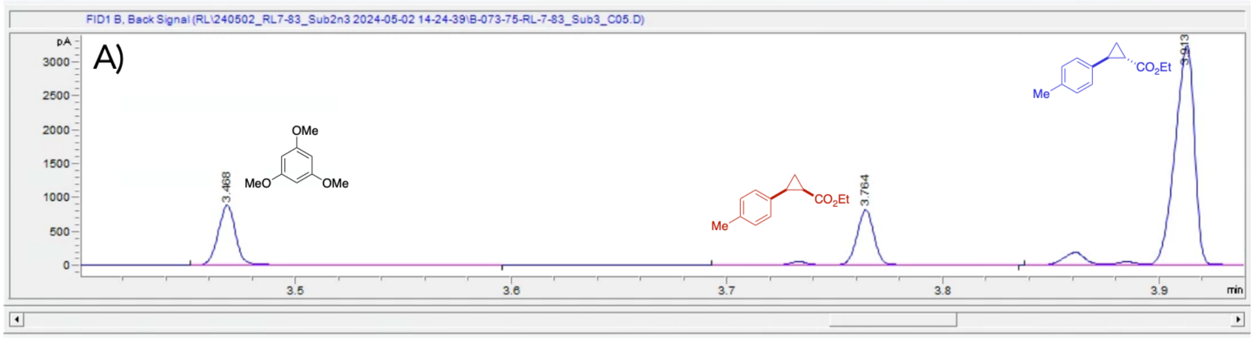


| # | Time | Type | Area | Height | Width | Area% | Symmetry |
|---|---|---|---|---|---|---|---|
| 1 | 3.468 | BV R | 503.4 | 857.2 | 9.2322E-3 | 15.859 | 0.894 |
| 2 | 3.764 | VB R | 479.2 | 790.9 | 9.0053E-3 | 15.097 | 1.325 |
| 3 | 3.913 | VB R | 2191.7 | 3206.6 | 0.0106 | 69.044 | 1.786 |

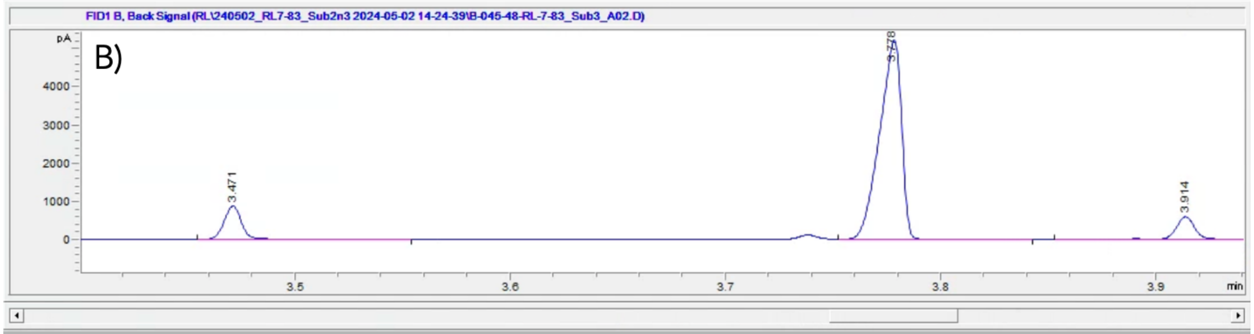


| # | Time | Type | Area | Height | Width | Area% | Symmetry |
|---|---|---|---|---|---|---|---|
| 1 | 3.471 | BB | 492.8 | 849.4 | 9.1693E-3 | 10.845 | 0.889 |
| 2 | 3.778 | VB | 3694 | 5026.7 | 0.011 | 81.289 | 1.434 |
| 3 | 3.914 | VB R | 357.5 | 575.6 | 9.5082E-3 | 7.866 | 1.063 |

**Figure S54.** A) Representative GC-FID trace for reactions of ParLQ with **1c** and EDA. B) GC-FID trace for the top-performing variant, MKFDY, in reactions with **1c** and EDA.

*Substrate 1d - 1-methyl-3-vinylbenzene*

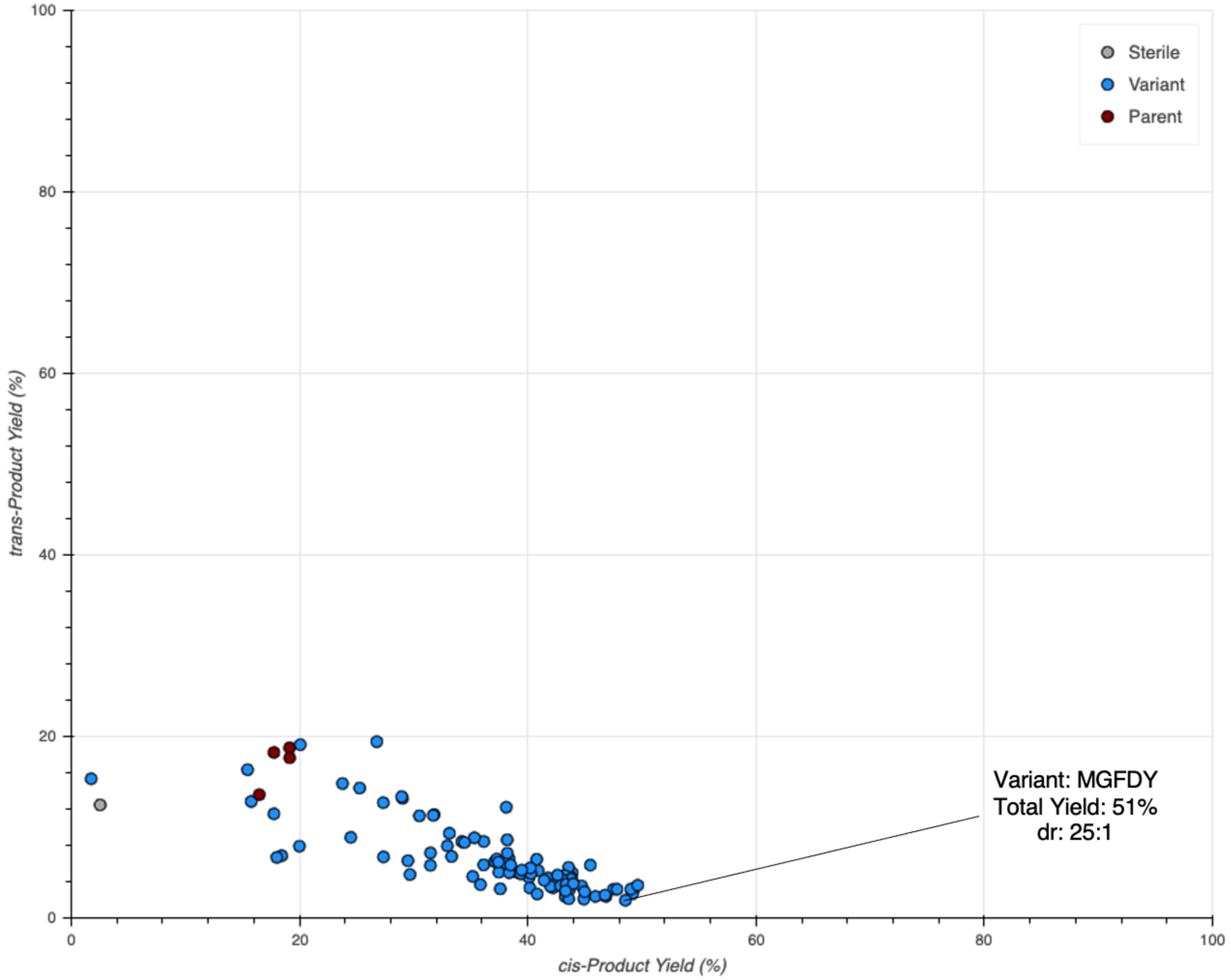


**Figure S55.** Activities of the second round of mutants predicted by ALDE with substrate **1d**. Observed yields for the formation of *cis*- and *trans*-**2d** are shown. The highest-performing variant, MGFDY, demonstrated a yield of 51% with a 25:1 selectivity for *cis*-**2d**.

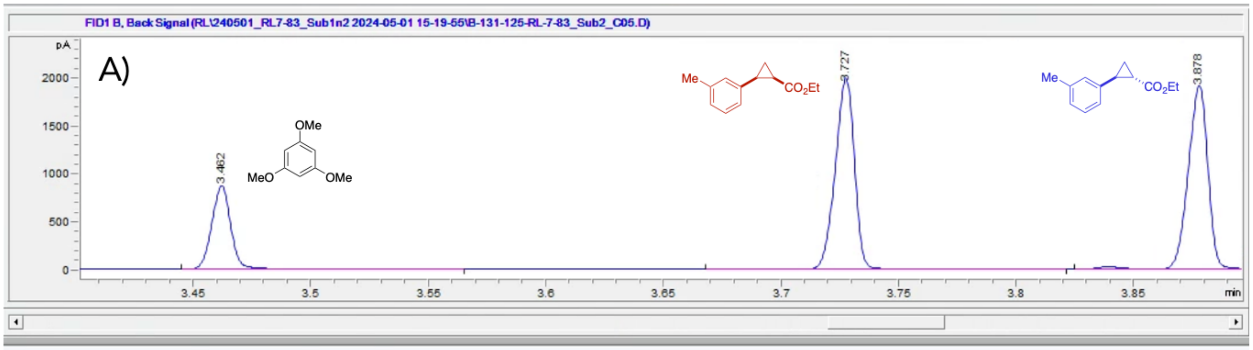


| # | Time | Type | Area | Height | Width | Area% | Symmetry |
|---|---|---|---|---|---|---|---|
| 1 | 3.462 | BB | 492.4 | 846.4 | 9.1873E-3 | 17.903 | 0.897 |
| 2 | 3.727 | VB R | 1133.7 | 1906.2 | 9.3229E-3 | 41.218 | 1.062 |
| 3 | 3.878 | VB R | 1124.4 | 1874.6 | 0.0101 | 40.879 | 1.169 |

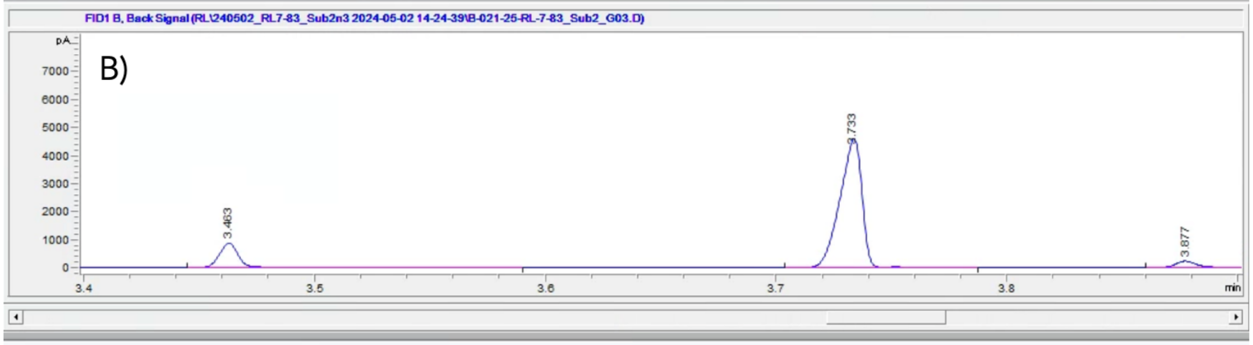


| # | Time | Type | Area | Height | Width | Area% | Symmetry |
|---|---|---|---|---|---|---|---|
| 1 | 3.463 | BV R | 491.7 | 843.4 | 9.1853E-3 | 13.379 | 0.882 |
| 2 | 3.733 | BB | 3042.7 | 4234 | 0.0116 | 82.799 | 1.325 |
| 3 | 3.877 | BB | 140.4 | 214.3 | 0.0108 | 3.822 | 0.769 |

**Figure S56.** A) Representative GC-FID trace for reactions of ParLQ with **1d** and EDA. B) GC-FID trace for the top-performing variant, MGFDY, in reactions with **1d** and EDA.

*Substrate 1e - 1-methyl-2-vinylbenzene*

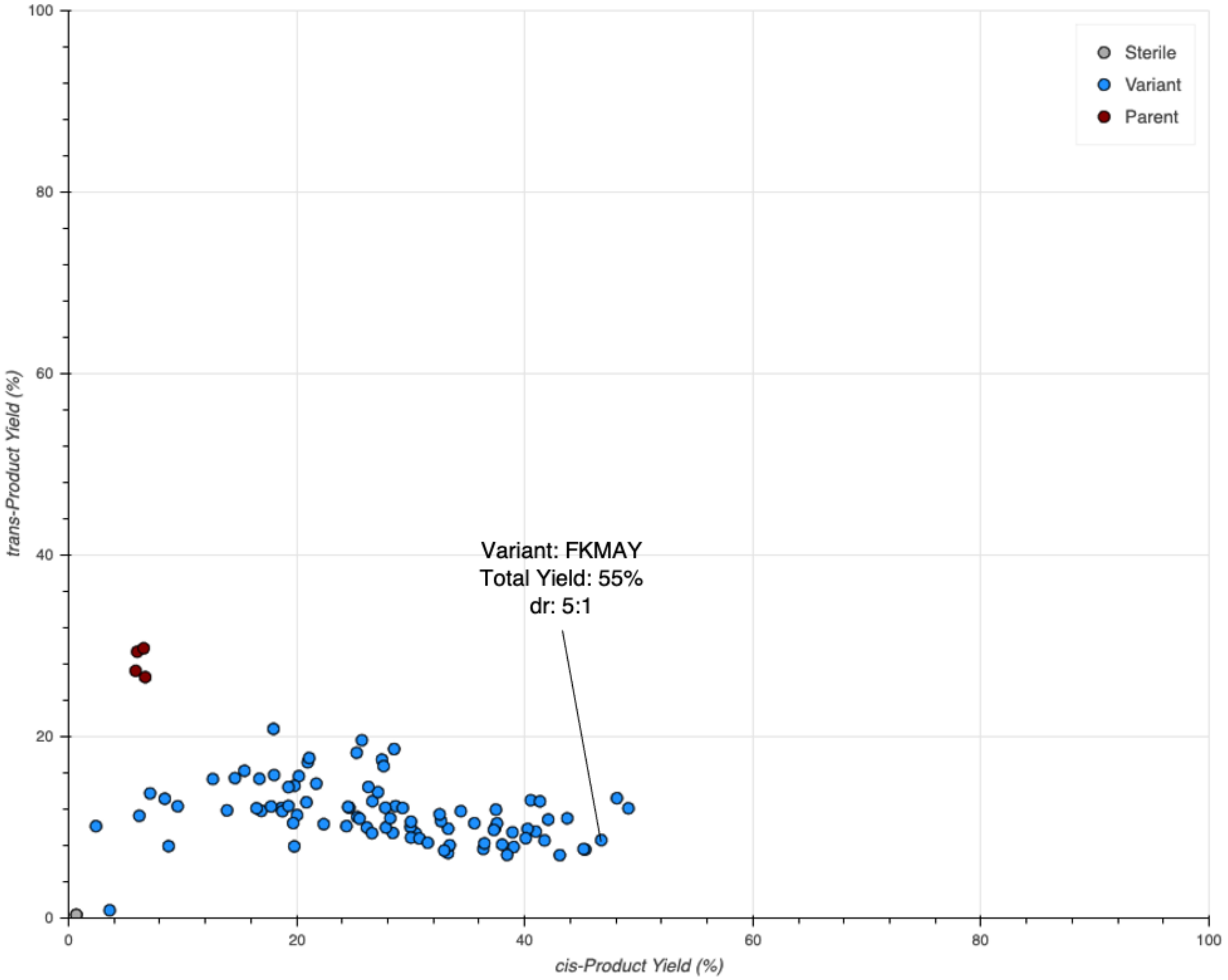


**Figure S57.** Activities of the second round of mutants predicted by ALDE with substrate **1e**. Observed yields for the formation of *cis*- and *trans*-**2e** are shown. The highest-performing variant, FKMAY, demonstrated a yield of 55% with a 5:1 selectivity for *cis*-**2e**.

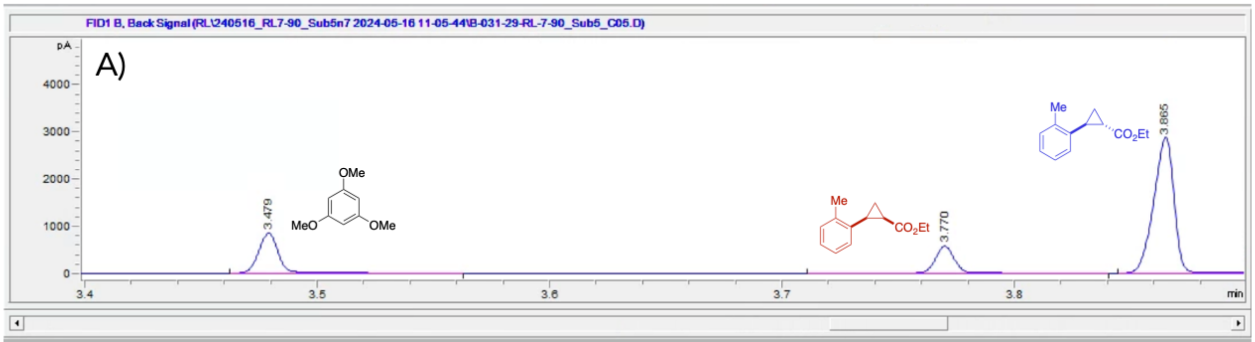


| # | Time | Type | Area | Height | Width | Area% | Symmetry |
|---|---|---|---|---|---|---|---|
| 1 | 3.479 | BB | 493.3 | 823.8 | 9.3867E-3 | 19.229 | 0.882 |
| 2 | 3.77 | VB R | 322.5 | 550.2 | 9.2388E-3 | 12.571 | 0.932 |
| 3 | 3.865 | BB | 1749.8 | 2835.2 | 9.6001E-3 | 68.201 | 1.131 |

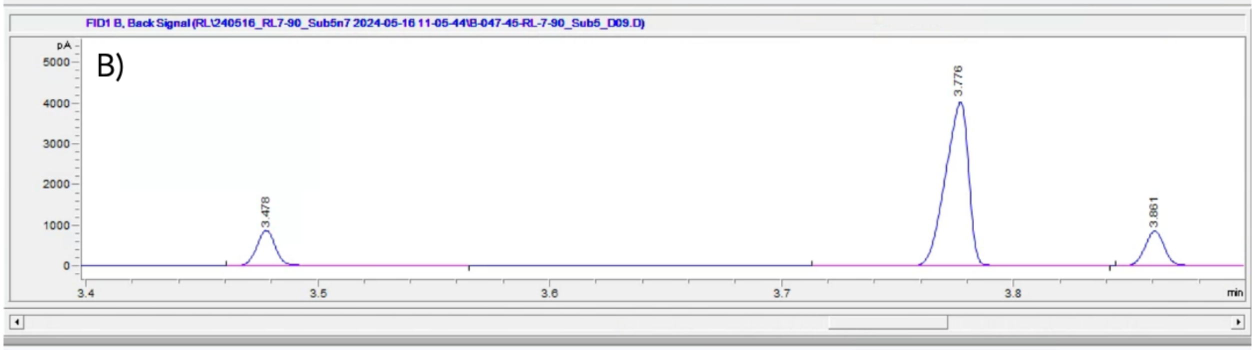


| # | Time | Type | Area | Height | Width | Area% | Symmetry |
|---|---|---|---|---|---|---|---|
| 1 | 3.478 | BB | 491.8 | 831.8 | 0.0101 | 13.051 | 0.884 |
| 2 | 3.776 | BB | 2781.1 | 4032.1 | 0.0112 | 73.799 | 1.345 |
| 3 | 3.861 | BB | 495.6 | 822.9 | 9.4255E-3 | 13.150 | 0.928 |

**Figure S58.** A) Representative GC-FID trace for reactions of ParLQ with **1e** and EDA. B) GC-FID trace for the top-performing variant, FKMAY, in reactions with **1e** and EDA.

*Substrate 1f - 1-chloro-4-vinylbenzene*

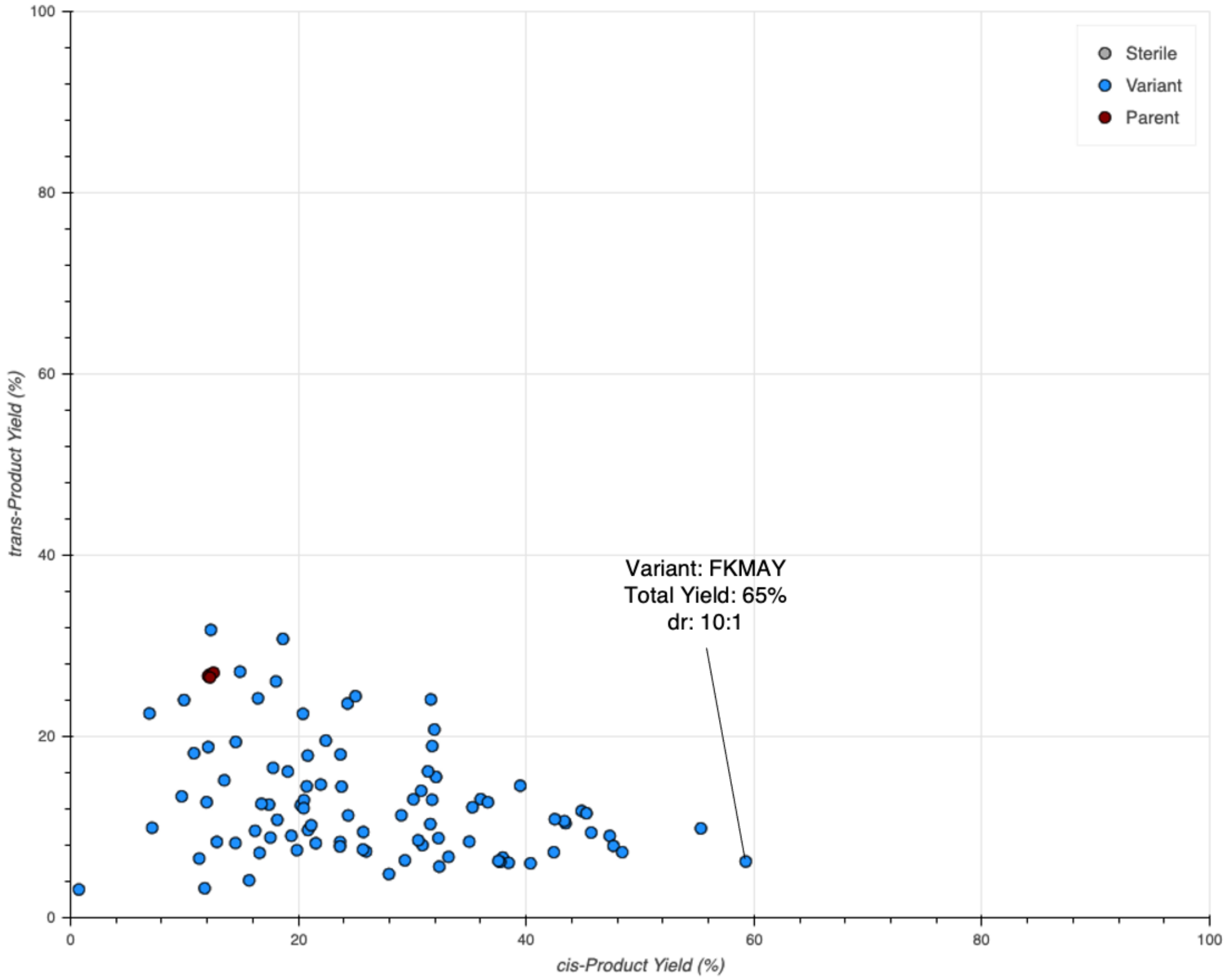


**Figure S59.** Activities of the second round of mutants predicted by ALDE with substrate **1f**. Observed yields for the formation of *cis*- and *trans*-**2f** are shown. The highest-performing variant, FKMAY, demonstrated a yield of 65% with a 10:1 selectivity for *cis*-**2f**.

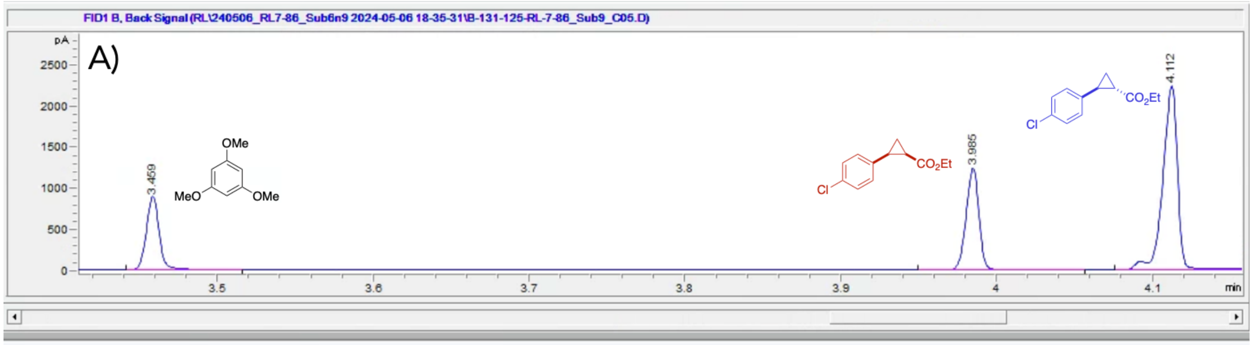


| # | Time | Type | Area | Height | Width | Area% | Symmetry |
|---|---|---|---|---|---|---|---|
| 1 | 3.459 | BB | 518.7 | 857.3 | 0.0103 | 18.794 | 0.855 |
| 2 | 3.985 | BB | 711.6 | 1215.5 | 9.2306E-3 | 25.783 | 1.016 |
| 3 | 4.112 | BV R | 1529.7 | 2207.6 | 0.0112 | 55.422 | 1.175 |

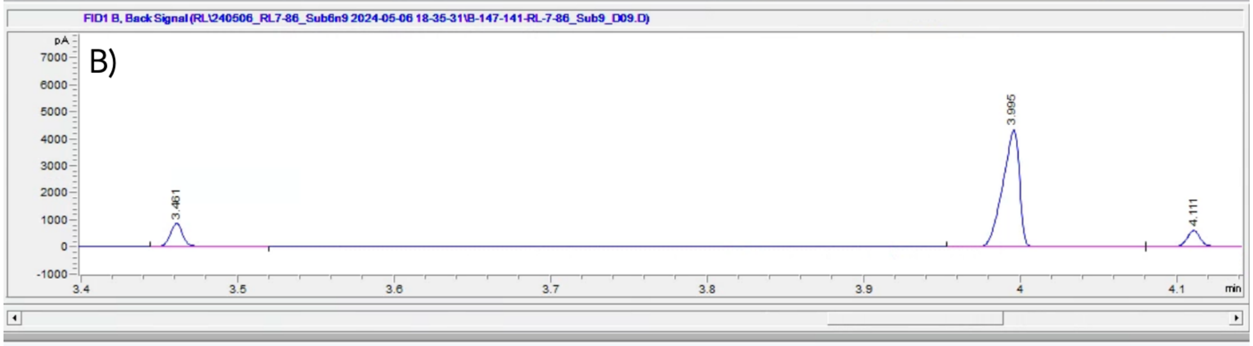


| # | Time | Type | Area | Height | Width | Area% | Symmetry |
|---|---|---|---|---|---|---|---|
| 1 | 3.461 | BB | 491 | 818.4 | 0.0102 | 12.153 | 0.839 |
| 2 | 3.995 | BB | 3194 | 4295.8 | 0.0119 | 79.054 | 1.496 |
| 3 | 4.111 | BB | 355.2 | 581.2 | 9.5304E-3 | 8.792 | 0.886 |

**Figure S60.** A) Representative GC-FID trace for reactions of ParLQ with **1f** and EDA. B) GC-FID trace for the top-performing variant, FKMAY, in reactions with **1f** and EDA.

*Substrate 1g - 1-bromo-4-vinylbenzene*

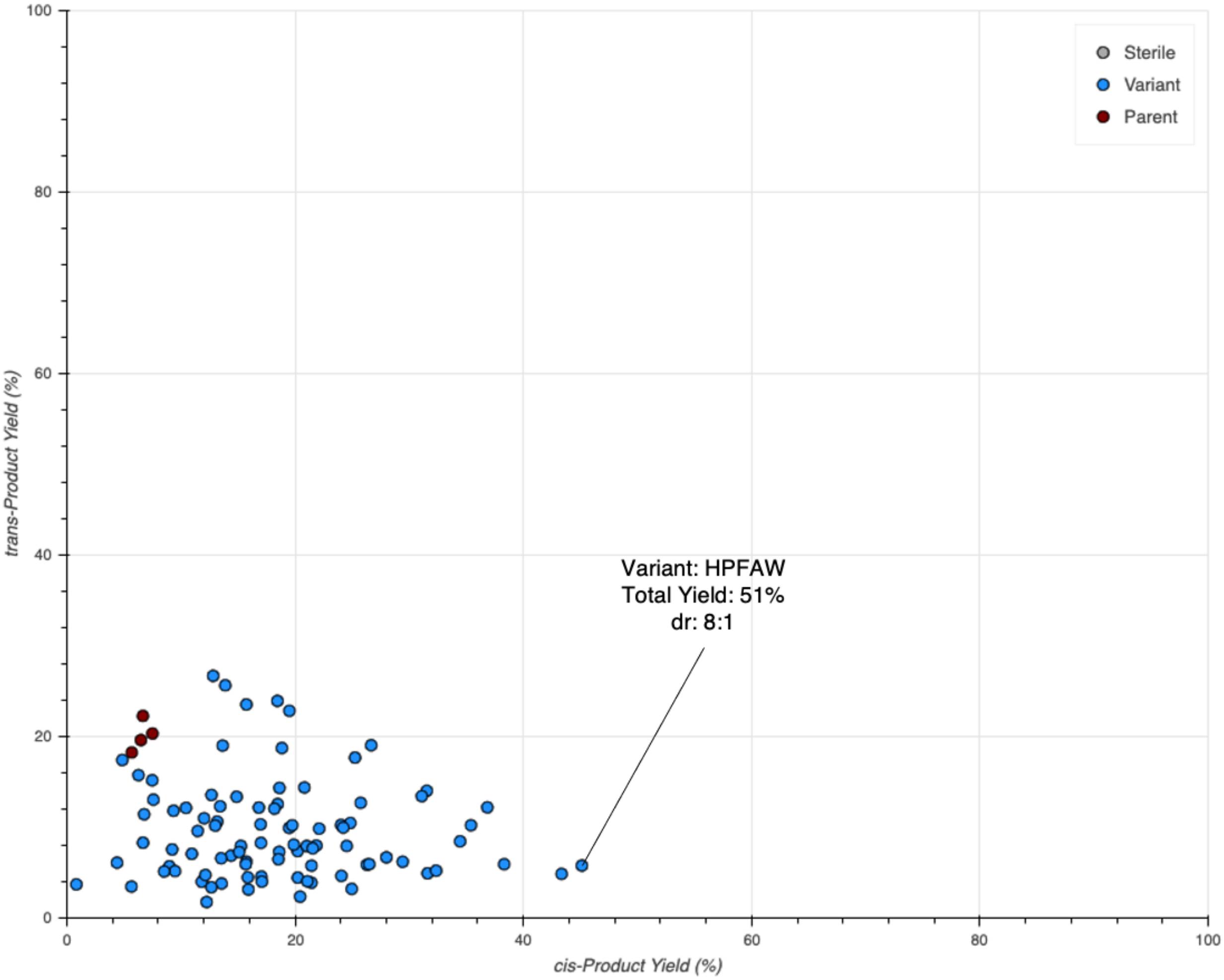


**Figure S61.** Activities of the second round of mutants predicted by ALDE with substrate **1g**. Observed yields for the formation of *cis*- and *trans*-**2g** are shown. The highest-performing variant, HPFAW, demonstrated a yield of 51% with a 8:1 selectivity for *cis*-**2g**.

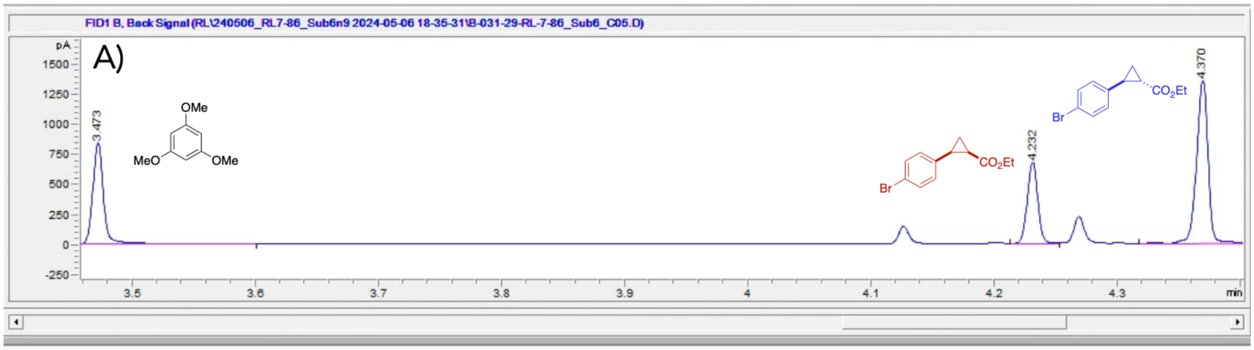


| # | Time | Type | Area | Height | Width | Area% | Symmetry |
|---|---|---|---|---|---|---|---|
| 1 | 3.473 | BV R | 507.7 | 814.4 | 0.0105 | 27.889 | 0.82 |
| 2 | 4.232 | BV | 404.4 | 654.9 | 0.0104 | 22.214 | 0.928 |
| 3 | 4.37 | VB R | 908.4 | 1329.6 | 0.0103 | 49.897 | 1.035 |

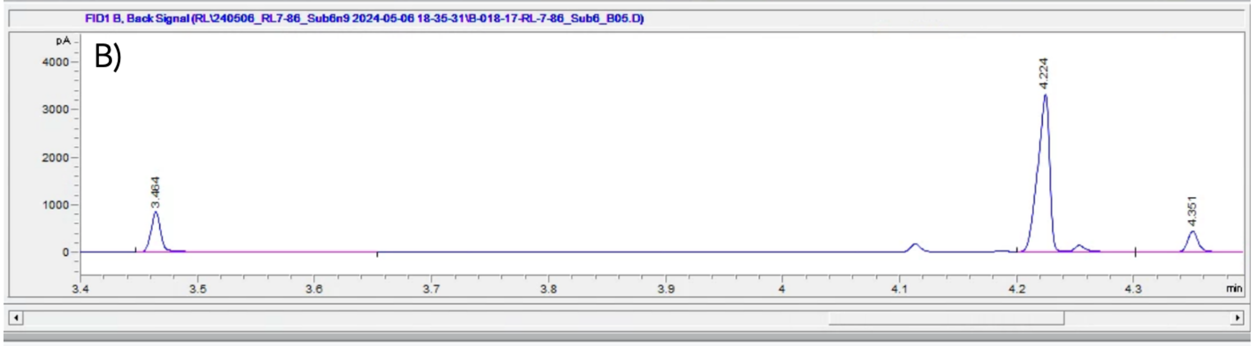


| # | Time | Type | Area | Height | Width | Area% | Symmetry |
|---|---|---|---|---|---|---|---|
| 1 | 3.464 | BV R | 503.9 | 812.9 | 9.5788E-3 | 15.585 | 0.784 |
| 2 | 4.224 | BV R | 2447.4 | 3309.6 | 0.0115 | 75.698 | 1.459 |
| 3 | 4.351 | VV R | 281.8 | 424.8 | 0.01 | 8.717 | 0.875 |

**Figure S62.** A) Representative GC-FID trace for reactions of ParLQ with **1g** and EDA. B) GC-FID trace for the top-performing variant, HPFAW, in reactions with **1g** and EDA.

*Substrate 1h - 1-(trifluoromethyl)-4-vinylbenzene*

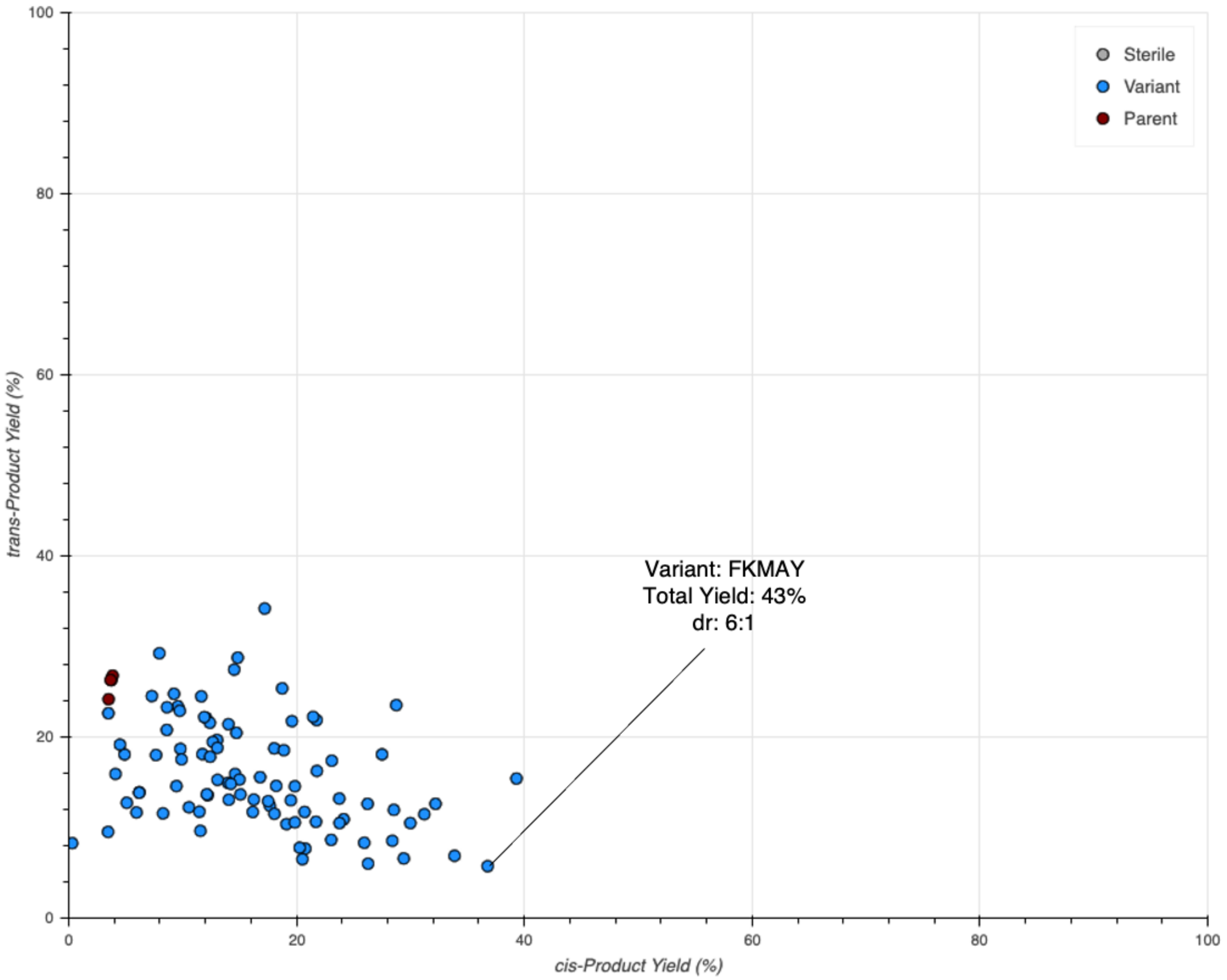


**Figure S63.** Activities of the second round of mutants predicted by ALDE with substrate **1h**. Observed yields for the formation of *cis*- and *trans*-**2h** are shown. The highest-performing variant, FKMAY, demonstrated a yield of 43% with a 6:1 selectivity for *cis*-**2h**.

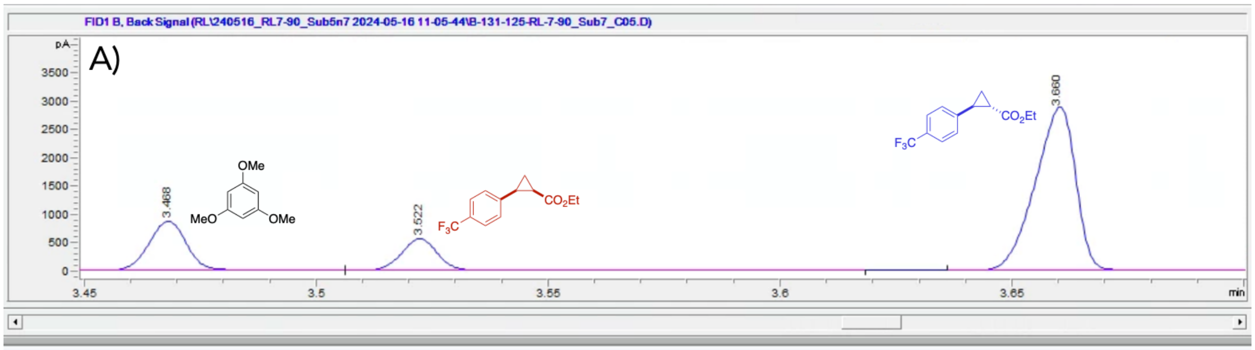


| # | Time | Type | Area | Height | Width | Area% | Symmetry |
|---|---|---|---|---|---|---|---|
| 1 | 3.468 | BV | 498.5 | 850.8 | 9.236E-3 | 19.342 | 0.912 |
| 2 | 3.522 | VV R | 309.9 | 534.3 | 9.9109E-3 | 12.023 | 0.906 |
| 3 | 3.66 | BV R | 1769 | 2817.9 | 9.6672E-3 | 68.635 | 1.18 |

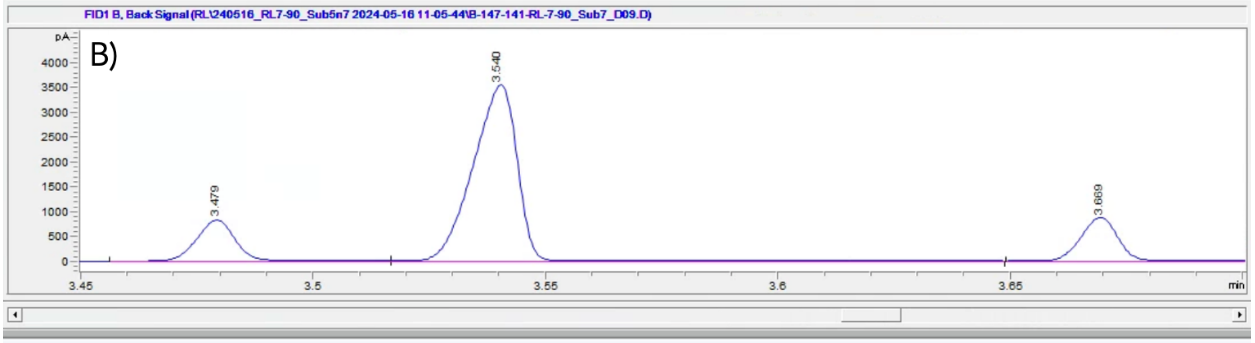


| # | Time | Type | Area | Height | Width | Area% | Symmetry |
|---|---|---|---|---|---|---|---|
| 1 | 3.479 | BV | 480.6 | 801.3 | 9.3973E-3 | 14.514 | 0.92 |
| 2 | 3.54 | VV R | 2331.9 | 3498.3 | 0.011 | 70.420 | 1.277 |
| 3 | 3.669 | BV R | 498.9 | 834.6 | 0.01 | 15.067 | 0.941 |

**Figure S64.** A) Representative GC-FID trace for reactions of ParLQ with **1h** and EDA. B) GC-FID trace for the top-performing variant, FKMAY, in reactions with **1h** and EDA.

*Substrate 1i – 2-Vinylnaphythelene*

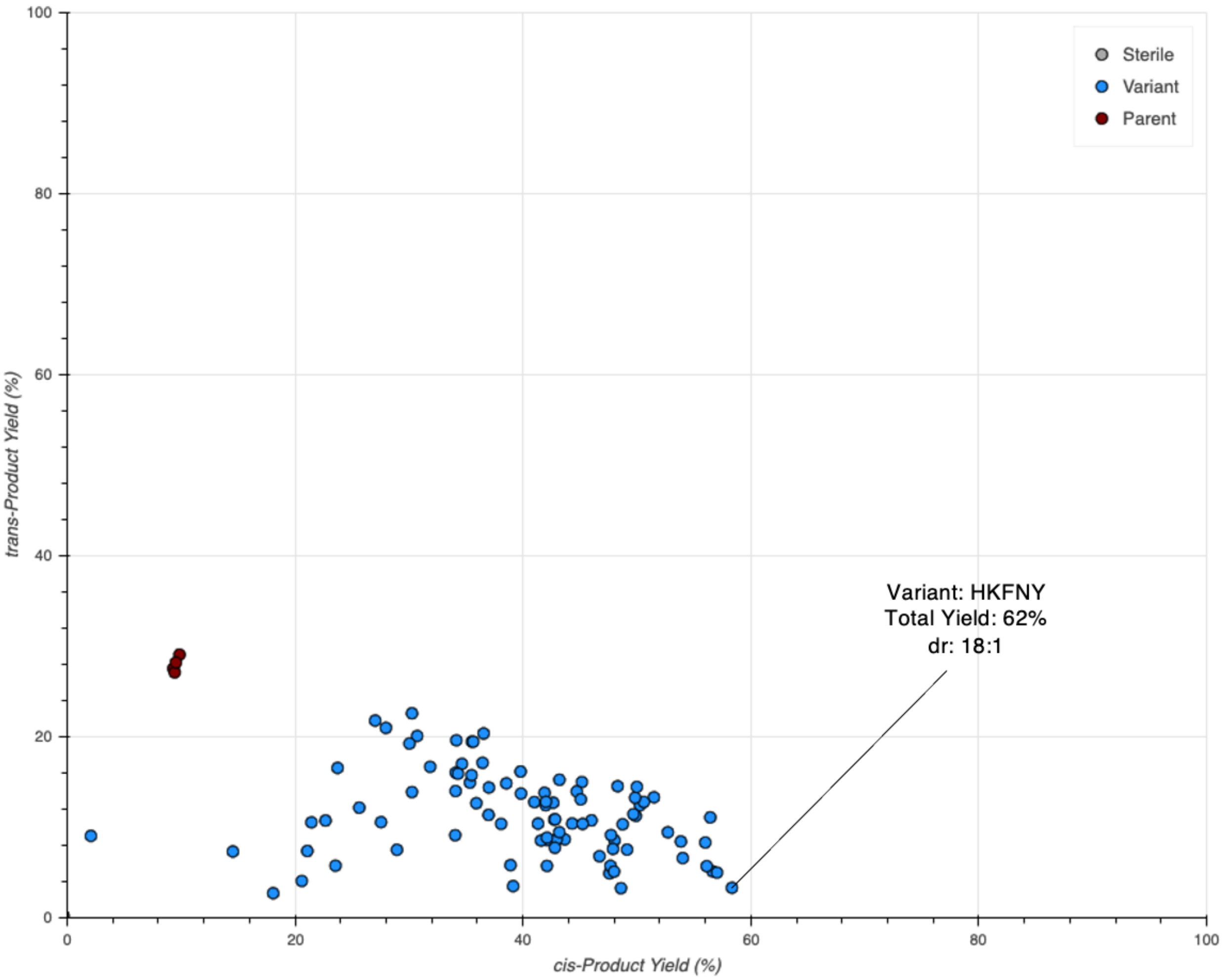


**Figure S65.** Activities of the second round of mutants predicted by ALDE with substrate **1i**. Observed yields for the formation of *cis*- and *trans*-**2i** are shown. The highest-performing variant, HKFNY, demonstrated a yield of 62% with an 18:1 selectivity for *cis*-**2i**.

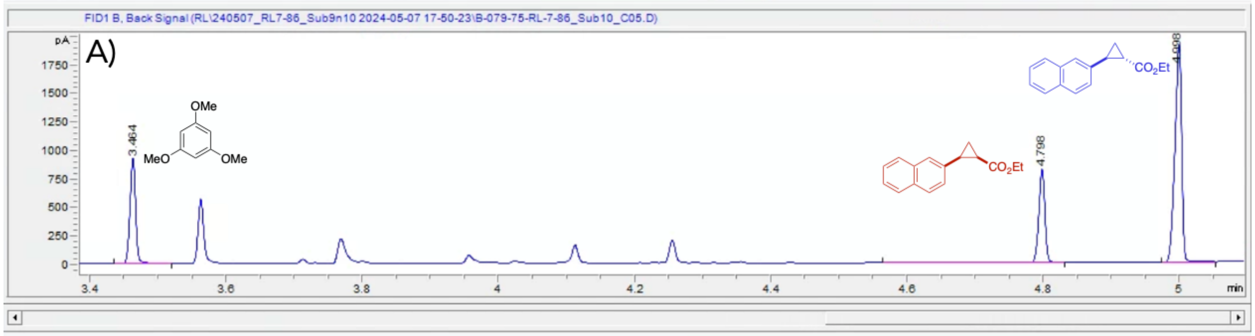


| # | Time | Type | Area | Height | Width | Area% | Symmetry |
|---|---|---|---|---|---|---|---|
| 1 | 3.464 | BB | 519.3 | 900.9 | 9.1252E-3 | 21.584 | 0.974 |
| 2 | 4.798 | VB R | 551.7 | 800.3 | 0.0106 | 22.931 | 1.569 |
| 3 | 4.998 | BB | 1334.9 | 1872.1 | 0.0107 | 55.485 | 1.27 |

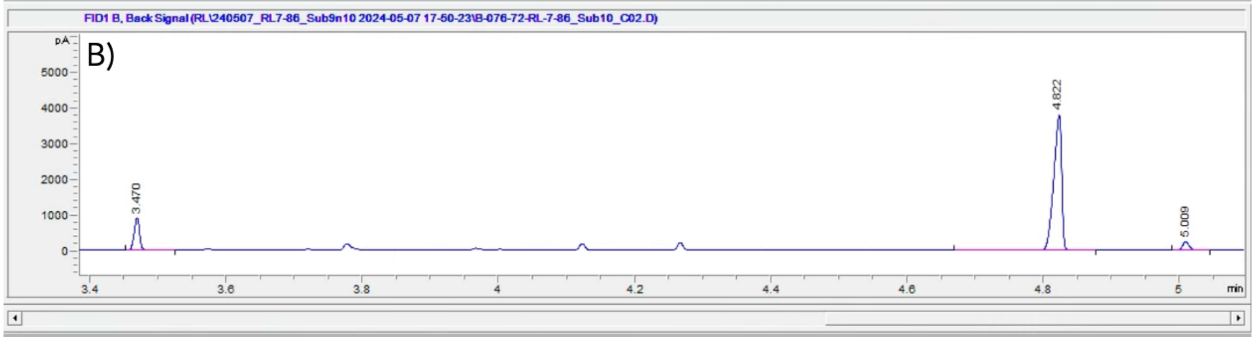


| # | Time | Type | Area | Height | Width | Area% | Symmetry |
|---|---|---|---|---|---|---|---|
| 1 | 3.47 | BB | 516.6 | 897.3 | 9.117E-3 | 13.677 | 0.968 |
| 2 | 4.822 | VB R | 3111.5 | 3776.4 | 0.0128 | 82.370 | 1.629 |
| 3 | 5.009 | BB | 149.3 | 227.5 | 0.0109 | 3.952 | 0.933 |

**Figure S66.** A) Representative GC-FID trace for reactions of ParLQ with **1i** and EDA. B) GC-FID trace for the top-performing variant, HKFNY, in reactions with **1i** and EDA.

### Chiral Traces for Determination of Enantiopurity

#### Determination of Enzyme Enantioselectivity

For reactions of **1a** with the libraries of ALDE-predicted variants, all samples were additionally screened for the enantiomeric ratio (er) of the *cis*- products. After analysis by achiral GC-FID samples were directly analyzed by chiral GC-FID using a chiral Agilent J&W CycloSil-B column.

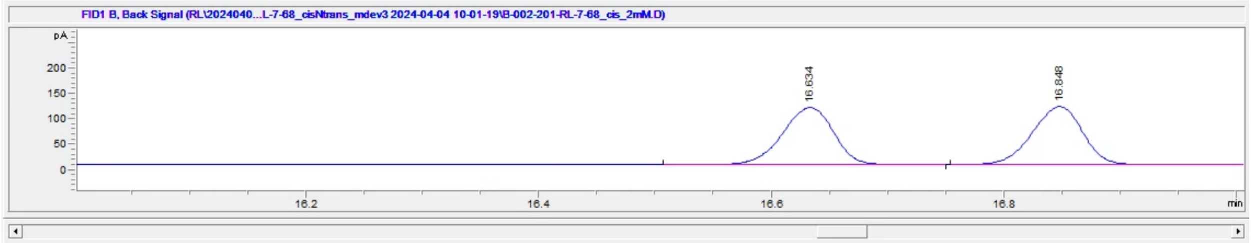


| # | Time | Type | Area | Height | Width | Area% | Symmetry |
|---|---|---|---|---|---|---|---|
| 1 | 16.634 | BB | 340 | 111.6 | 0.0485 | 49.877 | 1.166 |
| 2 | 16.848 | BB | 341.7 | 113.6 | 0.0471 | 50.123 | 1.146 |

**Figure S67.** Chiral GC trace of authentic standard for *cis*-**2a**

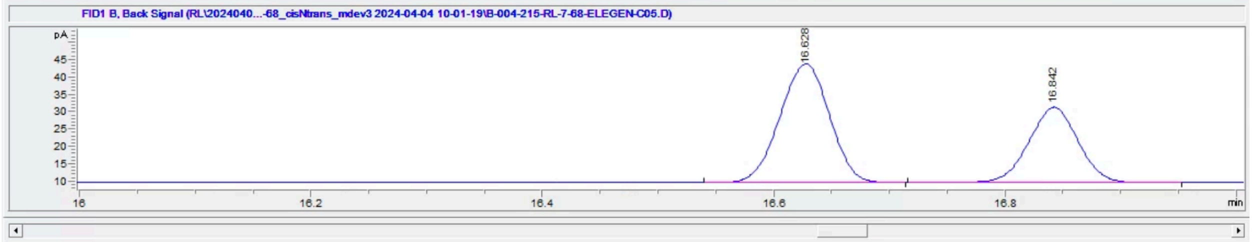


| # | Time | Type | Area | Height | Width | Area% | Symmetry |
|---|---|---|---|---|---|---|---|
| 1 | 16.628 | BB | 102.2 | 34.1 | 0.047 | 61.371 | 1.073 |
| 2 | 16.842 | BB | 64.4 | 21.7 | 0.0466 | 38.629 | 1.015 |

**Figure S68.** Chiral GC trace of products from reaction of **1a** with ParLQ

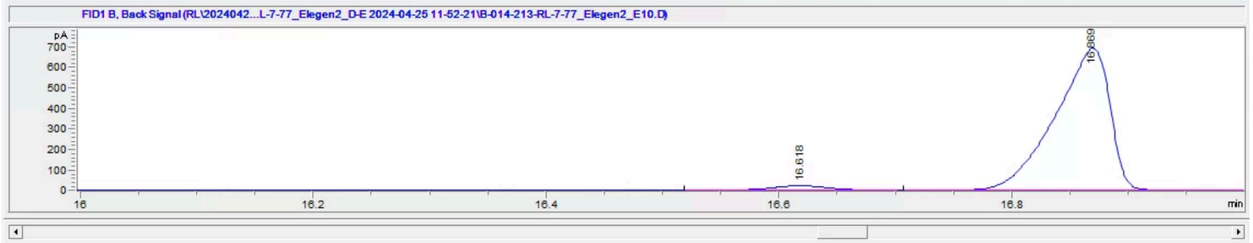


| # | Time | Type | Area | Height | Width | Area% | Symmetry |
|---|---|---|---|---|---|---|---|
| 1 | 16.618 | BB | 63.4 | 20.5 | 0.0491 | 2.750 | 0.997 |
| 2 | 16.869 | BB | 2240.9 | 688.6 | 0.047 | 97.250 | 2.095 |

**Figure S69.** Chiral GC trace of products from reaction of **1a** with MPFDY

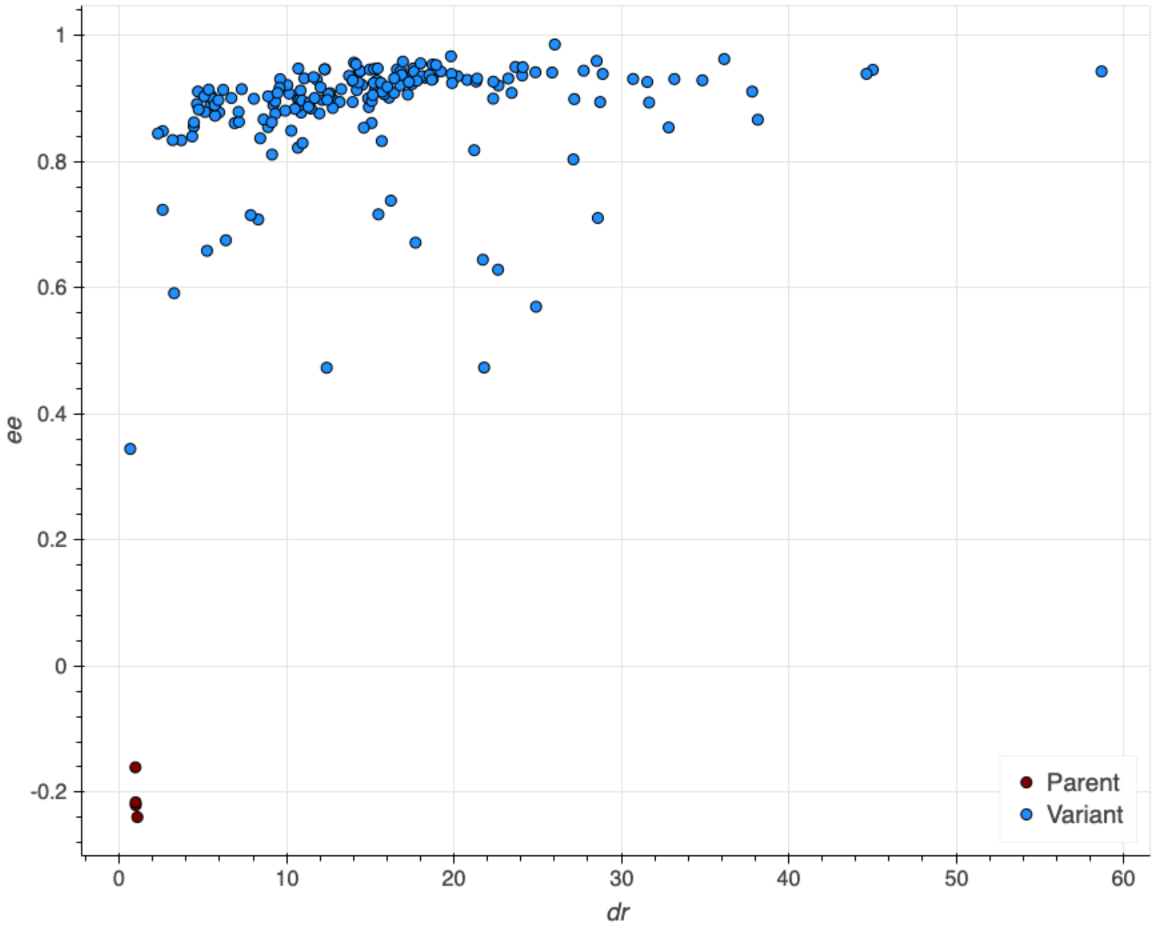


**Figure S70.** Enantioselectivity data for all ALDE predicted variants for the production of **2a**. ee values are plotted against dr values for each sample.

For all other substrates the enantioselectivity of the catalyst was only measured for the top-performing variant. All samples were measured using the same method as for the chiral separation of *cis*-**2a** by GC.

### [1]H NMR Spectra of Authentic Standards

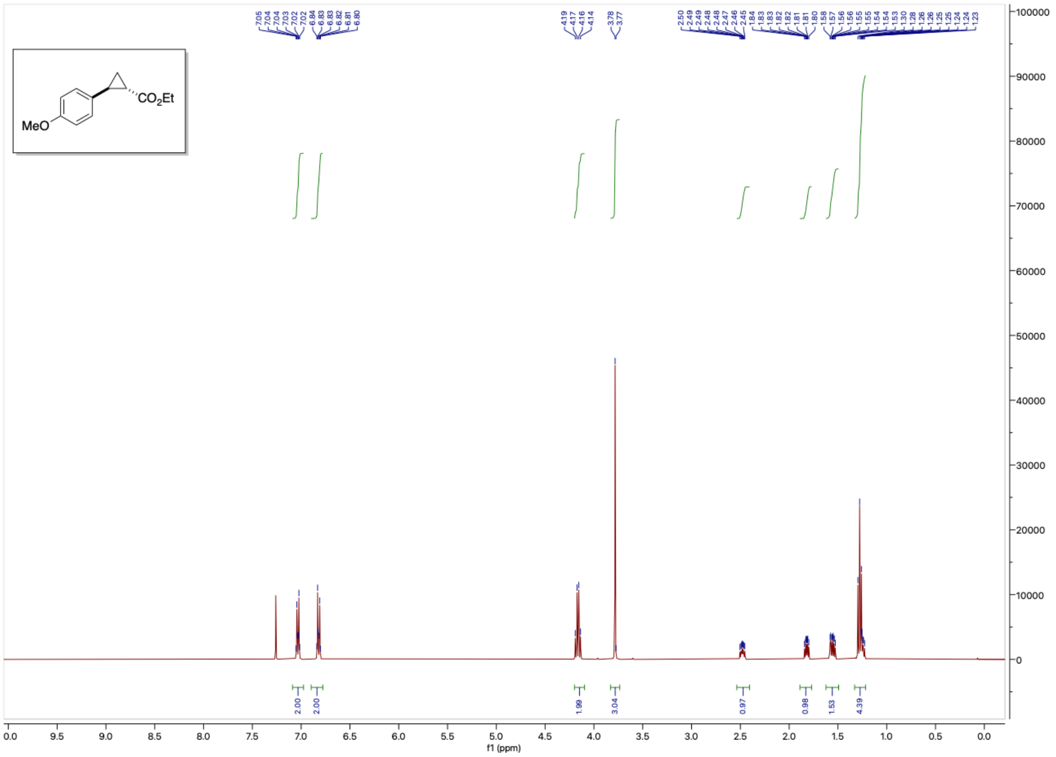

**Figure S71.** $^1$H NMR spectrum of racemic *trans*-**2a**.

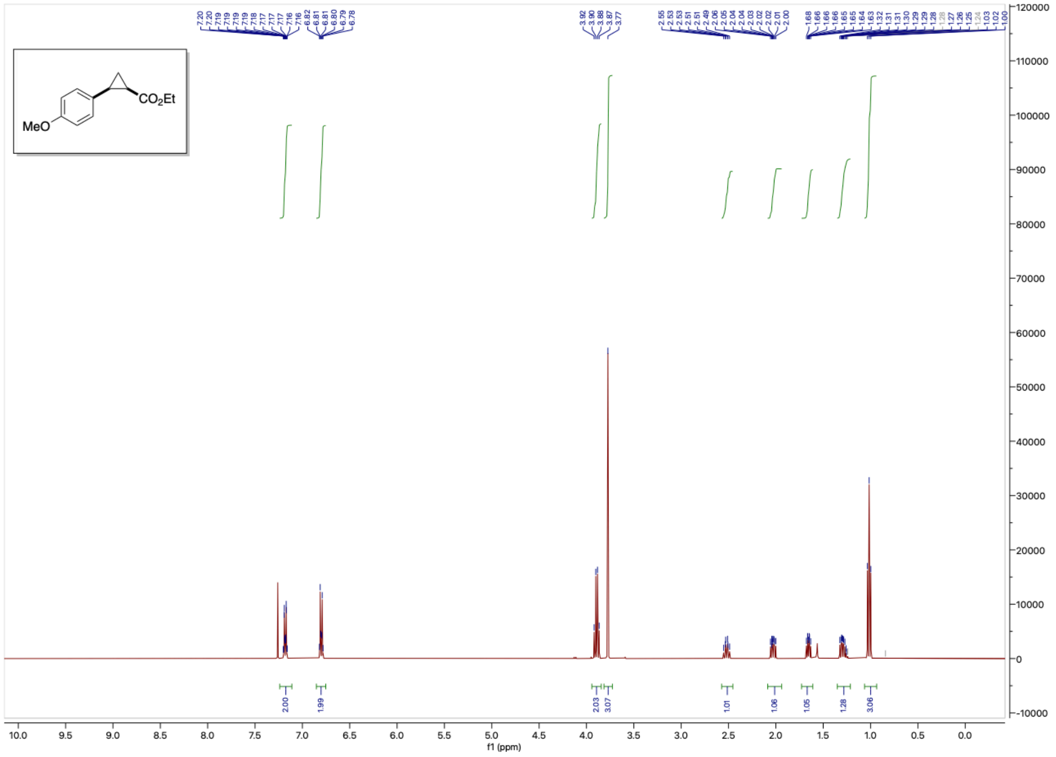

**Figure S72.** $^{1}H$ NMR spectrum of racemic *cis*-**2a**.

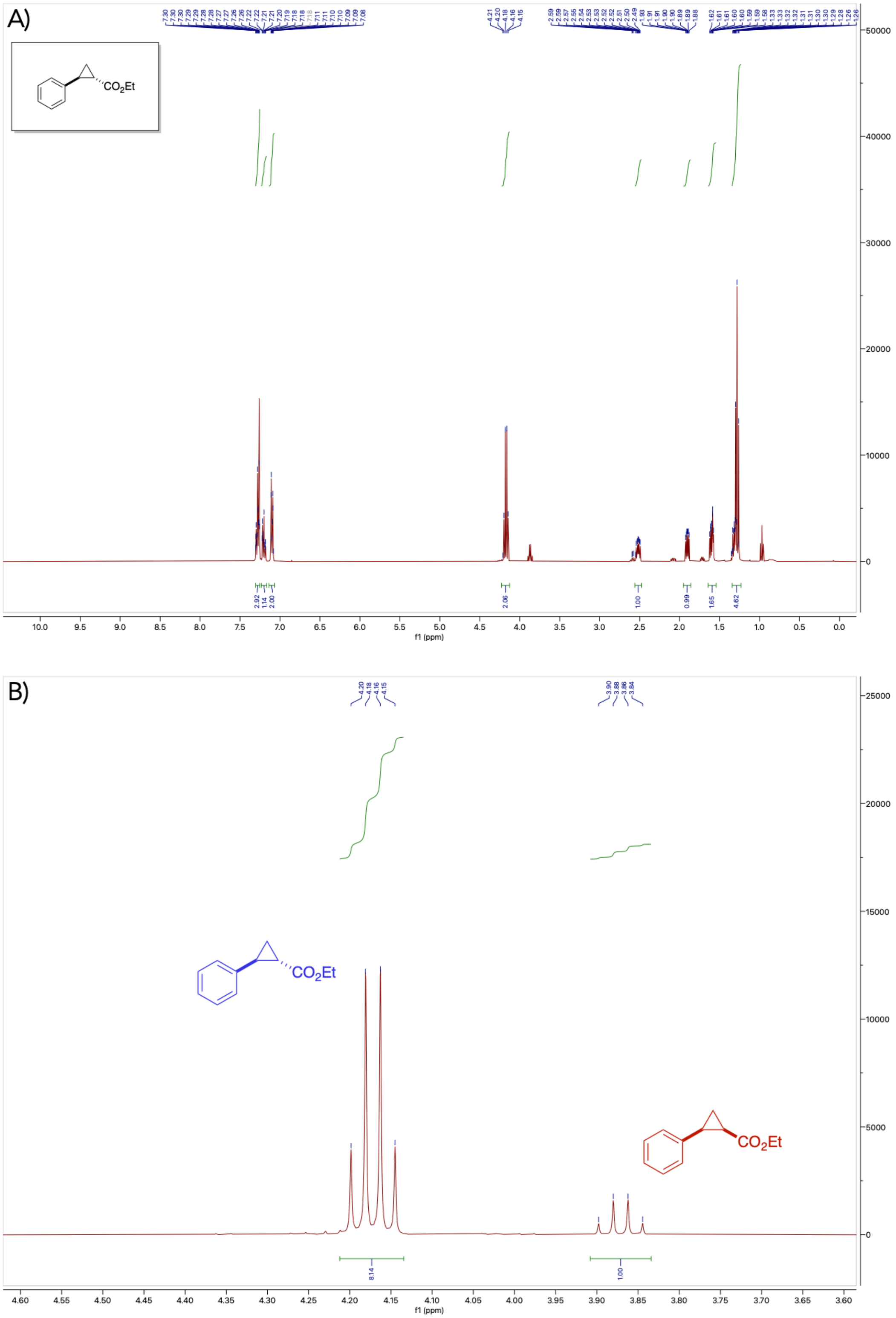


**Figure S73.** A) [1]H NMR spectrum of a 8:1 racemic *trans*-**2b**/*cis*-**2b**. B) Integrations of product ester methylene protons for *trans*-**2b** and *cis*-**2b** to determine the isolated molar ratio.

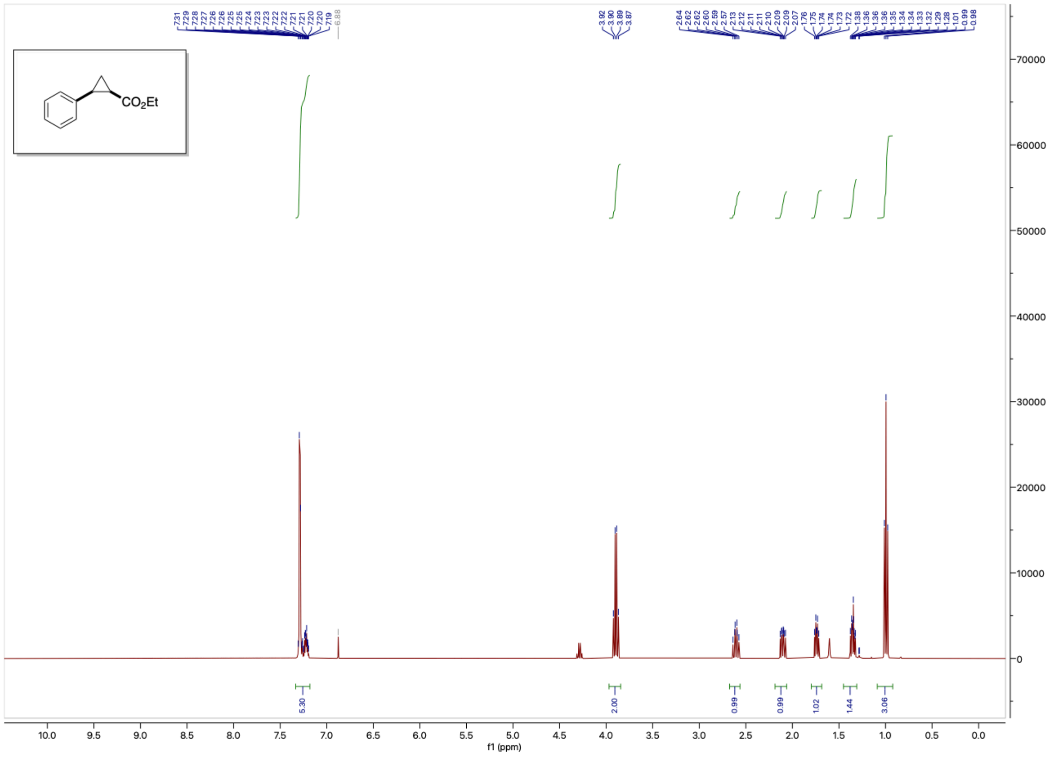

**Figure S74.** $^{1}$H NMR spectrum of racemic *cis*-**2b.** A trace impurity of diethyl fumarate can be observed.

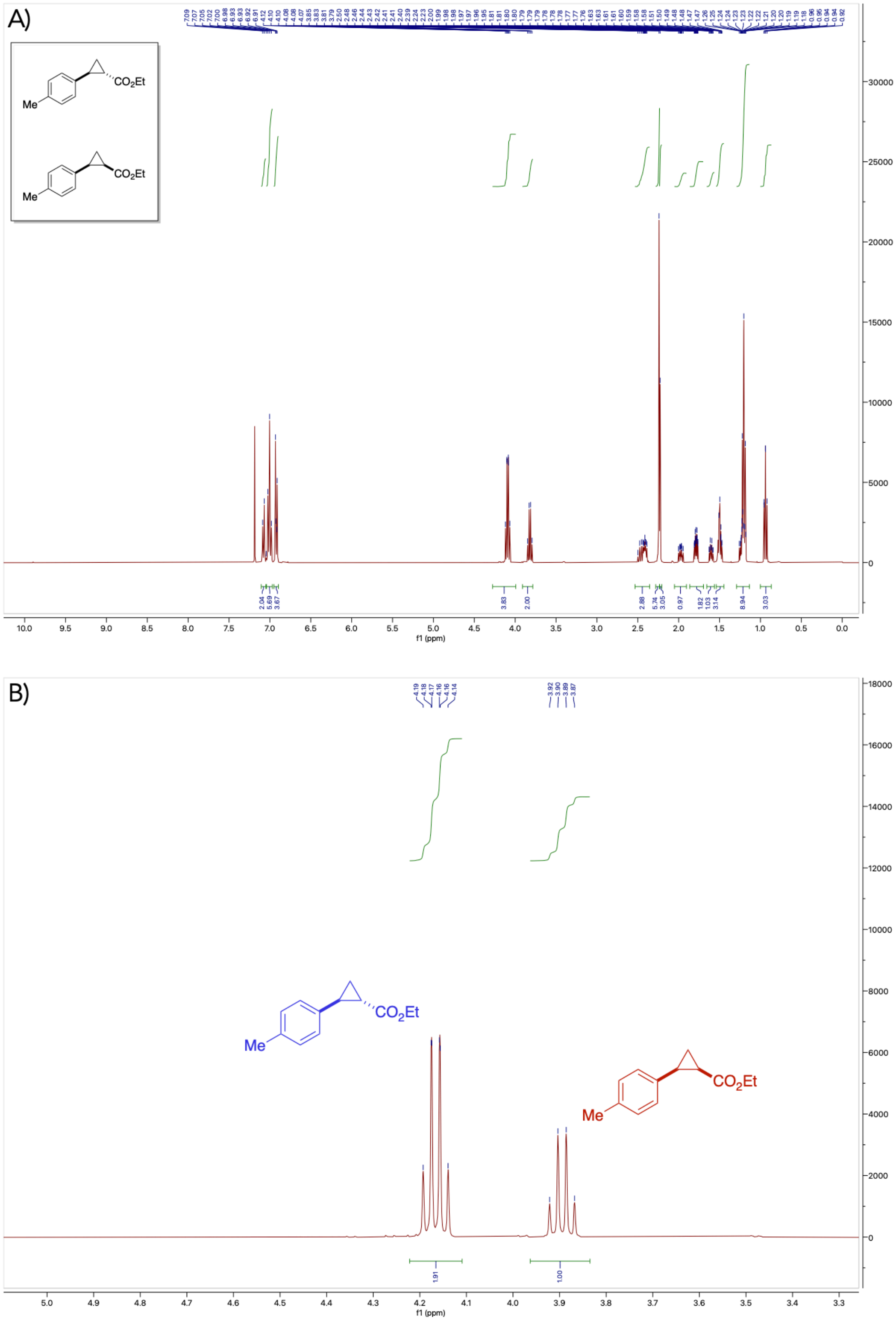


**Figure S75.** A) [1]H NMR spectrum of a 1.91:1 racemic *trans*-**2c**/*cis*-**2c**. B) Integrations of product ester methylene protons for *trans*-**2c** and *cis*-**2c** to determine the isolated molar ratio.

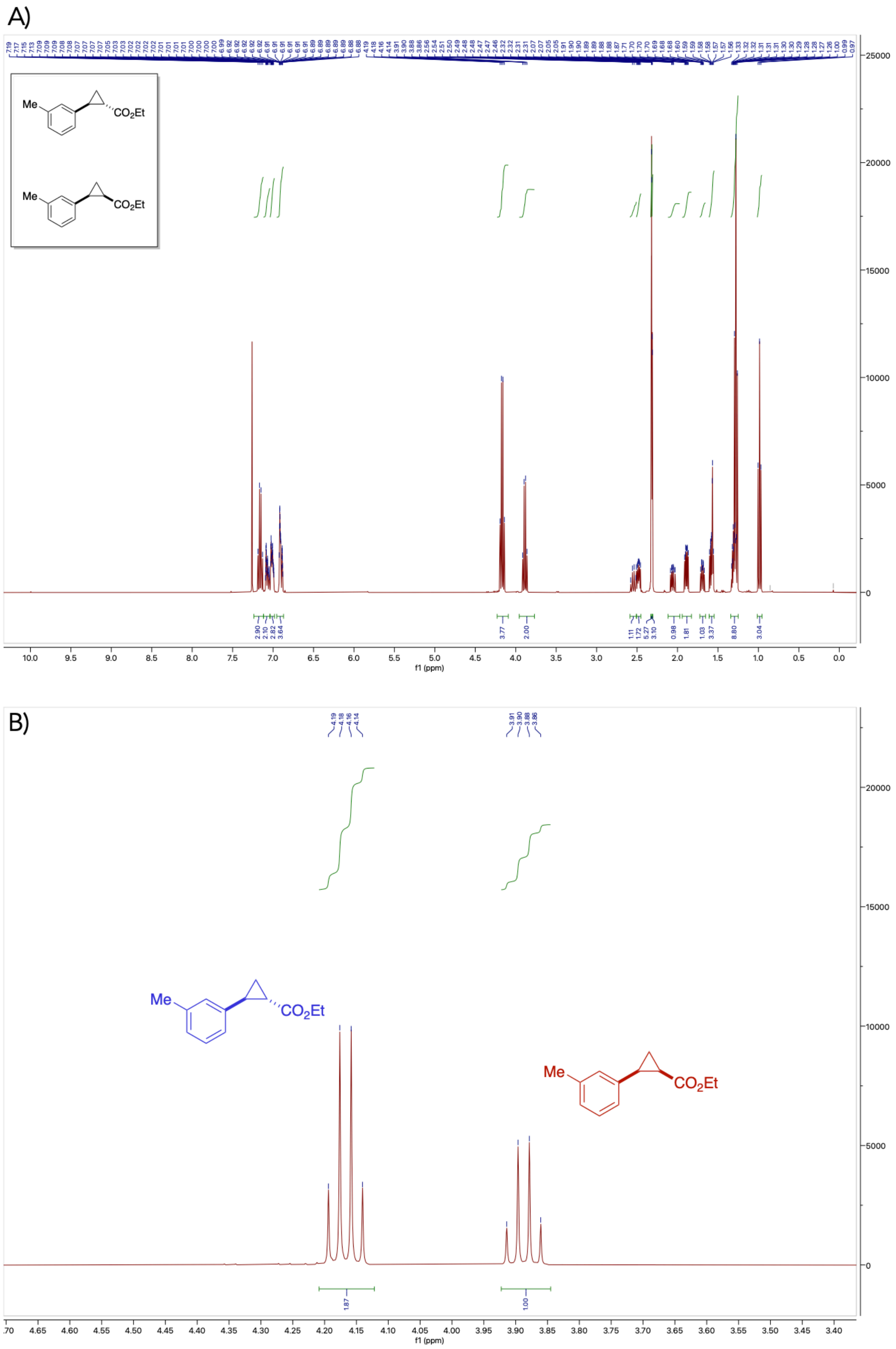


**Figure S76.** A) [1]H NMR spectrum of a 1.87:1 racemic *trans*-**2d**/*cis*-**2d**. B) Integrations of product ester methylene protons for *trans*-**2d** and *cis*-**2d** to determine the isolated molar ratio.

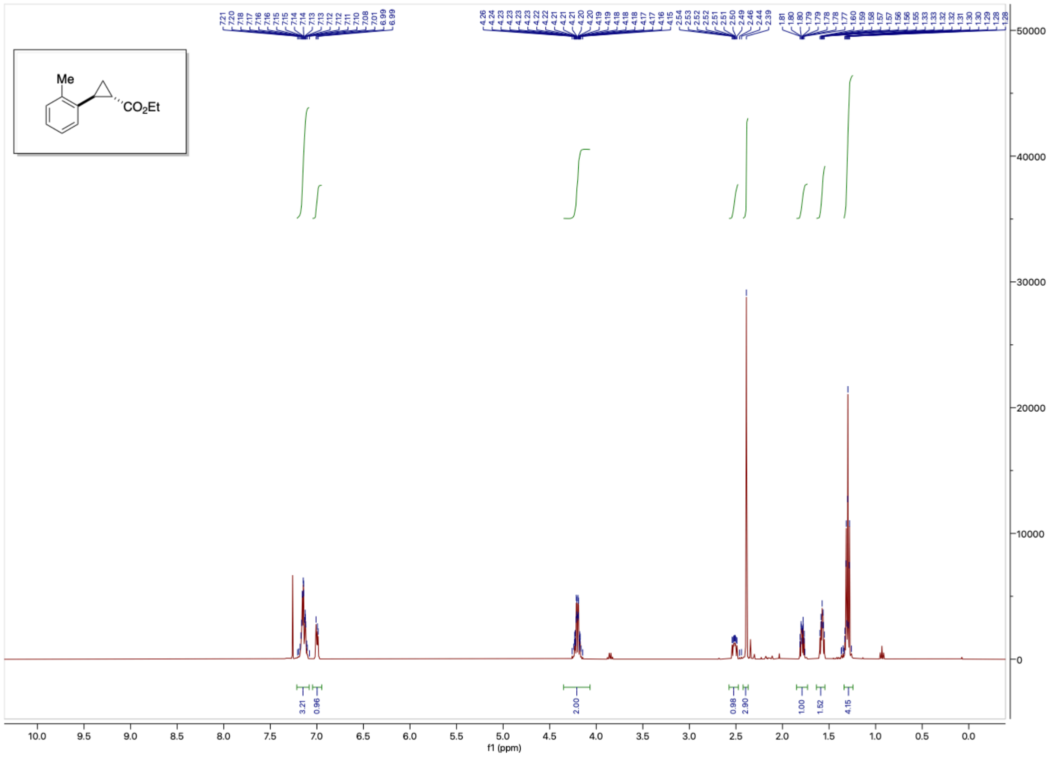

[1]H NMR spectrum of racemic *trans*-**2e**.

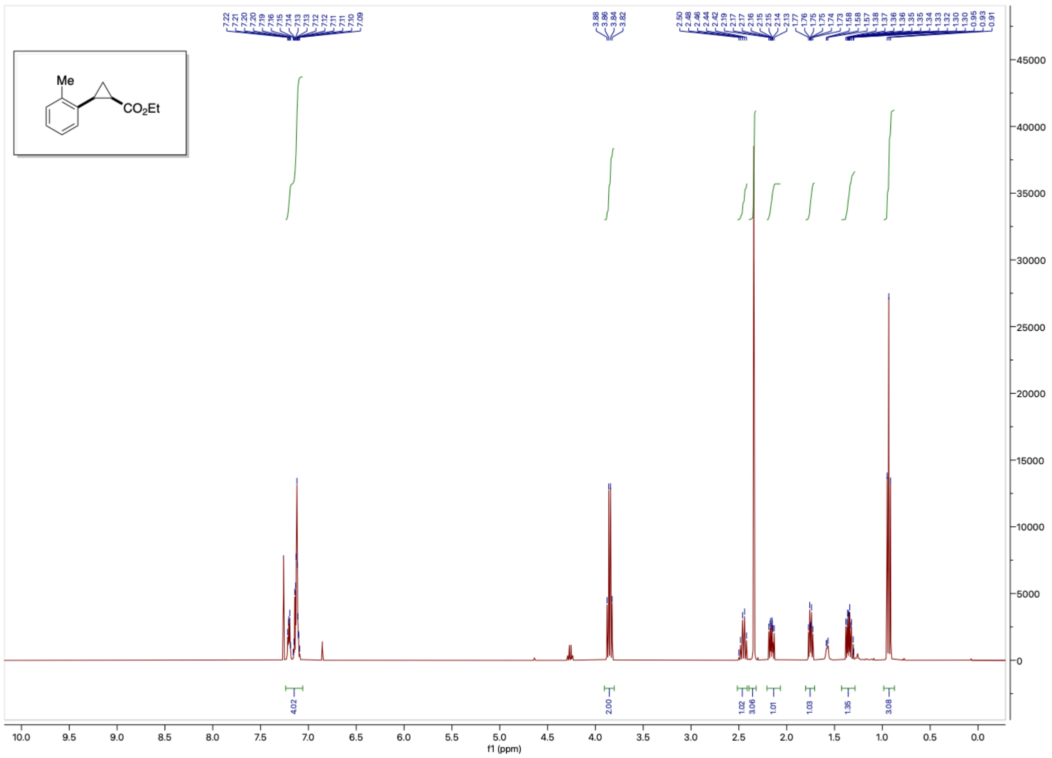

**Figure S77.** [1]H NMR spectrum of racemic *cis*-**2e**. A trace impurity of diethyl fumarate can be observed.

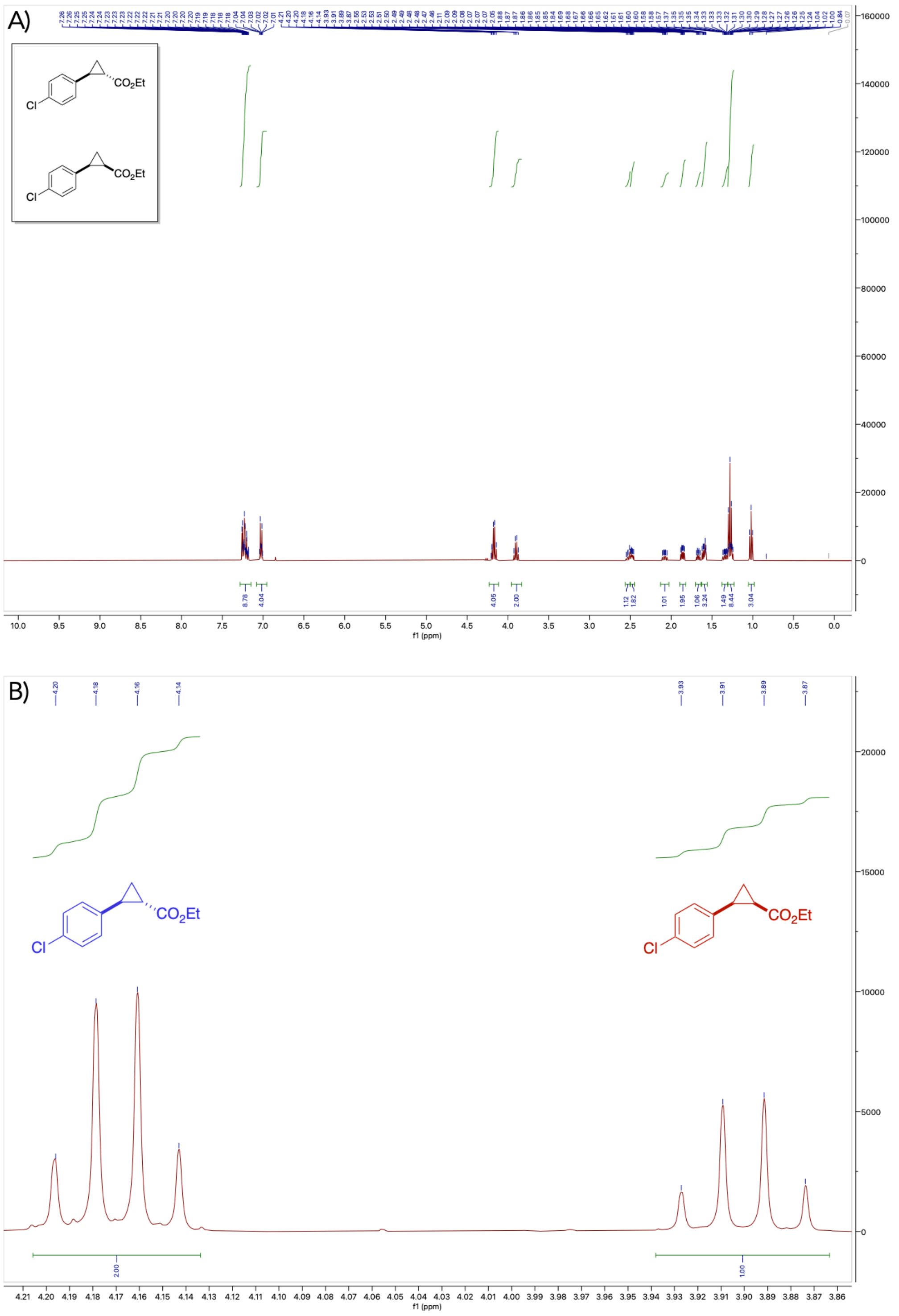


**Figure S78.** A) [1]H NMR spectrum of a 2:1 racemic *trans*-**2f**/*cis*-**2f**. A trace impurity of diethyl fumarate can be observed. B) Integrations of product ester methylene protons for *trans*-**2f** and *cis*-**2f** to determine the isolated molar ratio.

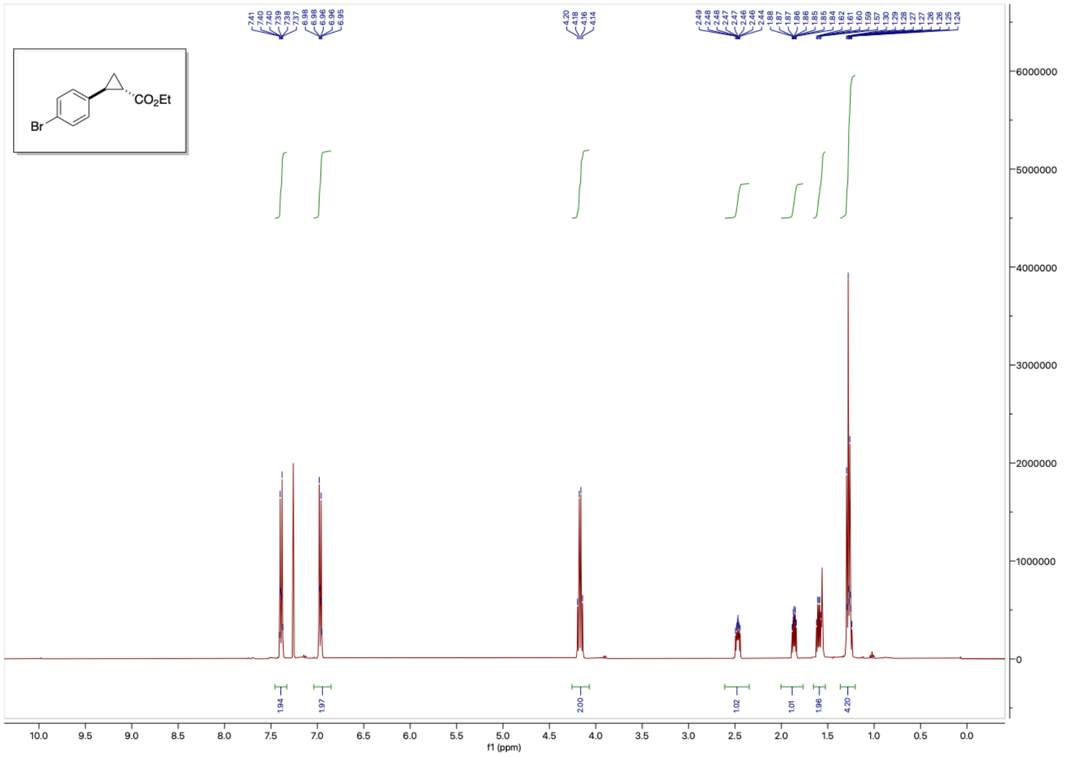

**Figure S91.** [1]H NMR spectrum of racemic *trans*-**2g**.

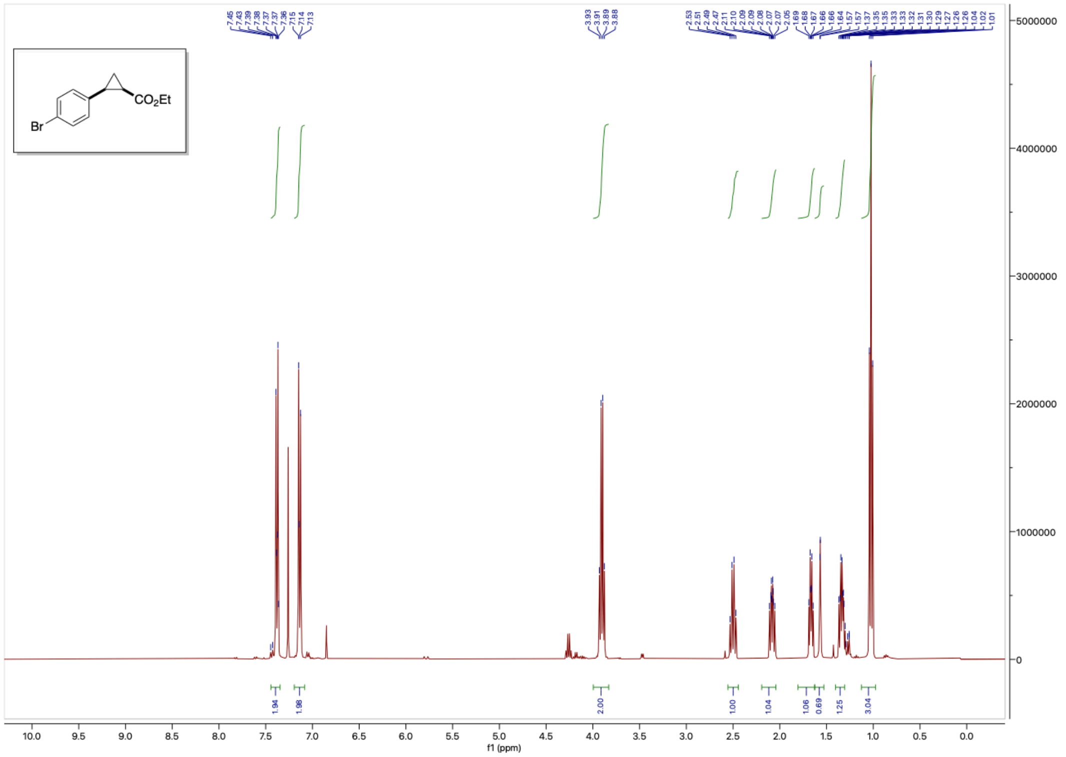

**Figure S79.** [1]H NMR spectrum of racemic *cis*-**2g**. A trace impurity of diethyl fumarate can be observed.

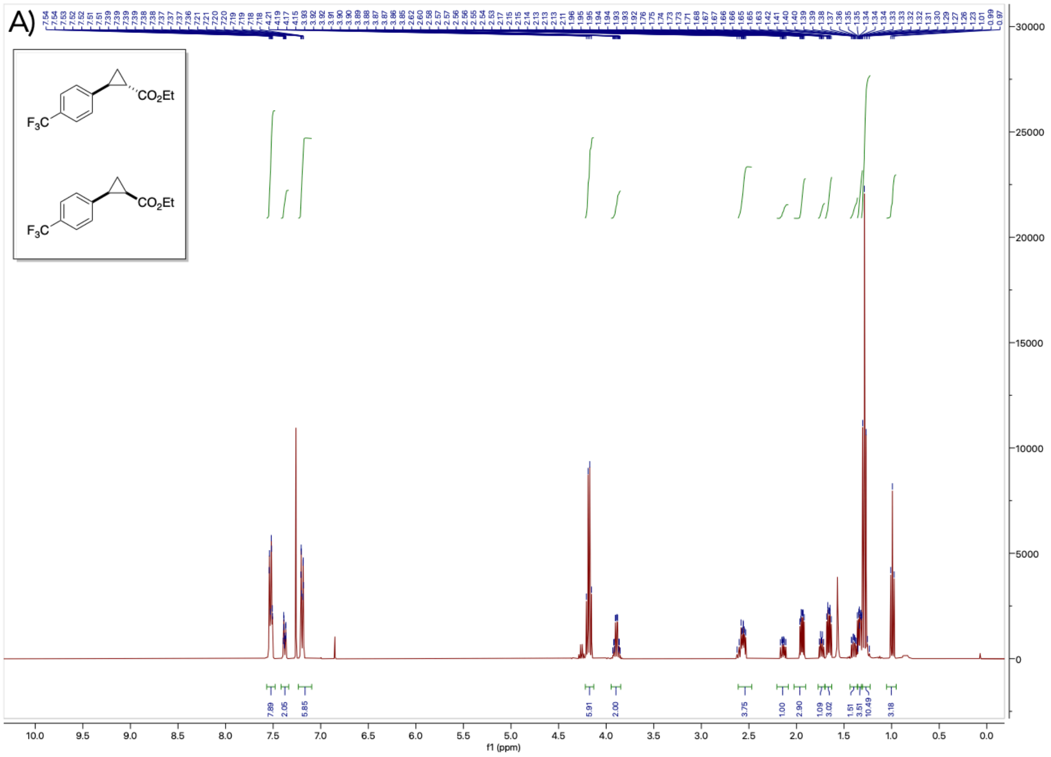

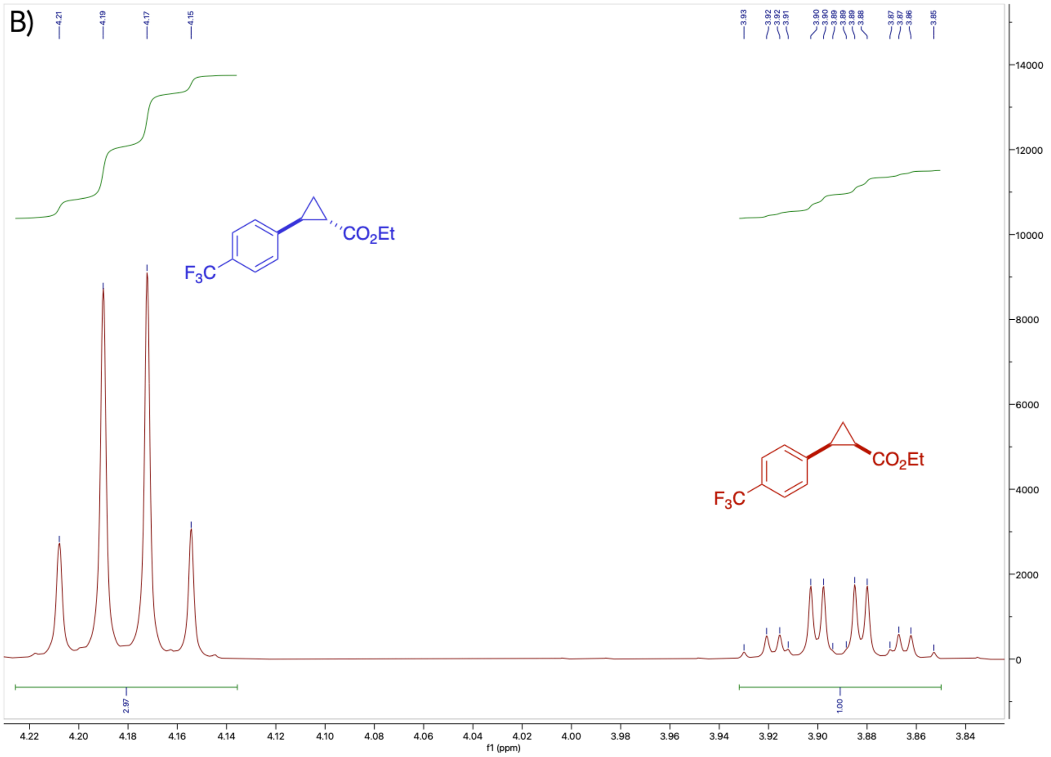

**Figure S80.** A) $^{1}$H NMR spectrum of a 2.97:1 racemic *trans*-**2h**/*cis*-**2h**. A trace impurity of diethyl fumarate can be observed. B) Integrations of product ester methylene protons for *trans*-**2h** and *cis*-**2h** to determine the isolated molar ratio.

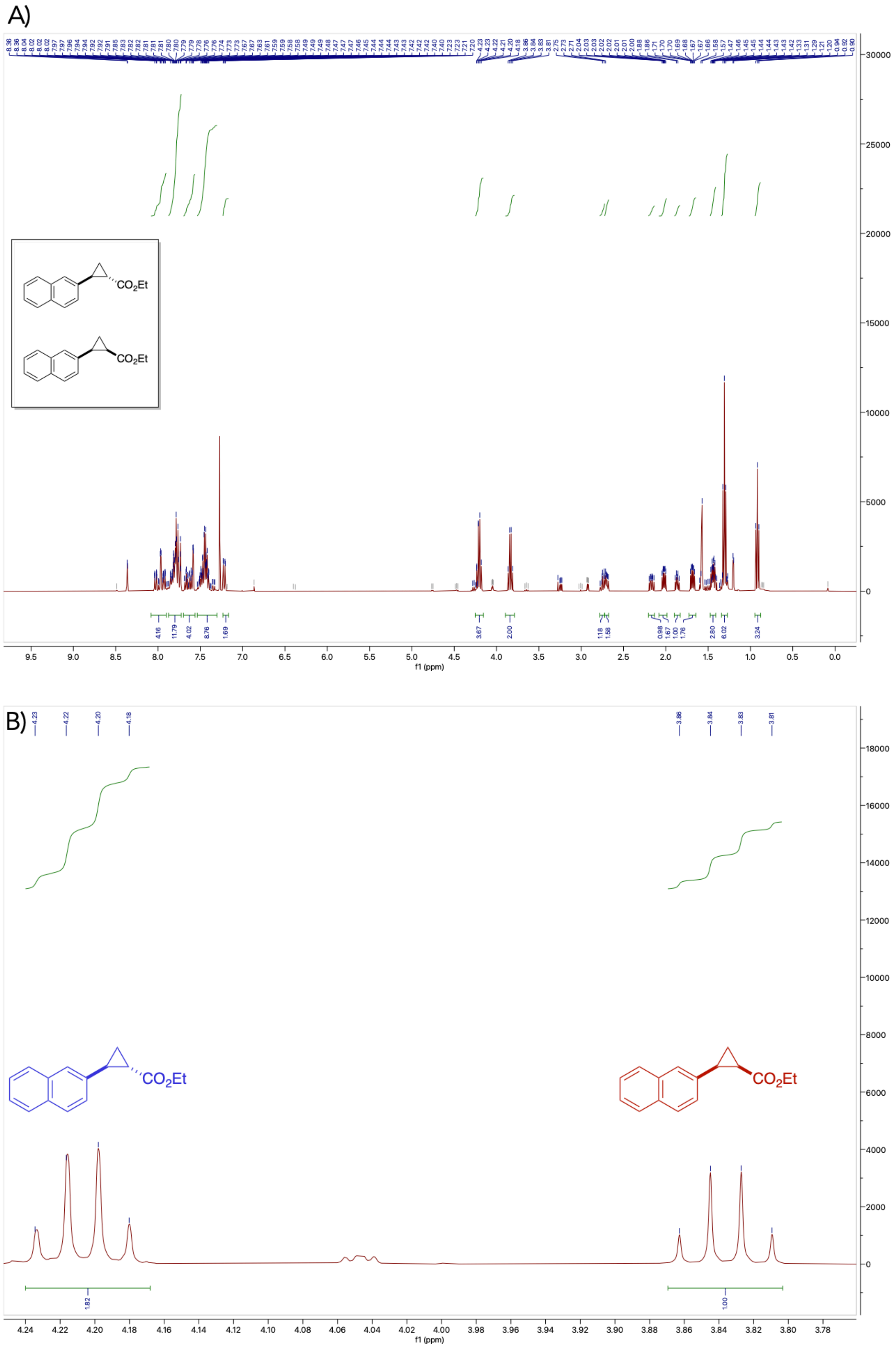


**Figure S81.** A) [1]H NMR spectrum of a 1.82:1 racemic *trans*-**2i**/*cis*-**2i**. Several trace impurities are observed. B) Integrations of product ester methylene protons for *trans*-**2i** and *cis*-**2i** to determine the isolated molar ratio.
